# Maturation-associated ERG3 and timed regulation of circuit stability

**DOI:** 10.1101/2024.09.22.614315

**Authors:** Hiroyasu Watanabe

## Abstract

Neural maturation remodels excitability, but how this supports stable computation remains unclear. In a layered network, unreliable transmission destabilised learning when forward and update perturbations differed. This motivated a developmental screen for conductances that could support reliable responses. Mouse *Kcnh7* (human *KCNH7*), encoding Kᵥ11.3 (ERG3), rose late postnatally. Adult single-cell data showed expression in excitatory and inhibitory populations. In a Purkinje-cell model, ERG3 raised tonic-firing threshold and broadened pre-tonic responses. ERG3-inspired adaptation prolonged effective learning but reduced associative recall at low sampling temperature. In recurrent models, ERG3 shaped history-dependent firing, and prediction depended on modulation timing and perturbation type. Clustered interruptions of receptor modulation increased departures from the maintained state in a wake-maintenance model, whereas cue-timed loss delayed value updating during sustained activity. Together with receptor expression and mouse orexin data, these findings motivate testing whether disrupted ERG3 modulation contributes to fragmented wakefulness or impaired adaptation to changed rules.

## Introduction

Artificial neural networks emerged from simplified abstractions of nervous systems and have shown how learning, memory and prediction can arise from interactions among elementary units. Such abstractions make it possible to isolate computational operations, including how a network assigns credit for an error.^1^ In recurrent networks, a readout can also use information about recent inputs retained in population activity.^2^ In biological circuits, the neurons that carry out those operations also change as the brain matures. Developmental and molecular studies can identify the channels and signalling pathways that change, while network models can identify the computations that become vulnerable. The challenge is to explain how a change in an identified cellular mechanism alters a computation and, in turn, the function of a circuit.

Postnatal maturation can improve the speed and reliability of synaptic transmission, as shown at the calyx of Held.^3^ The computational benefit of greater fidelity depends on what a circuit must preserve. In a layered learning system, an update must account for the effective transmission that produced the response. In a recurrent-network model, the effect of a fluctuation depends on how it changes the population activity used by the readout.^2^ This raises the question of which cellular mechanisms allow a circuit to retain a learned function through ordinary variability and which forms of disruption exceed that capacity.

Intrinsic conductances offer a way to address this question. Expression, voltage dependence and gating kinetics can be investigated separately, whereas the phenotype after constitutive gene loss can reflect both the immediate effect of losing a current and the circuit’s adaptation to that loss. Channels that operate near firing threshold are particularly informative because their gating can carry the effects of preceding activity into the response to a subsequent input. One candidate is the voltage-gated potassium channel Kᵥ11.3, also called ERG3, encoded by *KCNH7* in humans and *Kcnh7* in mice. In Purkinje cells, native ERG current contributes to the control of repetitive firing, and metabotropic glutamate receptor 1 activation changes both its amplitude and its voltage dependence.^4^ Mouse studies also implicate ERG3 in Purkinje-cell excitability^5^ and hippocampal seizure susceptibility.^6^ These findings motivate asking how the channel’s effects near threshold influence the activity available for learning and readout.

Orexin signalling provides a biological reason to ask how the timing of receptor input contributes to circuit function. Restoring orexin receptor 2 in the posterior hypothalamus improves wake maintenance in receptor-deficient mice^7^, and orexin-neuron activity varies around transitions from non-REM to REM sleep.^8^ In prefrontal cortical neurons, bursts can recruit a prolonged depolarisation through metabotropic glutamate receptor 5.^9^ These observations raise the question of how a circuit integrates receptor signals that recur at intervals, and whether modulation during ongoing activity serves a different function from modulation recruited around a task cue. The sensitivity of native Purkinje-cell ERG current to mGluR1 activation^4^ provides a basis for asking whether ERG3 participates in either process.

Here, I use a layered learning model to identify when unreliable transmission destabilises an acquired solution, then use developmental expression data to nominate *Kcnh7* as a candidate for regulating neuronal responses. In conductance-based models, ERG3 changes near-threshold firing and how recent activity shapes subsequent population responses. Its effects on prediction depend on when gating is altered and how the circuit is perturbed. Finally, models of receptor-linked modulation distinguish two consequences of losing that signal: a circuit can leave a maintained activity state, or it can remain active while taking longer to revise learned values.

## Results

### Low-fidelity transmission reveals delayed collapse under forward–update mismatch

Because postnatal maturation can increase the reliability and temporal precision of neural transmission, I asked how continuous learning behaves when state transmission remains unreliable. In seed-matched MNIST classifiers, I examined a mapping-level mismatch in which multiplicative perturbations were applied to the affine weights used during forward computation, whereas layerwise credit was propagated through the corresponding unperturbed weights. The loss was therefore generated by the perturbed forward output, but the update was assigned through a different effective mapping.

Despite this persistent mismatch, the classifiers acquired high clean-evaluation accuracy before performance abruptly fell to near-chance levels, accompanied by rapid and sustained expansion of the weight norm (Fig. 1a,b). Because forward perturbations were disabled during evaluation, the decline reflected destabilization of an acquired clean mapping rather than transient corruption of the test computation. The mapping-mismatch condition also used a lower learning rate. Its trajectory therefore illustrates the effect of that combination of settings.

**Figure 1.**
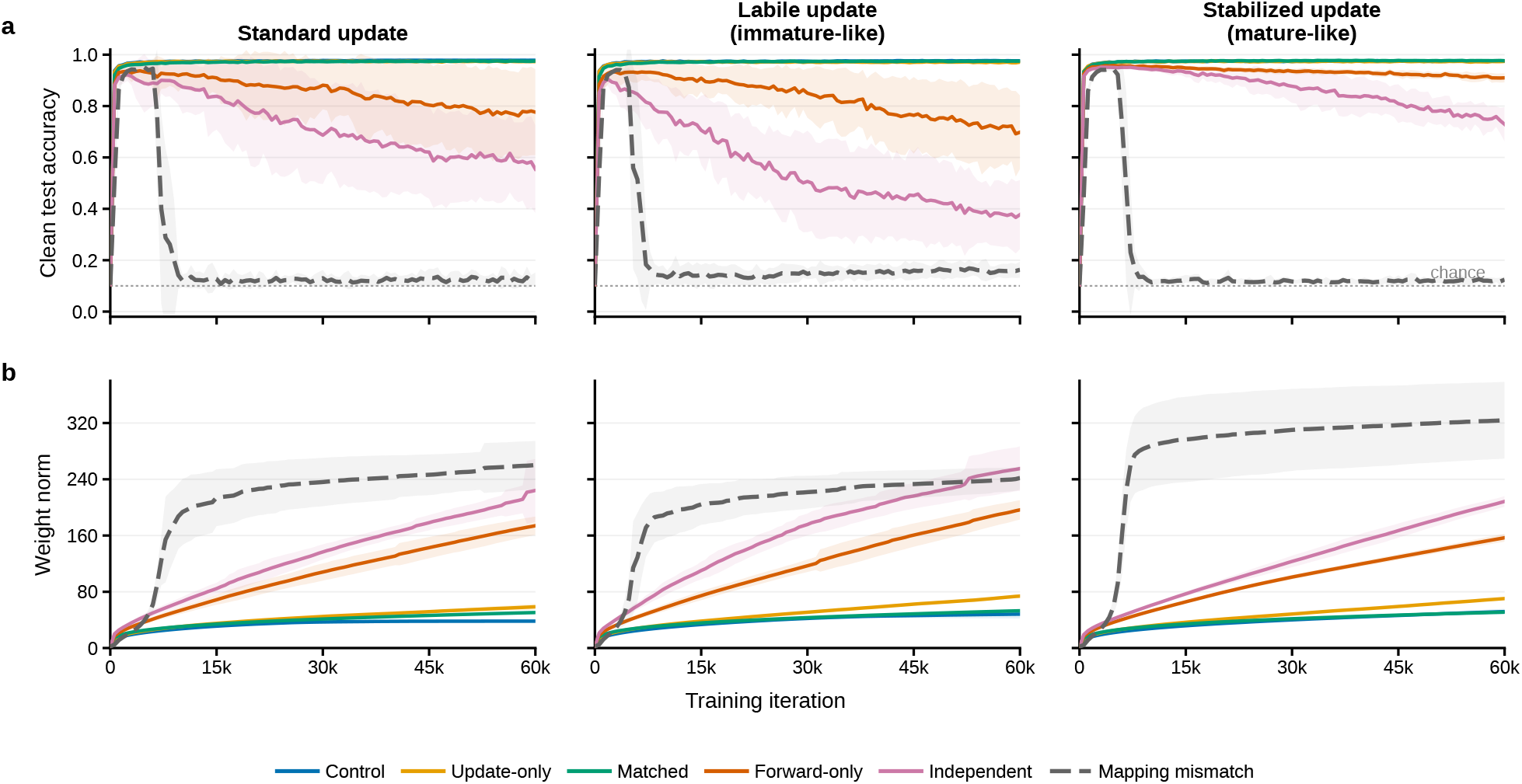
Low-fidelity transmission reveals delayed collapse under forward–update mismatch. a, Clean-evaluation accuracy under Standard update, Labile update (immature-like) and Stabilized update (mature-like) regimes. The five conditions using the effective forward weights for upstream error propagation were Control, Update-only (local gradient perturbed), Matched (local gradient reused the forward mask), Forward-only (local gradient omitted the forward mask) and Independent (separate forward and local-gradient masks). Mapping mismatch, shown as a dashed dark-grey curve, applied multiplicative perturbations to affine weights during forward computation, whereas layerwise credit was propagated through the corresponding unperturbed weights. The dotted horizontal line indicates chance accuracy of 0.1. b, Frobenius norm of model weights for the same runs. Curves and shaded regions show mean ± s.d. across ten seed-matched runs (n = 10). Clean-evaluation accuracy was measured with forward perturbations disabled. The mapping-mismatch condition used a learning rate of 0.1. The other conditions used 0.5. Update-regime labels denote operational computational settings rather than quantitative developmental fits.

To determine whether degraded transmission alone was sufficient to produce this failure, I next varied whether each local weight update incorporated its forward transmission mask, while propagating errors through the effective forward weights. Five conditions were compared under standard updating, a labile immature-like update regime and a stabilized mature-like update regime. These labels describe relative computational settings rather than quantitative models of developmental age.

Control, update-only perturbation and matched forward–update perturbation acquired and maintained high clean-evaluation accuracy. By contrast, forward-only and independently resampled perturbations produced progressive loss of accuracy after acquisition (Fig. 1a). This deterioration was present under standard updating, was strongest in the labile regime and was attenuated in the stabilized regime. The mapping-mismatch condition produced an abrupt accuracy collapse in all three update regimes. Weight trajectories showed the same overall separation (Fig. 1b).

Weight norms remained comparatively bounded in the Control, update-only and matched conditions, whereas forward-only and independent conditions showed sustained growth. This growth was greatest in the labile regime and reduced in the stabilized regime. Mapping mismatch instead produced an early, rapid expansion of the weight norm across the update regimes.

Together, these results show that degraded transmission alone was insufficient to account for learning instability. Among the five conditions with a common learning rate, acquired performance declined when the local weight gradient omitted or independently resampled the forward perturbation. Stabilized update dynamics attenuated this progressive failure. Mapping-level mismatch revealed a more abrupt loss of an acquired solution, distinguishing the ability to learn a task initially from the ability to preserve that learning during continued updating.

### Developmental potassium-channel screening identifies *Kcnh7* as a maturation-associated candidate

I next sought a molecular entry point through which postnatal changes in intrinsic conductance could be followed from gene expression to cellular and network function. Because potassium channels are major determinants of firing threshold, repetitive activity and temporal response properties, I screened the Allen Developing Mouse Brain Atlas for potassium-channel genes that increased between P4 and P56 across the telencephalic vesicle, midbrain and medullary hindbrain.^10,11^ This screen identified *Kcnh7*, encoding the Kv11.3 potassium channel commonly termed ERG3, as a candidate with higher expression energy at P56 than at P28 in all three matched structures (Fig. 2a).

**Figure 2.**
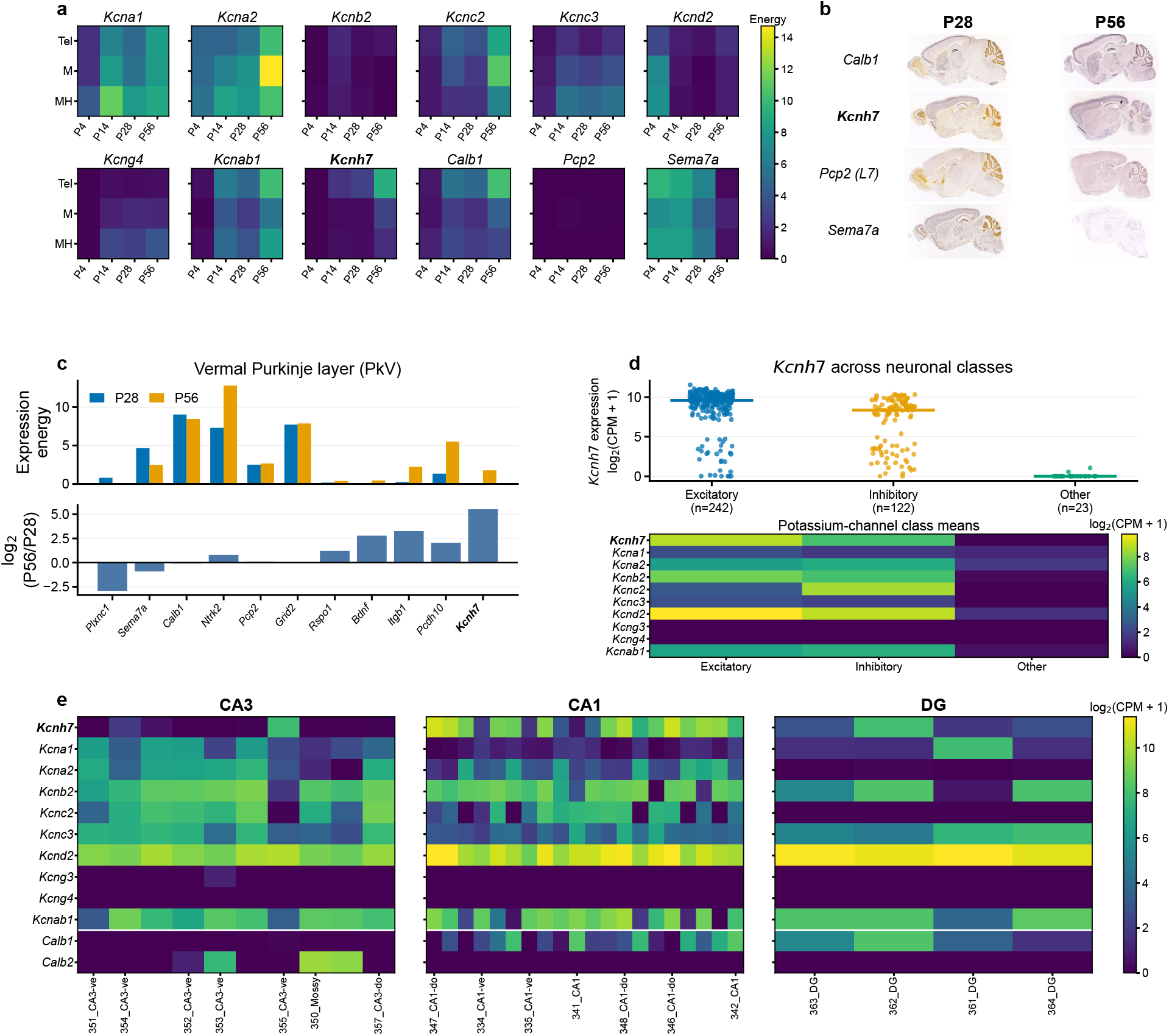
Developmental screening identifies *Kcnh7* as a maturation-associated potassium-channel candidate. a, Mini heatmaps of Allen Developing Mouse Brain Atlas in situ hybridization signal at postnatal day (P) 4, P14, P28 and P56 in the telencephalic vesicle (Tel), midbrain (M) and medullary hindbrain (MH). Values are Structure Unionize expression-energy measurements from one representative Section Dataset per gene–age combination when available. The same three structure IDs were used at all ages. b, Representative sagittal ISH images for *Calb1*, *Kcnh7*, *Pcp2* (L7) and *Sema7a* at P28 and P56. *Calb1* and *Pcp2* are shown as Purkinje-cell reference genes, and *Sema7a* as a developmental comparison. Images in b are from the Allen Institute, Allen Developing Mouse Brain Atlas (P28 and P56) and Allen Mouse Brain Atlas (P56). Source Data Fig. 2 links to each section. These images are excluded from any licence granted by the author and remain subject to the Allen Institute Terms of Use. c, Expression in the vermal Purkinje cell layer at P28 and P56 (PkV, structure 16841). Top, absolute Structure Unionize expression-energy values; bottom, log₂(P56/P28) fold change calculated from the same records. The parallel comparison in the hemispheric Purkinje layer (PkH, structure 16948) is in Source Data Fig. 2. d, Single-cell RNA-seq analysis of the Allen mouse_ctx-hpf_10x dataset. Top, cluster-level *Kcnh7* expression across transcriptomic clusters grouped as excitatory, inhibitory or other; each point represents one cluster. Bottom, class-level mean expression of the displayed potassium-channel panel. Values are trimmed mean log₂(CPM + 1), where CPM denotes counts per million. e, Absolute expression of the potassium-channel panel together with *Calb1* and *Calb2* across CA3-, CA1- and dentate-gyrus (DG)-associated clusters. Clusters are shown in Allen taxonomy order where available. The trimmed mean for *Calb2* was zero in each of the four DG clusters.

I compared P28 and P56 within the same anatomically defined Purkinje cell layer (Fig. 2b,c). In the cerebellar vermis, *Kcnh7* expression energy rose from 0.03846 at P28 to 1.76803 at P56, the largest fold increase among the 11 displayed genes. The same comparison in the hemispheric Purkinje layer gave 0.03947 and 1.00811, respectively. *Plxnc1* and *Sema7a* declined at both sites.

I next examined which neuronal populations expressed *Kcnh7* in the adult Allen mouse isocortex and hippocampal formation taxonomy.^12,13^ *Kcnh7* was detected across both excitatory and inhibitory neuronal clusters (Fig. 2d). All four dentate gyrus clusters had positive trimmed-mean expression of *Kcnh7* and *Calb1*, and both genes reached their highest value in the same cluster (Fig. 2e). The trimmed mean for *Calb2* was zero in all four. In CA1, the cluster with the highest *Kcnh7* expression had a *Calb1* trimmed mean of zero.

Together, the developmental ISH screen identifies *Kcnh7* as a potassium-channel candidate whose expression increases late postnatally.

### ERG3 conductance raises tonic-firing threshold, broadens pre-tonic response diversity and enhances weak rhythmic recruitment

I reanalysed mouse visual-cortex Patch-seq data from the Allen/NeMO release^14^ to identify electrophysiological features associated with *Kcnh7* expression. After adjustment for donor and transcriptomic batch within transcriptomic types, all five pooled partial correlations were negative (Fig. 3a). For short- and long-square threshold currents, the estimates were small (r = −0.029 and −0.014, respectively), and both 95% confidence intervals included zero. The intervals for resting membrane potential and membrane time constant lay below zero. The interval for input resistance included zero. Adding the *Calb1*–*Calb2* expression covariate yielded the same pattern.

**Figure 3.**
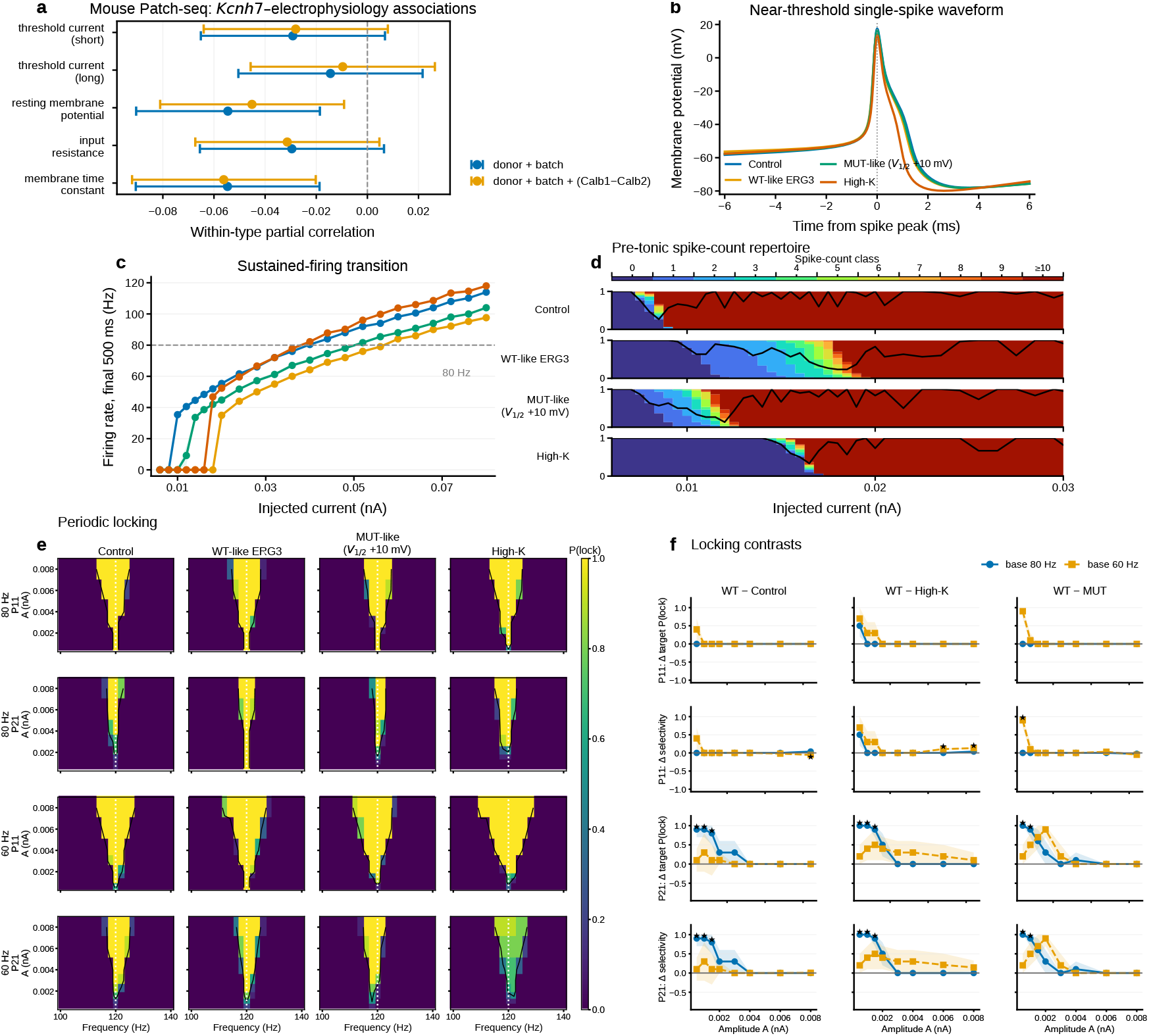
ERG3 conductance raises tonic-firing threshold, broadens pre-tonic response diversity and enhances weak rhythmic recruitment. a, Within-transcriptomic-type partial correlations between mouse *Kcnh7* expression and electrophysiological features in the Allen/NeMO mouse visual-cortex Patch-seq data. Points and intervals show Fisher meta-analytic estimates and 95% confidence intervals. Blue indicates donor and transcriptomic-batch adjustment; orange additionally includes a *Calb1*–*Calb2* expression covariate. b, Near-threshold single-spike waveforms aligned to the spike peak under Control, WT-like ERG3, MUT-like (V₁/₂ +10 mV) and High-K conditions. Control lacked ERG3 conductance; MUT-like used a +10-mV activation-half-voltage shift; High-K increased fast-K conductance without ERG3 gating. c, Final-500-ms f–I curves, showing mean ± sample s.d. across ten seeds. The dashed line marks 80 Hz. d, Spike-count distributions across 30 seeds during 500-ms stimulation. Colours denote counts 0–9 and a pooled ≥10 category. The black line shows the maximum probability among original, unpooled integer count classes. The displayed interval (0.006–0.030 nA) covers the pre-tonic transition. Source Data for Fig. 3d (Source_Data_Fig3d.csv) gives the full current sweep. e, Locking-probability landscapes across ten seeds. Rows show P11 and P21 for 80- and 60-Hz base frequencies. P11 and P21 denote 1:1 and 2:1 modulation-to-spike relationships, respectively. White dotted lines mark target frequencies; black contours indicate P_lock = 0.5. f, WT-like ERG3 minus Control, High-K or MUT-like contrasts in target locking probability and target selectivity; positive values favour WT-like ERG3. Selectivity is target locking probability minus mean off-target probability within the same mode-specific ±20-Hz window, excluding target ±4 Hz. Blue circles and orange squares denote the 80- and 60-Hz analyses. Shading shows 95% seed-paired percentile-bootstrap confidence intervals. Stars indicate Benjamini–Hochberg-adjusted q < 0.05 from exact within-seed condition-label permutation tests; correction families are defined in Methods.

I added a literature-constrained ERG3 conductance to the Khaliq–Gouwens–Raman Purkinje-cell model using the neonatal baseline described by Niculescu et al.^4,15,16^ Four conditions were compared: Control, which lacked the ERG3 component; WT-like ERG3; MUT-like (*V* _1/2_ + 10 mV); and High-K, in which fast-K conductance was increased without ERG3 gating. The MUT-like condition imposed an approximately +10-mV depolarizing shift in activation half-voltage to approximate one electrophysiological feature of the *KCNH7* p.Arg394His variant reported in Amish families with bipolar-spectrum illness.^17^ Only this voltage-dependence shift was represented; the condition was not intended to reproduce the complete molecular or kinetic phenotype of the variant.

Aligned near-threshold single-spike waveforms differed only modestly among the four conditions, with the clearest separation occurring during repolarization (Fig. 3b). Passive-state measurements were more sensitive to the conductance manipulation. Relative to Control, WT-like ERG3 shifted resting membrane potential in the negative direction and reduced input resistance and membrane time constant (Supplementary Fig. S3-1a–c). The shifts in resting potential and membrane time constant agreed in direction with their associations with cortical *Kcnh7* expression (Fig. 3a).

The final-500-ms f–I curves showed that WT-like ERG3 shifted the onset of sustained firing towards larger injected currents (Fig. 3c). High-K also delayed sustained-firing onset, indicating that elevation of the effective firing threshold was not unique to ERG3 gating. The spike-count distributions, however, distinguished the two manipulations. Near the transition into sustained firing, WT-like ERG3 produced an extended interval in which several integer spike-count classes were occupied across trials, whereas this broad pre-tonic repertoire was not reproduced to the same extent by High-K (Fig. 3d). Thus, elevation of tonic-firing threshold was separable from expansion of near-threshold response diversity.

To determine whether this expanded repertoire produced a general increase in readout discriminability, I analysed paired square-step responses across current offsets and cumulative decision windows. Count- and timing-based effects varied with current offset, decision window and score definition rather than showing a uniform directional improvement (Supplementary Fig. S3-2a,b). I therefore tested a more explicitly temporal input structure using periodic current modulation across fine frequency and amplitude grids.

Locking probability depended on both the locking mode and the conductance condition (Fig. 3e). P11 locking at the base modulation frequency was recruited over relatively broad regions and showed limited separation among conditions. By contrast, P21 locking revealed a clearer WT-like ERG3 advantage at low modulation amplitudes. Pairwise contrasts showed increased target-frequency locking probability and target-minus-off-target selectivity for WT-like ERG3 relative to Control and High-K at selected low amplitudes; the WT-like-minus-MUT-like contrast showed a corresponding advantage of WT-like gating (Fig. 3f). The effect was therefore selective for weak-input P21 recruitment rather than a general elevation of locking across all frequencies, amplitudes and locking modes.

Together, these analyses show that ERG3 conductance does not act solely as an outward-current brake. Although both WT-like ERG3 and High-K delayed the transition into sustained firing, only WT-like ERG3 combined this threshold shift with broader pre-tonic response-class occupancy and selective recruitment by weak rhythmic input. This combination was therefore not explained by increased potassium conductance alone.

### *KCNH7*-inspired adaptation differentially regulates learning stability and associative retrieval

ERG3 recruitment depends on preceding activity. To ask what this property contributes to learning, I represented it as slow negative feedback in three reduced models. A decaying penalty reduced hidden-unit responses in Q-learning, a phenomenological slow potassium conductance followed spikes in the spiking network, and recent hidden-unit activity lowered subsequent activation probability in a sampling model. In GridWorld Q-learning,^18^ a decaying hidden-activity penalty was combined with graded ReLU or binary step responses and matched or mismatched transmission-dependent updates (Fig. 4a,b). Under stronger perturbation (noise lower bound 0.60), matched ReLU agents with slow-K initially maintained high returns and short episodes before deteriorating. Relative to slow-K-off agents, the fall in return and rise in episode length occurred later, indicating that adaptive feedback extended the early interval of effective performance rather than merely shifting the final score. Once deterioration occurred, slow-K also reduced its severity in both matched and mismatched agents.

**Figure 4.**
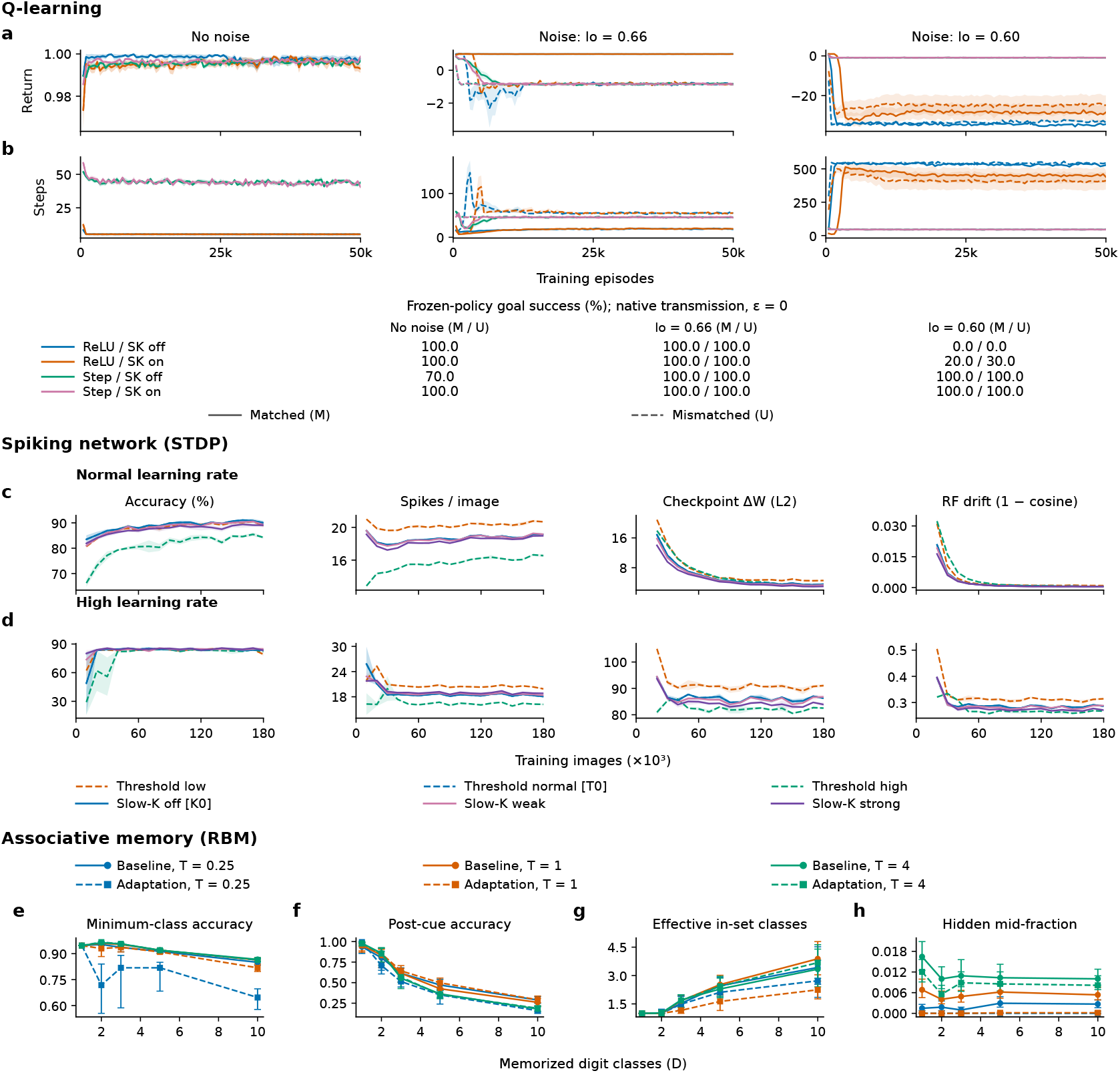
Regime-dependent effects of *KCNH7*-inspired adaptation on learning and associative retrieval. a,b, Mean training return per episode (a) and episode length (b) in GridWorld Q-learning with ReLU or step responses and slow-K off or on. Columns show no perturbation and multiplicative transmission factors sampled from U(0.66, 1) or U(0.60, 1). Solid and dashed lines denote matched and mismatched updates, respectively. Lines and shading show mean ± SEM across ten independently trained seeds per condition. The table reports mean goal success across seeds after 50,000 training episodes, evaluated with frozen parameters, native transmission and no exploration (ε = 0); it is not the endpoint of the training curves. M and U denote matched and mismatched updates. c,d, MNIST spiking-network results at normal (c) and high (d) STDP learning rates. Columns show native-evaluation accuracy, population spikes per 350-ms stimulus plus 150-ms rest cycle, Euclidean weight change between consecutive saved checkpoints, and receptive-field drift defined as the mean of one minus the cosine similarity between corresponding incoming weight vectors. Dashed lines denote nominal threshold manipulations; solid lines denote slow-K manipulations at the normal nominal threshold. Lines and shading show mean ± sample s.d. across three training seeds. Threshold-normal (T0) and slow-K-off (K0) are separately acquired nominal controls and are not pooled. e–h, RBM minimum-class cue-completion accuracy (e), overall terminal accuracy after cue removal (f), effective in-set class number among free-run samples (g), and the fraction of hidden activation probabilities in [0.2, 0.8] (h). Solid circles and dashed squares denote baseline and hidden adaptation, respectively. Training and evaluation used the indicated common temperature. Points and error bars show medians and interquartile ranges across 150 trained models per condition. D denotes the number of trained digit classes, not measured representation dimensionality; D = 1 used an intact cue. Effective class number measures generated class occupancy, not memory capacity. Reduced models were inspired by activity-history-dependent feedback and did not reproduce the full ERG3 conductance.

A related temporal effect was evident at the intermediate perturbation level (noise lower bound 0.66). With mismatched updates, slow-K delayed departure from the initial high-return, short-episode regime and attenuated the transient deterioration. With matched updates, ReLU agents retained high returns and short episodes throughout the 50,000-episode observation period, with no comparable collapse in either slow-K condition. All conditions at this perturbation level nevertheless achieved 100% goal success in the final frozen-policy evaluation under native transmission without exploration. Endpoint success therefore did not capture differences in the time course of performance during continued updating and exploration.

At the stronger perturbation level of 0.60, frozen-policy success after 50,000 episodes increased from 0% to 20% with slow-K in matched ReLU agents and from 0% to 30% in mismatched agents. Step-response agents retained 100% success under this perturbation, although they required more actions than ReLU agents during noise-free training. Slow-K thus provided a condition-dependent buffer for graded value learning: it prolonged an initially effective operating regime and attenuated subsequent deterioration, without universally preventing eventual failure. Its contribution was therefore expressed in the timing and severity of performance loss as well as in final task success.

In an MNIST network trained with spike-timing-dependent plasticity (STDP)^19^, I varied excitatory and inhibitory spike thresholds with slow-K off. In a separate comparison, I held both nominal thresholds at their normal values and varied the conductance recruited by each excitatory spike. At the normal learning rate, high nominal thresholds reduced spike output during checkpoint evaluation, delayed the rise in classification accuracy and yielded lower final accuracy (Fig. 4c). At the high learning rate, accuracy differences narrowed after the initial learning period, while high-threshold networks retained lower spike output and smaller late checkpoint-to-checkpoint weight changes and receptive-field drift (Fig. 4d). Threshold elevation therefore imposed an acquisition cost that depended on the strength of plasticity, rather than simply improving learning stability.

At the normal learning rate, strong slow-K reduced early checkpoint-to-checkpoint weight displacement and receptive-field drift at fixed nominal thresholds (Fig. 4c). From 10,000 to 20,000 presentations, mean drift fell from 0.0208 to 0.0164 and weight displacement from 16.85 to 13.99. Evaluation accuracy changed from 85.2% to 84.0%. At the high learning rate, mean drift was slightly higher with strong slow-K over the first two measurable intervals. At every checkpoint from 40,000 to 180,000 presentations, all three matched seeds had lower drift with strong slow-K (Fig. 4d). Over the prespecified late window (120,000–180,000 presentations), mean drift fell from 0.284 to 0.272 and weight displacement from 86.01 to 83.82, while evaluation spike output rose from 18.39 to 18.92 spikes per image and accuracy changed from 83.93% to 84.64%. Thus, late synaptic reorganization was attenuated at the same nominal thresholds even as the trained network produced slightly more spikes during evaluation.

I then asked whether the same form of history-dependent constraint would remain beneficial when retrieval depended on stochastic state sampling. In a Bernoulli restricted Boltzmann machine (RBM),^20,21^ a sampled hidden unit accumulated a decaying penalty that lowered its probability of firing on subsequent sampling steps. I varied temperature during training and cue-driven image generation and scored the generated images with a common fixed classifier (Fig. 4e–h). Adaptation increased the temperature dependence of completion accuracy. At D = 10 memorized digit classes, median minimum-class accuracy decreased from 85.00% to 64.67% at T = 0.25 and from 86.33% to 81.83% at T = 1, whereas the corresponding values at T = 4 were 86.67% and 86.33% (Fig. 4e). Thus, the adaptation imposed a substantial retrieval cost in the low-temperature regime, but little change in median completion accuracy at the highest temperature tested.

Completion, post-cue generation and spontaneous class occupancy responded differently to adaptation. At D = 10 and T = 1, median post-cue accuracy increased from 26.00% to 29.10%, while the effective number of generated in-set classes decreased from 3.89 to 2.24 (Fig. 4f,g). At T = 0.25, both post-cue accuracy and effective class number decreased, whereas at T = 4 the effective class number increased from 3.34 to 3.68. Adaptation could therefore favour cue-consistent terminal outputs while narrowing spontaneous class occupancy in one regime, yet impair both completion and post-cue output in another. Its effect was not a uniform reduction in the diversity of generated classes.

Hidden intermediate-probability occupancy decreased with adaptation at all three temperatures (Fig. 4h). This decrease coexisted with the pronounced low-temperature impairment, showing that more polarized hidden probabilities were not sufficient for accurate retrieval. Together with the changes in class occupancy, this pattern suggests that history-dependent inhibition can overconstrain reconstruction when sampling is already relatively deterministic, whereas a more stochastic regime can accommodate it with less loss of completion performance. The results support a role for slow-K-like feedback in regulating task-relevant responses: it can buffer learning deterioration or moderate synaptic change, but the same form of constraint can become detrimental to associative retrieval.

### ERG conductance and kinetics shape forecasting, delayed-input reconstruction and firing history

Whether a change in potassium conductance raised or lowered one-step prediction error depended on the input waveform and recurrent connectivity (Fig. 5a). Without recurrence, the lowest mean test mean squared errors (MSEs) among the tested WT ERG multipliers occurred at ×1.5 for the reference Mackey–Glass input A1 (0.073), ×1 for the long-cycle input A2 (0.084) and ×2 for the phase surrogate A3 (0.083). The corresponding ERG-off MSEs were 0.206, 0.278 and 0.224. With recurrence at a 0.001 nS cap, the lowest WT means occurred at ×0.25 for A1 and A3 and at ×0.5 for A2. At the same cap, WT ×1.5 increased mean error relative to ERG off for all three inputs.

**Figure 5.**
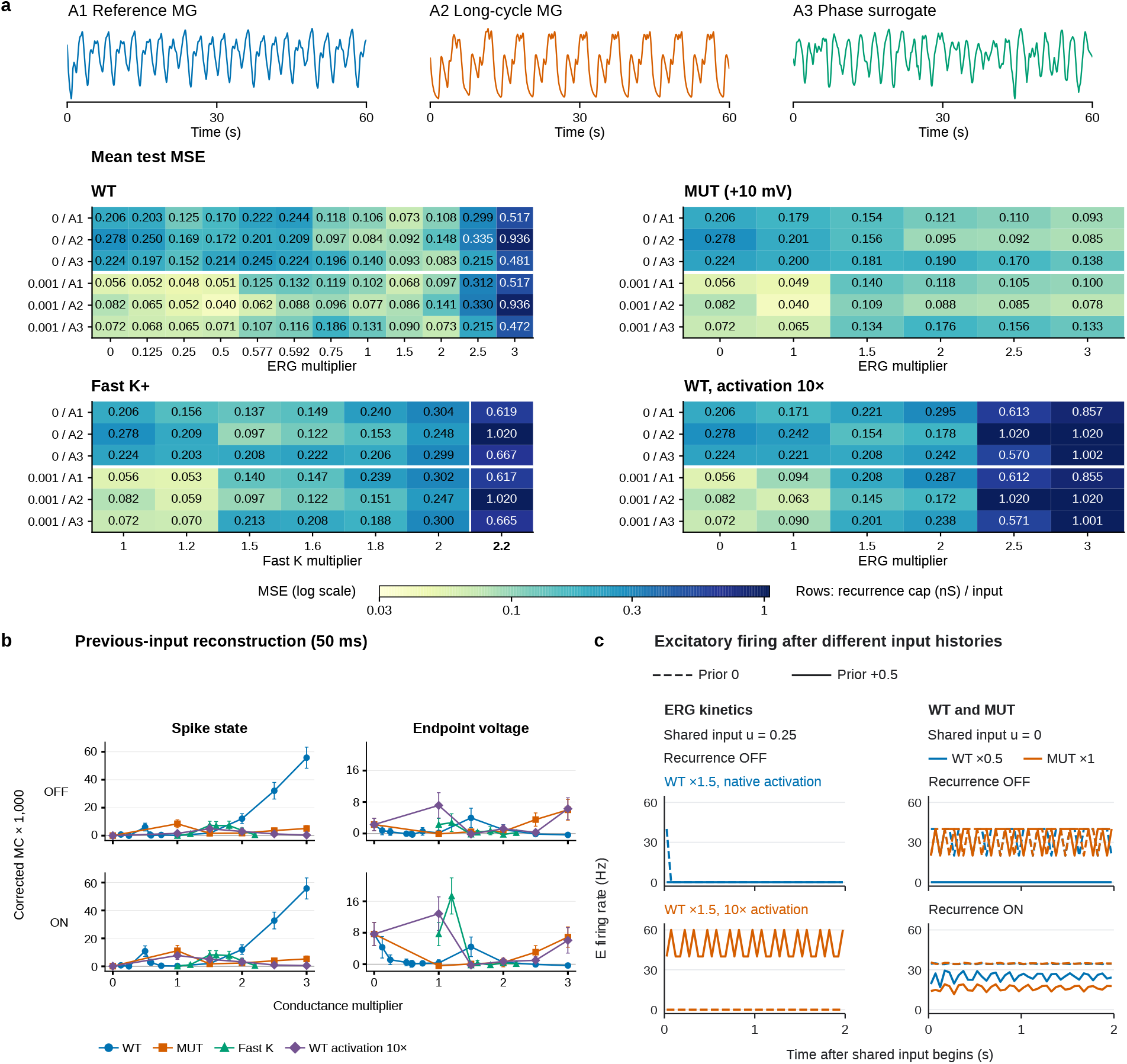
Potassium conductance, single-lag reconstruction and firing responses to input history. a, Held-out one-step mean squared error (MSE) for reference Mackey–Glass (A1), long-cycle Mackey–Glass (A2) and a phase surrogate of A1 (A3). Each heatmap cell is the mean of 30 network seeds. Recurrence caps were 0 and 0.001 nS. Heatmaps share a logarithmic colour scale. The ERG-off condition retains native Fast K⁺ and is labelled Fast K⁺ ×1. The main Fast K⁺ series extends to ×2.2. MUT shifts the ERG activation midpoint by +10 mV. The accelerated WT condition increases both activation rate coefficients tenfold. b, Reconstruction of the preceding independent input sample (d = 1, 50 ms), with background events disabled. Columns show the weighted spike state and endpoint voltage. Rows show the two recurrence caps. The score is the positive part of the observed held-out predictive R² minus the mean positive part across 200 refitted target permutations, displayed ×1,000. The ridge penalty was selected using training data across delays 1–60. Points and bars show means and pointwise 95% t intervals across 30 seeds. ERG off appears at ×0 for ERG and at ×1 for Fast K⁺. c, Excitatory firing following a preceding 1-s input of 0 (dashed) or +0.5 (solid). At time zero, input became identical within each comparison and remained fixed for 2 s. Left, native and tenfold accelerated WT activation at ERG ×1.5 with a common input of 0.25 and recurrence disabled. Right, WT ×0.5 and MUT ×1 with a common input of 0 at the two recurrence caps. Native spike counts were converted to rates per excitatory cell in 50-ms bins. Curves and ribbons show means and pointwise 95% t intervals across 30 matched network seeds. Input level 0 retained 15 pA tonic current per cell. The +0.5 prefix added 15 pA per excitatory cell. The common 0.25 suffix added 7.5 pA.

The ERG activation-midpoint shift and Fast K⁺ conductance changes gave distinct input profiles. At ×1, the +10 mV shift (MUT) increased MSE relative to WT in A1–A3 without recurrence and reduced it in A1–A3 with recurrence. At Fast K⁺ ×2, mean MSE was lower for A2 than for A1 and A3 at both recurrence caps. At Fast K⁺ ×2.2, A2 had the highest mean error (1.020) and produced no excitatory or inhibitory test spikes. At ×2.2, ×2.5 and ×3, Fast K⁺ increased MSE relative to both Fast K⁺ ×2 and ERG off in all 30 paired seeds for each input and recurrence cap. All three input conditions were silent at ×3 (Supplementary Fig. S5-3).

Input structure also changed the ordering of channel conditions. Without recurrence, WT ×2.5 and Fast K⁺ ×2 gave similar MSE for A1 (0.299 and 0.304). WT ×2.5 gave higher error for A2 (0.335 versus 0.248) and lower error for A3 (0.215 versus 0.299). A1 and its phase surrogate A3 had similar autocorrelation decay times (0.455 and 0.452 s) and the same dominant spectral period (2.56 s). A2 had a longer decay time (0.767 s) and dominant period (6.40 s). In an exploratory one-seed comparison of ten fixed waveforms, the direction of the MUT ×1 versus WT ×1 difference was consistent across waveforms within each recurrence cap. In the same comparison, stretching a regularized waveform reversed the WT ×2.5 versus Fast K⁺ ×2 ordering. These manipulations changed several properties of the input together. The shared time scale of A1 and A3 alongside their divergent dose responses suggests that the dominant period and autocorrelation decay alone do not determine the channel effect (Supplementary Fig. S5-4).

In a separate assay using independent random inputs, I decoded the input from the preceding 50-ms bin using the current reservoir state (Fig. 5b). Increasing native WT ERG from ×1 to ×3 increased the permutation-corrected spike-state reconstruction R² for the preceding input by 0.0556 without recurrence and 0.0555 with recurrence, with increases in all 30 paired seeds at each cap. For MUT, the same conductance step increased endpoint-voltage scores by 0.0061 and 0.0073 and decreased spike-state scores by 0.0034 and 0.0059, respectively. At matched ERG ×3, tenfold faster activation decreased spike-state scores relative to native WT by 0.0554 and 0.0553 and increased endpoint-voltage scores by 0.0067 and 0.0064, respectively. In the tenfold-activation series itself, increasing conductance from ×1 to ×3 did not improve endpoint-voltage reconstruction. Voltage at the end of a bin can reflect subthreshold polarisation, spike phase or post-spike repolarisation.

Short-lag reconstruction and forecasting had distinct dose dependences. Raising WT from ×1 to ×3 improved reconstruction of the preceding random input but increased one-step forecast MSE for A1, A2 and A3 in all 30 paired seeds at both recurrence caps. For A2, mean MSE rose from 0.084 to 0.936 without recurrence and from 0.077 to 0.936 with recurrence. At ERG ×3, accelerating activation improved reconstruction from endpoint voltage while increasing forecast error for every input at both caps. Increasing MUT conductance from ×1 to ×3 also improved endpoint-voltage reconstruction at both caps, but lowered forecast MSE only without recurrence. A voltage or spike pattern that distinguished the previous random input therefore did not necessarily provide a better readout of the next sample of a structured input.

To test whether earlier stimulation changed the response to the same later drive, I compared two 1-s input histories. One added 15 pA per excitatory cell to the tonic drive, and the other left the tonic drive alone. For the next 2 s the input and background events were identical within each pair (Fig. 5c). With recurrence disabled and WT ERG at ×1.5, the earlier pulse reduced the mean response during the common drive from two spikes per excitatory cell to zero. Accelerating ERG activation tenfold at the same conductance reversed this effect. After the pulse, excitatory cells fired at a mean 48 Hz over the shared 2-s window, compared with no spikes after the control history. Thus a brief input could silence or recruit subsequent firing in this model depending on activation kinetics, even without recurrent synapses. In the separate comparison at tonic drive alone, the earlier pulse reduced output by 68 spikes per excitatory cell for WT ×0.5 and by one for MUT ×1 without recurrence. With recurrence, the respective reductions were 19.10 and 36.90 (pointwise paired 95% intervals, 18.31–19.89 and 36.14–37.65). The shared-input comparison identifies the change in firing before a readout is applied. Changing ERG activation kinetics could reverse the effect of the preceding pulse on those spikes.

### The timing of ERG modulation alters both ongoing prediction and the readout learned during training

In mouse motor cortex Patch-seq data^22^, the expression score of 18 Gq-associated receptor genes showed positive adjusted rank associations with *Kcnh7* in excitatory and inhibitory cells (0.425 and 0.375, respectively). The pointwise 95% mouse-bootstrap intervals were [0.288, 0.550] and [0.293, 0.459]. The estimates lay above the central 95% ranges of the saved 18-gene scores sampled from the housekeeping reference pool. Associations differed among individual glutamatergic, muscarinic, α1-adrenergic, and serotonergic receptor genes (Fig. 6a,b and Supplementary Fig. S6-2).

**Figure 6.**
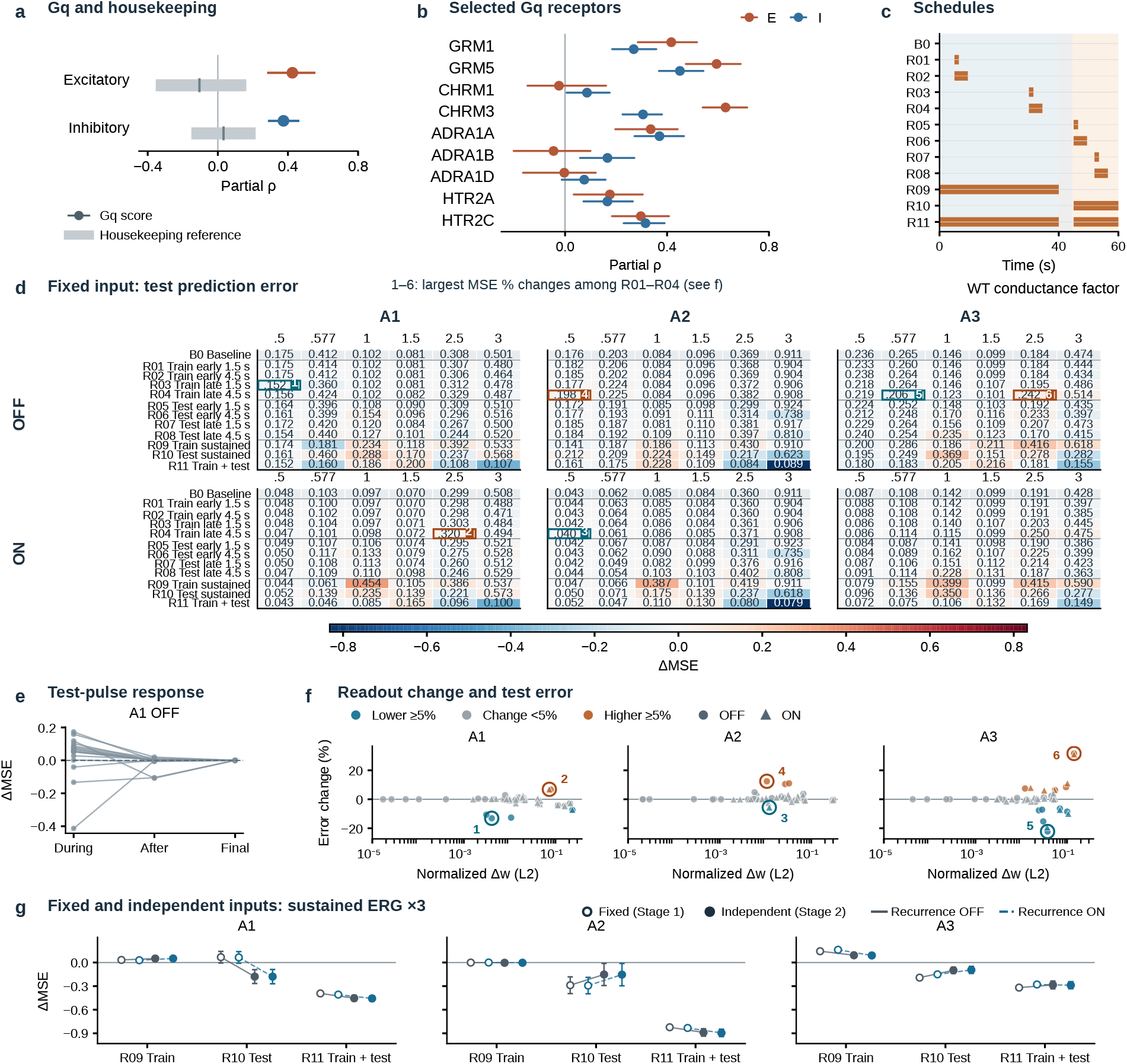
Timing of ERG modulation and prediction. a, Partial rank correlation of *Kcnh7* with an 18-gene Gq-associated receptor score in mouse motor cortex (410 excitatory cells from 150 mice and 817 inhibitory cells from 220 mice). Points and bars show estimates and pointwise 95% mouse-bootstrap intervals. Grey bands show the central 95% range of 1,000 housekeeping reference scores, with their median marked. b, Partial correlations for nine receptor genes, with intervals as in a. Uppercase labels in b identify the corresponding mouse genes. c, Schedules of the reversible +10-mV ERG activation shift, with training, washout and testing at 0–40, 40–45 and 45–60 s. d, Mean absolute test mean squared error (MSE, printed) and change from the corresponding no-shift baseline (colour). B0 denotes the baseline. Numbers 1–6 identify the largest relative improvement and worsening among finite training pulses for each input and link to f. e, Test-pulse MSE differences during the pulse, immediately after offset and in a common final window, for A1 with recurrence OFF. f, Relative change in test MSE and normalized distance from the corresponding baseline readout weights, excluding bias. Colours mark changes of at least 5% in either direction. g, Whole-test MSE effects of shifting the activation midpoint during training, during testing, or during both periods at ERG ×3. The shift was off during the intervening washout in all three schedules. Stage 1 used one fixed waveform per input family across seeds. Stage 2 tested a new realization per family and seed. Points and bars show mean changes from the matched baseline and pointwise 95% paired-seed t intervals. Each model condition used ten network seeds.

In neonatal mouse Purkinje cells, mGluR1 activation reduces the maximal native ERG current and shifts its activation curve towards more positive voltages by approximately 10 mV^4^. To isolate the timing of this gating change, I applied a reversible +10-mV shift to the ERG activation midpoint during selected training or test intervals at fixed maximal conductance. I compared ten network seeds at each input, conductance and recurrence setting (Fig. 6c,d). With A2 at ERG ×0.5, a 4.5-s late-training pulse increased mean test MSE from 0.1764 to 0.1983 without recurrence and decreased it from 0.04272 to 0.04033 with recurrence (+12.45% and −5.61%). The ×0.577038 setting came from an A1 pilot in which WT conductance was calibrated to approximate the mean ERG current at MUT ×1. In A3 without recurrence, the same pulse lowered MSE by 22.27% at ×0.577038 and increased it by 31.66% at ×2.5. These examples were selected by the magnitude and direction of the observed error change within each input.

With a fixed readout, test-time perturbation effects depended on the measurement window (Fig. 6e). A late 1.5-s A1 pulse without recurrence increased MSE during the pulse by 0.159 at ERG ×1 and decreased it by 0.414 at ×2.5. Fourteen of the 16 displayed dose–schedule conditions had zero saved MSE difference in the common final window in every seed. The remaining two mean differences were +0.00109 and −0.00098. For training pulses, similar distances from the matched unperturbed weights could accompany opposite changes in test error (Fig. 6f). In A3 with recurrence at ×3, early and late 4.5-s pulses gave similar normalized weight distances (0.1068 and 0.1021) but opposite changes in MSE (−10.03% and +11.09%). Each direction occurred in all ten seeds.

The fixed-waveform experiment isolated the effect of changing the activation schedule while ten network seeds received the same input. To ask which effects persisted for other examples of each waveform family, I repeated the grid with a new input for each of the same ten seeds, using that input for every comparison within the seed (Fig. 6g and Supplementary Fig. S6-1). At ERG ×3, the shift lowered MSE for A1, A2 and A3 when it was applied during training and reapplied during testing. This held in all ten pairs in each stage with and without recurrence. For A3, limiting the shift to training instead raised MSE in every pair. Without and with recurrence, the mean increases were 0.143 and 0.162 for the fixed input and 0.092 and 0.089 for the new inputs. By contrast, the effect of shifting activation only during the A1 test changed sign across stages. Mean MSE rose by 0.068 and 0.065 for the fixed waveform but fell by 0.179 at both settings across the new waveforms, with improvement in nine of ten pairs at each setting. For A3, prediction therefore depended on whether the training shift was reinstated at test. For A1, the direction of the test-only effect depended on the waveform sampled.

### Synaptic perturbations and coordinated population drive reveal distinct sensitivities of fixed readouts

I first tested whether a trained circuit could both predict accurately and maintain its prediction under fluctuations in recurrent transmission. For each channel setting, I kept the readout trained on unperturbed transmission fixed. Figure 7a shows its unperturbed prediction error (MSE, horizontal axis) and the distance its prediction moved when individual synaptic events varied in strength (waveform RMS, vertical axis). Under low background noise, adding ERG ×1.5 reduced mean MSE from 0.260 to 0.0436 for A1 and from 0.613 to 0.0296 for A2 relative to ERG off. Mean waveform RMS also fell, from 0.0298 to 0.0106 and from 0.0276 to 0.0102, respectively. Increasing ERG further to ×3 produced similarly small displacements but raised MSE to 0.579 and 0.683. At Fast K⁺ ×3, both low-noise trajectories were silent in every seed, giving zero displacement and mean errors of 1.025 and 0.990. At the same Fast K⁺ dose under high background noise, the mean displacements were 0.0010 and 0.0017, while mean MSE remained high (0.973 and 0.982). A small displacement therefore occurred both when prediction was accurate and when the circuit had ceased to spike.

**Figure 7.**
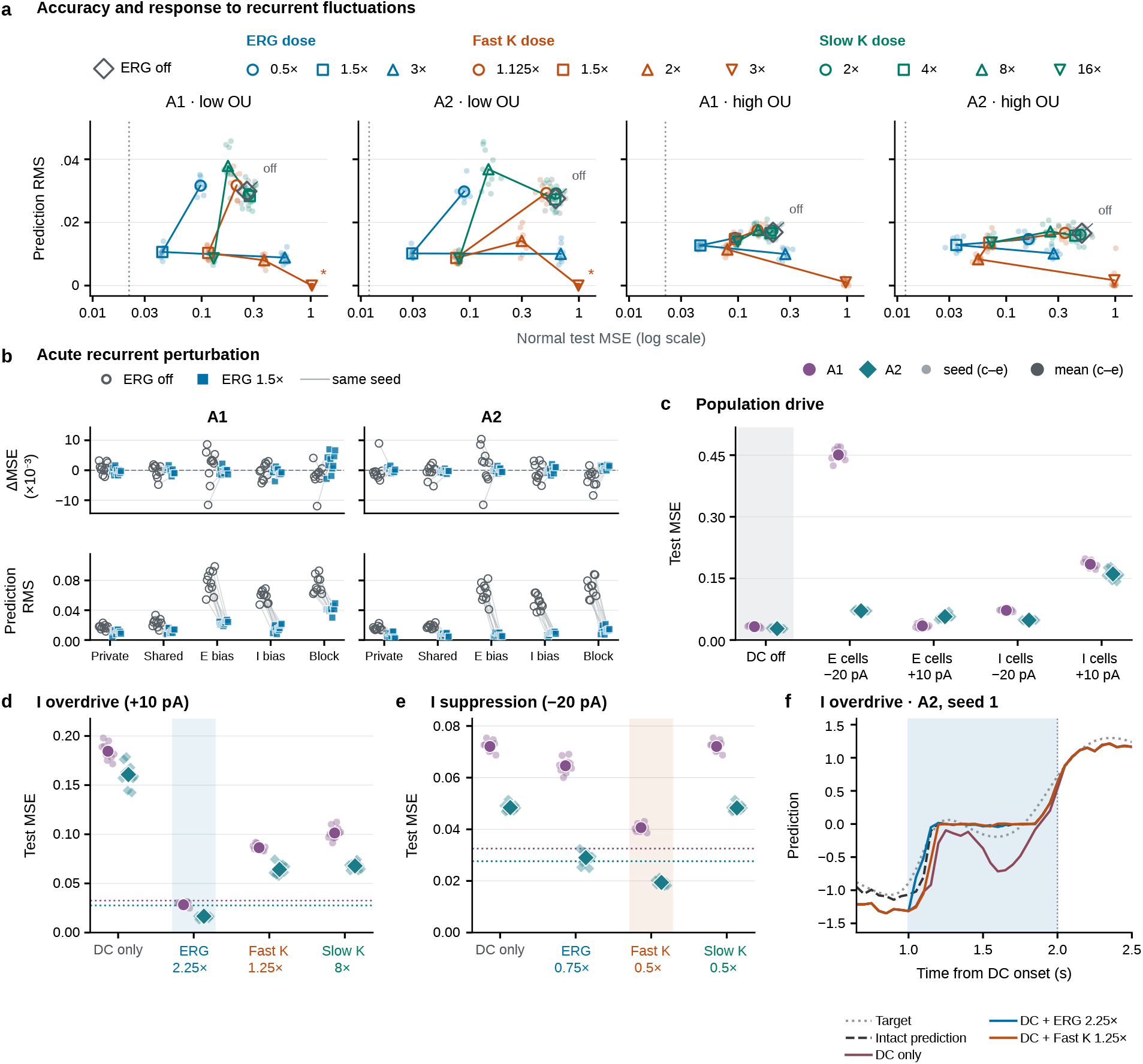
Prediction under synaptic and population perturbations. a, Unperturbed test mean squared error (MSE) versus the root mean square (RMS) difference between Normal and Uniform transmission predictions with recurrence ON. Dots show ten seeds, outlined markers show means and dotted lines show persistence MSE. Asterisks mark Fast K⁺ ×3 conditions in which both trajectories were silent in all ten seeds. b, Acute test-onset transmission perturbations. Lines pair ERG off and ERG ×1.5 within input and seed. The upper row shows target MSE minus each setting’s unperturbed target MSE. The lower row shows the RMS difference between perturbed and unperturbed predictions. Private and Shared denote independent and shared-bin fluctuations, E bias and I bias target the indicated presynaptic population, and Block prevents new recurrent transmission. In a,b, each channel condition uses its own Normal-trained readout. c, Absolute MSE under current applied to every cell of the indicated population for 0–2 s. DC off is the matched intact reference. d,e, MSE during continued I-cell drive. At 1 s, ERG, Fast K⁺ or Slow K⁺ was changed in I cells at the indicated multiplier. Dotted lines show intact means. f, Example A2 predictions for seed 1 under +10-pA I-cell drive. Shading marks conductance modulation. Current and modulation ended at 2 s. The Fast K⁺ trace retains background ERG ×1.5. Panels b–e use the 1–2-s window, low Ornstein–Uhlenbeck (OU) noise and ten paired seeds per input. Panels c–f use the same native ERG ×1.5 readout within each input and seed.

I next perturbed recurrent transmission only after the test began, so that each challenged trajectory had the same preceding activity as its own unperturbed reference (Fig. 7b). ERG ×1.5 produced a smaller waveform displacement than ERG off in every paired seed for both inputs and all five perturbations. After recurrent transmission was blocked, mean waveform RMS fell from 0.0737 to 0.0412 for A1 and from 0.0702 to 0.0136 for A2. Despite this smaller displacement, the change in MSE from each setting’s own unperturbed baseline was more adverse with ERG ×1.5. The between-setting differences in this change were +0.00452 for A1 and +0.00356 for A2 (pointwise paired 95% intervals, 0.00025–0.00879 and 0.00142–0.00571). Absolute MSE under block remained lower at ERG ×1.5 for both inputs. The intervals for the other eight input–perturbation contrasts in MSE change included zero.

I then left synaptic transmission intact and applied the same current to every excitatory cell or every inhibitory cell, holding the native ERG ×1.5 readout fixed (Fig. 7c). Without applied current, mean MSE in the 1–2-s window was 0.0325 for A1 and 0.0276 for A2. Applying −20 pA per excitatory cell raised these means to 0.4507 and 0.0709, whereas +10 pA gave 0.0344 and 0.0570. Applying the same two currents to inhibitory cells gave 0.0721 and 0.0484 at −20 pA, and 0.1845 and 0.1607 at +10 pA. Among these four challenges, excitatory suppression produced the largest error for A1 and inhibitory excitation produced the largest for A2. The two current polarities differed in magnitude.

The relative effects of potassium conductance interventions depended on the direction of drive. During +10-pA inhibitory drive, raising ERG from ×1.5 to ×2.25 reduced mean MSE to 0.0283 for A1 and 0.0166 for A2. The corresponding means were 0.0864 and 0.0642 for Fast K⁺ ×1.25 and 0.1013 and 0.0678 for Slow K⁺ ×8. The paired mean ERG-minus-Fast K⁺ differences were −0.0580 for A1 and −0.0476 for A2, with lower ERG error in every seed (Fig. 7d). During −20-pA drive, reducing ERG to ×0.75 gave higher mean errors than reducing Fast K⁺ to ×0.5, again in all ten seeds for each input. The differences were +0.0240 and +0.00960. Reducing Slow K⁺ to ×0.5 changed the mean error little relative to continued drive alone (Fig. 7e). Coordinated inhibitory drive disrupted prediction. At the doses tested, increasing ERG reduced error more than increasing Fast K⁺ during positive drive, whereas reducing Fast K⁺ gave lower error than reducing ERG during negative drive (Fig. 7d–f).

### The timing of modelled receptor-linked ERG3 modulation shapes state retention and value updating

I modelled a hypothesised Gq-linked receptor shift in ERG3 activation and asked whether sustained loss of that shift could make a circuit leave an established activity state. In a model with two competing populations, the receptor signal q shifted the ERG3 activation midpoint towards more positive voltages at fixed maximal conductance. Removing that shift increased outward ERG3 current at a matched voltage and reduced effective drive to the initially favoured population. Under sustained loss, all 48 trials retained that state at a background drive of −0.55. At −0.65, four of 48 planned trials failed to establish it. Of the 44 that did, 14 exited within 120 s and 30 retained it. At −0.70, none established the initial state (Fig. 8a). At the intermediate drive of −0.65, intact modulation and a constant drive that compensated for signal loss at the intact reference voltage each supported retention in all 48 trials (Fig. 8a and Supplementary Table 2). These comparisons distinguish failure to establish a state from loss of a state already established.

**Figure 8.**
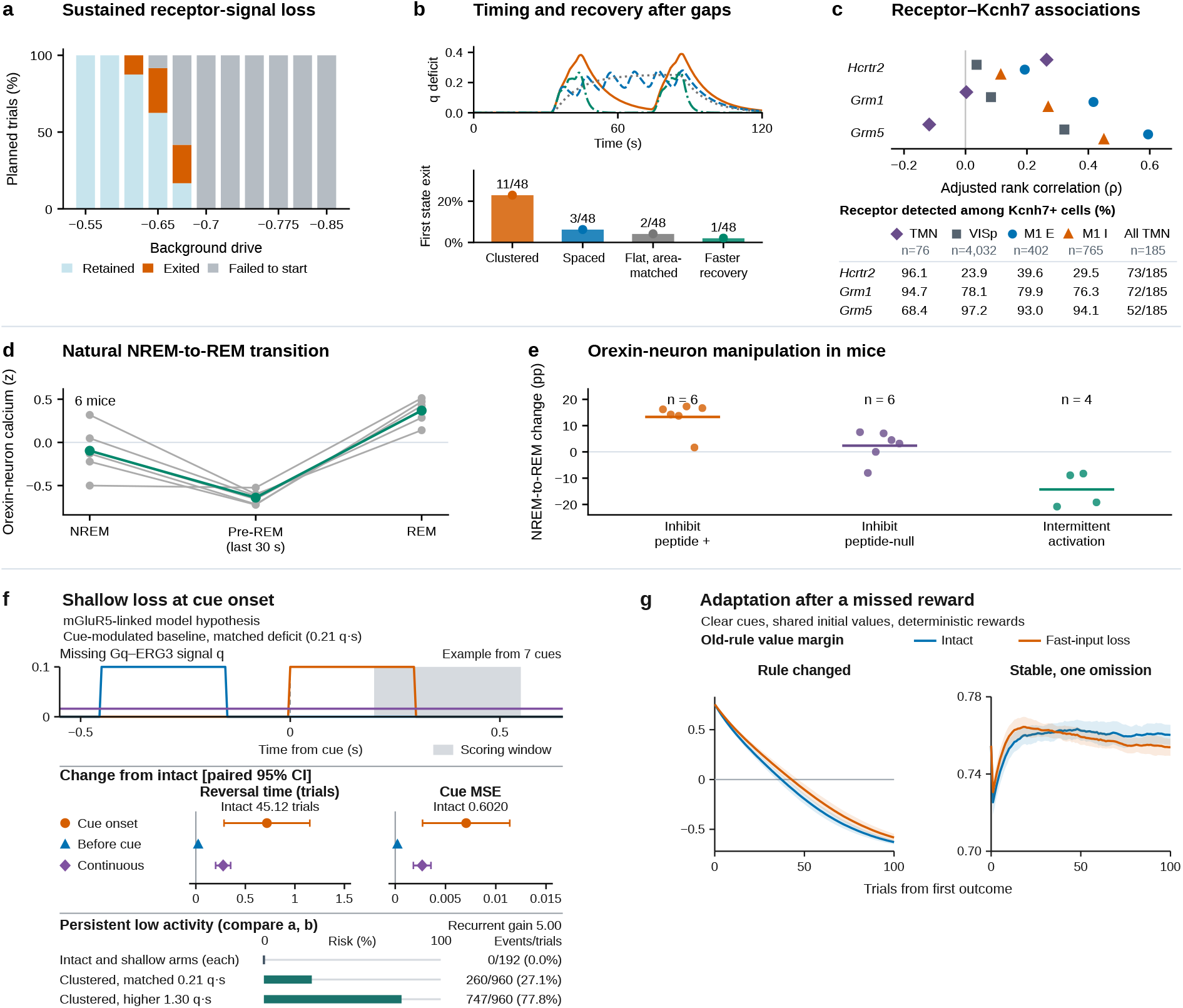
Receptor signal history and value updating. a, Outcome after sustained signal removal across 48 planned simulations per background drive (12 circuits, four noise realizations each). At −0.65, 30 retained the initially favored state, 14 exited after establishing it and four failed to establish it. Supplementary Table 2 compares sustained loss with intact modulation and tonic drive compensation at the same background drive. b, Effective signal deficit q and fraction with a first state exit within 120 s. Clustered and spaced schedules each contain six 3-s gaps. The flat control matches the integrated filtered deficit of the clustered schedule, and faster recovery follows the clustered input. c, Within-cohort adjusted transcript rank correlations and receptor detection percentages among *Kcnh7*-positive cells in tuberomammillary nucleus (TMN), visual cortex (VISp) and motor cortex (M1). The rightmost column gives counts among all 185 TMN cells. d, Orexin-neuron calcium in NREM, the last 30 s before REM and REM. Gray lines are individual mice and green denotes the mean (n = 6). e, Change in the fraction of NREM bouts followed by REM after orexin-neuron inhibition in peptide-expressing and peptide-null mice (n = 6 each), and after intermittent activation in a separate cohort (n = 4). Points denote mice and lines show means. f, Cue-timed, cue-preceding and continuous shallow losses of equal area (0.21 q·s). Gray shading marks the scoring window. Points show changes in reversal time and cue mean squared error (MSE) from intact modulation with paired 95% t intervals across eight circuits. Lower bars show persistent low-activity counts at recurrent gain 5.00 (192 trajectories per intact or shallow arm, 960 per clustered arm). g, Mean old-minus-new action value before the first outcome (trial 0) and after successive trials. Both branches began with the same forced unrewarded choice. The rule then changed (left) or remained stable (right). Shading shows pointwise 95% t intervals across eight circuits, each with 512 sessions per condition. Value reversal required a negative old-minus-new margin for five consecutive trials. The right ordinate is expanded. Panels d,e reanalyze published mouse-level data^8^. pp, percentage points.

I next asked whether the same total interruption time could have different effects on an established state. At partial receptor efficacy, I introduced six 3-s interruptions with the same first and last onset and varied the spacing of the intervening gaps. Short intervals within each cluster produced 11 first exits in 48 trials, compared with 3/48 when the interruptions were spaced farther apart. Spreading the same integrated deficit in the filtered signal q over the full interruption period gave 2/48 exits. Increasing the recovery rate between clustered interruptions gave 1/48 exits and reduced the integrated deficit (Fig. 8b). At matched integrated deficit, a concentrated loss therefore produced more first exits than a flat loss over the same period.

I used public mouse transcriptomic data to examine which receptors were expressed alongside *Kcnh7*^14,22,23^. The adjusted within-cohort rank correlations with *Grm5* ranged from 0.32 to 0.59 in cortical Patch-seq cohorts and were −0.12 in tuberomammillary nucleus (TMN) histaminergic cells. The corresponding *Hcrtr2* estimates were 0.04–0.19 in cortex and 0.26 in TMN. Among the 76 TMN cells with detectable *Kcnh7*, *Hcrtr2*, *Grm1* and *Grm5* were detected in 73, 72 and 52 cells, respectively (96.1%, 94.7% and 68.4%). Across the cortical cohorts, *Grm5* was detected in 93.0–97.2% of *Kcnh7*-positive cells and *Hcrtr2* in 23.9–39.6% (Fig. 8c). This descriptive reversal in the receptor ranking identifies candidate cell populations for testing receptor-dependent ERG3 current.

I reanalysed published mouse data to examine orexin-neuron activity around the NREM-to-REM transition and how manipulating these neurons affected the probability of that transition^8^. In each of the six mice, mean orexin-neuron calcium activity during the last 30 s of NREM before spontaneous REM onset was lower than in NREM overall, and the mean during REM was higher than in that pre-REM window (Fig. 8d). During orexin-neuron inhibition, the percentage of NREM bouts followed by REM rose by 13.3 percentage points relative to baseline in peptide-expressing mice and by 2.4 points in the peptide-null group. Intermittent activation in a separate cohort reduced the proportion by 14.3 points (Fig. 8e).

To test how the same gating assumption could affect task processing, I applied shallow losses of equal integrated q to a recurrent excitatory–inhibitory circuit and held the cue decoder fixed. Loss at cue onset increased cue mean squared error (MSE) by 0.00705 (95% confidence interval, 0.00272 to 0.01139) and prolonged value reversal by 0.717 trials (0.284 to 1.151) relative to intact modulation. At the same integrated deficit, reversal was 0.69 trials slower after loss at cue onset than after loss before the cue (0.26 to 1.13) and 0.44 trials slower than under continuous loss (0.08 to 0.81). The paired intervals for changes in cue-sign accuracy included zero in both timing comparisons. Persistent low activity was absent in all shallow conditions at each tested gain (0/192 per condition and gain). At recurrent gain 5.00, delivering the same integrated deficit as a single contiguous withdrawal produced persistent low activity in 260/960 trajectories, rising to 747/960 at the larger deficit (Fig. 8f). The direction of the shallow cue-timing effect reversed with a tonic reference or greater loss depth (Supplementary Fig. S8-1). Thus, a brief loss can alter task-related output and value updating within a retained activity regime, whereas a sufficiently concentrated interruption can move the same circuit into persistent low activity.

I next asked whether the effect on learned values depended on whether adverse feedback signaled a persistent change. Using archived cue scores, I compared intact modulation with selective removal of the modeled fast input to the ERG3 gating pathway. Both models started with the same action values and experienced the same first omitted reward. With deterministic feedback after that trial, the loss model reached the internal value-reversal criterion after 48.09 trials versus 42.63 for the intact model (paired difference, 5.46 trials, 95% confidence interval, 1.98 to 8.93). Across the first 20 trials, loss increased the old-rule value margin more after true reversal than after a single omission under the stable rule (interaction, +0.0290, 0.0113 to 0.0467). This interaction had the same direction across reward-reliability, cue-mixing and starting-value sensitivities, with all 12 nominal intervals excluding zero. These settings share the same eight circuits. Under the unchanged rule, sustained false value reversals occurred in 0/4,096 replays per model. Sampled choices satisfied a separate eight-of-ten switch criterion in 134/4,096 replays per model, and the omission produced a small transient change in value (Fig. 8g). The divergence between the two environments was therefore clearest in the trajectory of internal values and the time to update them.

## Discussion

Postnatal maturation can make individual synapses faster and more reliable, as demonstrated at the calyx of Held.^3^ What does that gain in fidelity buy a network that learns or computes through recurrent interactions? In the layered network, learning remained stable when the update reused the perturbation that had affected the forward pass (Fig. 1). An acquired solution deteriorated when updating omitted that perturbation or sampled it independently. Variability confined to the update pathway was also compatible with stable learning. In a recurrent circuit with a fixed readout, the question shifts to whether the population still represents the information that the readout was trained to extract. This distinction suggests that maturation may improve computation both by refining transmission at individual synapses and by keeping population activity within a range that downstream circuits can interpret.

A recurrent circuit can tolerate considerable fluctuation in individual inputs if their effects cancel at the readout or fall in dimensions that it scarcely uses. A coordinated change in inhibition, by contrast, can move many task-relevant features together and distort the prediction. The waveform and error measures in Fig. 7 make this distinction concrete. ERG ×1.5 reduced how far the prediction moved after acute transmission block, even though the increase in target error from its own baseline was larger. The direction of the displacement relative to the original prediction error mattered as well as its size. This suggests an experimental criterion for disruptive perturbations: at comparable levels of synaptic variability and mean firing, ask whether the perturbation changes the component of population activity used by the trained readout. A change in its projection would help explain why a circuit can tolerate some forms of transmission noise yet remain vulnerable to a coordinated shift in population drive.

ERG3 offers a way to connect this population-level vulnerability to membrane physiology. Near the transition to sustained firing, a history-dependent potassium current can change which inputs recruit spikes and how long preceding activity influences the response. A receptor-dependent shift in ERG3 gating could therefore alter when a population contributes to the circuit and how strongly it does so. If the shift occurs at different times in excitatory and inhibitory populations, their relative recruitment could change even while individual synapses continue to transmit reliably. The resulting failure would arise from the timing and distribution of the modulation. This hypothesis predicts that a brief mismatch between populations could have a larger effect than a comparable change in average modulation applied coherently across the circuit. The conductance and population-drive results motivate this comparison. Establishing it would require varying the relative timing directly while measuring the population response and its projection onto the readout.

The Gq–phospholipase C–PKC pathway is a plausible molecular route for such timing dependence. Group I metabotropic glutamate receptors can recruit PKC through diacylglycerol and calcium signalling. In hippocampal neurons, metabotropic and ionotropic glutamate inputs can cooperate to recruit PKCγ, providing a precedent for kinase activation that depends on the coincidence of receptor input and neuronal activity.^24^ And in neonatal Purkinje cells, mGluR1 stimulation shifts native ERG activation to more positive voltages and reduces maximal ERG current.^4^ The models isolate the activation shift at fixed maximal conductance. Whether OX2R or mGluR5 engages a PKC-dependent change in ERG3 current in the populations studied here remains an experimental question. Measuring the current during repeated receptor stimulation, with cell-specific tests of *KCNH7* and PKC dependence, would determine whether the proposed pathway carries a history of receptor input on the timescale relevant to circuit operation.

The state-maintenance and task models in Fig. 8 suggest two ways that disrupted modulation could become visible as a phenotype. A signal that supports an ongoing state must be renewed before the circuit loses that state. Interruptions that arrive in close succession can exhaust this support even when their integrated deficit in the modelled signal matches that of a less disruptive schedule. This gives the orexin comparison a specific implication. The distribution of gaps in receptor signalling, together with the recovery of the downstream current, could help determine the probability of an abrupt transition. Posterior hypothalamic OX2R restoration supports sustained wakefulness in mice^7^, and manipulation of orexin neurons changes the probability of transitions from NREM to REM sleep.^8^ These are distinct physiological endpoints. They motivate testing whether the spacing of orexin input influences state retention through a common postsynaptic current in a defined target population.

When modulation is recruited around a task cue, its loss could instead alter how evidence is assigned to an action. In the model, cue scores inform the updating of action values. Their alteration can therefore change how successive outcomes accumulate even when the learning rule itself is held fixed. Following a genuine reversal, the old values persisted longer after removal of the modelled fast input. Isolated adverse feedback under an unchanged rule did not produce a corresponding sustained reversal of internal values. This suggests a particular route to reduced cognitive flexibility: a perturbation upstream of value updating could prolong reliance on the old rule even when that rule was initially learned and the circuit remains active. The difference in Fig. 8g was clearest in internal values. Its behavioural prediction is a change in the time course of choices after sustained evidence, which should be evaluated separately from switching after an isolated misleading outcome.

This distinction gives the schizophrenia comparison greater specificity. Probabilistic reversal studies have reported difficulty adapting after contingency changes^25^, and some clinical cohorts also switch away from appropriate choices too readily after misleading feedback.^26^ An overall switching rate can obscure these differences. The model identifies one candidate process for the first pattern: altered representation of task evidence can slow the revision of established values. A translational test would measure the persistence of the old choice after a true reversal and the tendency to abandon a still-valid choice after misleading feedback in the same task. The receptor–channel hypothesis would gain support if manipulating the candidate pathway selectively changed the predicted component of behaviour.

If cognitive inflexibility reflects insufficient modulation during the period when task evidence is processed, the pharmacological objective would be to restore the response to endogenous glutamate within that period. There is a precedent for a selective cognitive effect. The mGluR5 positive allosteric modulator LSN2463359 improved reversal learning in a neurodevelopmental rat model, with a different outcome in an acute pharmacological model whose behavioural deficit had a different profile.^27^ There are also concrete safety constraints. The mGluR5 modulator 5PAM523 produced neurotoxicity in rats despite lacking detectable intrinsic agonism in the reported cellular assay.^28^ Development of RG7342 stopped after a first-in-human study because its long half-life posed challenges in the context of observed adverse events.^29^ Conversely, VU0409551 showed that a compound could retain cognitive effects in rodents while sparing potentiation of mGluR5-dependent NMDA currents in the tested preparation.^30^ These studies make both the signalling profile and the duration of exposure relevant to what constitutes a milder intervention.

A testable approach would be to look for modest, glutamate-dependent potentiation that restores the candidate current response around relevant activity, while preserving the ability to disregard an isolated misleading outcome. Both behavioural measures matter. A manipulation that accelerates reversal at the cost of frequent inappropriate switching would simply move the circuit to a different point in the trade-off. The same principle applies to state maintenance, where the relevant benefit is sustained activity through ordinary interruptions while retaining the capacity to transition when conditions change. The broader implication is that intrinsic modulation may help circuits determine which changes to absorb and which changes to act on. ERG3 provides a candidate mechanism through which that distinction can be tested from membrane current to population activity and behaviour.

Taken together, these analyses suggest that a circuit can become more reliable by preserving the population activity needed for learning and readout, even while individual signals remain variable. A history-dependent conductance could contribute to this reliability by changing how the circuit responds to a disturbance at a particular moment. The distinction is between preserving the state in which computation takes place and revising a decision within that state. In the models, both depend on when modulatory support is lost and on the activity it normally supports. ERG3 provides a candidate mechanism for testing whether the timing of a receptor-evoked current helps a circuit maintain an ongoing state, absorb an isolated disruption or revise an established choice when new evidence persists.

## Methods

### Supervised learning under controlled forward–backward mismatch

Supervised analyses used the MNIST handwritten-digit dataset^31^ and a fully connected classifier with architecture 784–50–50–10, hidden ReLU responses capped at 500, and a softmax cross-entropy loss. Inputs were scaled to [0, 1]. Weights were initialized independently from a zero-mean Gaussian distribution with s.d. 0.01 and biases were zero. Each of the 60,000 updates used 100 training examples sampled without replacement within that minibatch. Evaluation used a fixed subset of 2,000 test images for each seed after update 1 and every 600 updates. Standard stochastic gradient descent was used in all conditions. In the mixed-learning-rate rendering shown in Fig. 1a,b, the weight-perturbation mismatch condition used a learning rate of 0.1, whereas all other supervised conditions used 0.5.

Each seed specified the same initial weights, training minibatches and test subset across conditions. In the five conditions run at learning rate 0.5, each affine layer propagated the upstream error through the effective weights used in its forward pass. The conditions differed in the multiplicative factors applied to the forward weights and to the local weight gradient. Control used unit factors for both. Update-only perturbed the local gradient alone. Matched reused the forward factors in the local gradient. Forward-only perturbed the forward weights and used unit factors for the local gradient. Independent drew separate factors for the two operations. The mapping-mismatch condition perturbed the forward weights, used unit factors for the local gradient and propagated the upstream error through the original weights. This condition also used the lower learning rate specified above.

Whenever a transmission or local-update mask was sampled, a uniform factor with mean 1 and coefficient of variation 0.55 was drawn independently for each weight. The layer shared the mask across examples in that minibatch. Each transmission condition was tested under three update regimes. In auxiliary plasticity off, the auxiliary plasticity term was disabled while standard gradient learning was retained. In the immature update regime, the auxiliary process was configured with low trace persistence, stochastic update variability, and probabilistic update failure (rho = 0.1, cv_plas = 0.3, p_fail = 0.1, clip = None). In the mature update regime, it was configured with high trace persistence and stable updates (rho = 0.9, cv_plas = 0, p_fail = 0, clip = 1.0). These settings were used as relative computational regimes to compare mismatch sensitivity under unstable versus stabilized update dynamics rather than as direct biophysical fits.

To evaluate whether the learned mapping itself remained coherent, I measured clean test accuracy, defined as performance when forward perturbations were temporarily disabled at evaluation. To ensure that evaluation did not alter the subsequent training trajectory, the layer RNG state, affine-layer buffers, and forward-noise settings were snapshotted before clean evaluation and restored immediately afterward. Weight growth was summarized as the square root of the sum of squared entries across the three weight matrices. Biases were excluded. Curves in Fig. 1a,b show mean ± s.d. across ten seeds, using the sample s.d.

### Developmental ISH and single-cell RNA-seq analyses

I retrieved in situ hybridization records from the Allen Brain Map API. The P4–P28 experiments belong to the Allen Developing Mouse Brain Atlas. All 19 selected P56 experiments are registered in both the developing atlas and the Allen Mouse Brain Atlas.^10,11,32,33^ A potassium-channel candidate panel was examined at P4, P14, P28 and P56 in the telencephalic vesicle, midbrain and medullary hindbrain. For each gene–age combination, one representative Section Dataset was selected when available, prioritizing sagittal delegate experiments. Structure Unionize expression energy was extracted using the same Allen structure identifiers from the Developing Mouse 3 graph (graph 17, ontology 12) at all four ages: 15739 (telencephalic vesicle), 16649 (midbrain) and 17352 (medullary hindbrain). Source Data Fig. 2 lists the selected Allen records, structures and original record selection (3 September 2026) and the matched-structure extraction (18 November 2026).

Representative sagittal ISH images for *Calb1*, *Kcnh7*, *Pcp2* and *Sema7a* were obtained from the selected P28 and P56 Section Datasets. For each experiment, sections were sorted by section number and the middle entry was displayed. Quantitative comparisons were based on Structure Unionize measurements rather than image-pixel intensity.

For the Purkinje-layer comparison, Structure Unionize expression-energy values were extracted at P28 and P56 from the vermal Purkinje cell layer (PkV, structure 16841) for *Kcnh7*, Purkinje-cell reference genes and selected developmental, synaptic-contact and pruning-related comparison genes. The displayed panel included *Kcnh7*, *Calb1*, *Pcp2*, *Grid2*, *Plxnc1*, *Sema7a*, *Ntrk2*, *Rspo1*, *Bdnf*, *Itgb1* and *Pcdh10*. The same experiments were also evaluated in the hemispheric Purkinje layer (PkH, structure 16948). For each gene, the P56/P28 change was calculated as

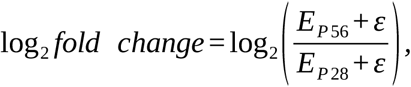

where *E* denotes Structure Unionize expression energy and *ε* is the fixed pseudocount used to avoid division by zero. The pseudocount was 10⁻⁶. All selected structure identifiers and measurements are provided in Source Data Fig. 2.

Single-cell analyses used the Allen mouse_ctx-hpf_10x transcriptomic taxonomy and associated cluster metadata.^12,13^ Trimmed mean expression values were used as provided and are reported as log₂(CPM + 1). Clusters were grouped as excitatory, inhibitory or other using the Allen class label for each cluster. Class-level values were calculated by averaging the displayed cluster-level trimmed means within each broad class.

For hippocampal analysis, the CA1, CA3 and dentate gyrus groups were selected from the Allen subclass and cluster labels. Allen dendrogram order was retained when available; otherwise, the source cluster order was used. All three heatmaps used the same colour scale, from zero to the 99.5th percentile of the finite expression values pooled across the displayed genes and clusters.

In DG, *Calb2* and *Calb1* were used as maturation-related reference markers because calretinin is transiently expressed during an early postmitotic stage of granule-cell differentiation, whereas calbindin is associated with more mature granule cells.^34,35^ I compared the displayed cluster-level expression profiles of *Kcnh7*, *Calb2* and *Calb1* descriptively. Because the expression matrix contains cluster-level trimmed means, this comparison represents alignment among transcriptomic clusters rather than direct within-cell coexpression, birth dating or lineage tracing. The same *Calb1*–*Calb2* relationship was not assumed to represent a neurogenic lineage axis in CA1 or CA3.

#### Allen resource documentation

Source Data Fig. 2 links each of the eight displayed sections to its Allen image and experiment record. It lists the 144 broad-region and 44 Purkinje-layer measurements, the queries that retrieve them and the source of the adult cluster expression values. Original images and Allen-supplied measurements are subject to the Allen Institute Terms of Use (https://alleninstitute.org/legal/terms-of-use).

### Mouse Patch-seq association analysis

The Allen cortical Patch-seq protocol combines electrophysiological recording with transcriptomic profiling of individual neurons.^36^ I used the June 2020 Allen/NeMO mouse Patch-seq release^14^ for transcriptomic counts per million (CPM), cell metadata and transcriptomic-type annotations. The numerical electrophysiology feature table came from the public NSS PatchSeqDataset resource^37^. The Fig. 3 code and source-data package identifies the three input files, their source URLs and their SHA-256 checksums. Expression samples sharing a cell specimen identifier were averaged before taking the natural logarithm of 1 + CPM for *Kcnh7*, *Calb1* and *Calb2*. Electrophysiological records were joined by the same identifier.

The electrophysiological features analysed were short-square threshold current, long-square threshold current, resting membrane potential, input resistance and membrane time constant. Associations were estimated separately within Allen Institute transcriptomic (AIT)-defined types rather than from the pooled cell population. Before within-type analysis, numerical electrophysiological features were retained if at least 90% of the matched cells had measurements and the feature s.d. was at least 10⁻⁶. Cells required complete values across all 41 retained features, nonzero summed expression across the available potassium-channel gene panel and complete type, donor and batch metadata. These steps retained 3,693 of 4,435 matched cells. The minimum of 40 complete cells per type left 3,045 cells across 30 types for each reported association.

*Kcnh7* expression and each electrophysiological feature were residualized against donor identity and transcriptomic batch using ridge regression with one-hot-encoded categorical covariates, an intercept and a ridge penalty of 1.0. The within-type partial-correlation estimate was calculated as the Pearson correlation between the resulting residual vectors. Correlations were recorded as missing when either residual vector had negligible variance.

The analysis was repeated after inclusion of a *Calb1*–*Calb2* expression covariate defined as

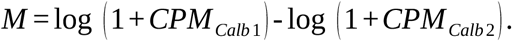

Numeric covariates were standardized within each retained transcriptomic type before residualization. Within-type correlations were Fisher z-transformed and combined using weights of n − 3, where n denotes the number of complete-case cells in each type. The pooled z estimate was the weighted mean, and its standard error was calculated as the inverse square root of the sum of the weights. Ninety-five per cent confidence intervals were calculated on the z scale using ±1.96 standard errors and transformed back to the correlation scale for Fig. 3a.

### Conductance-based Purkinje-cell model

I used the single-compartment Purkinje-cell model of Khaliq, Gouwens and Raman (ModelDB 48332, https://modeldb.science/48332).^15,38^ Its sodium mechanism uses the Raman–Bean kinetic scheme.^39^ I used the three parameter changes that Niculescu et al. introduced to model neonatal mouse Purkinje cells.^4^ They reduced the open-to-blocked-state rate prefactor ε in naRsg from 1.75 to 10⁻¹² ms⁻¹ to suppress resurgent sodium current, reduced maximal sodium conductance from 0.015 to 0.010 S cm⁻² and doubled maximal fast-K conductance (kpkj) from 0.004 to 0.008 S cm⁻² to suppress spontaneous firing. All four conditions shared this baseline before their respective channel manipulations. The compartment had one segment, 20 µm in length and diameter. Baseline mechanisms were naRsg, kpkj, kpkj2, kpkjslow, bkpkj, cadiff, cap, lkpkj and hpkj. ERG3-enabled conditions additionally contained the erg3 mechanism.

The intracellular and extracellular concentrations were Kᵢ = 123 mM, Kₒ = 5 mM, Naᵢ = 9 mM, Naₒ = 149.5 mM, Caᵢ = 0 mM and Caₒ = 2 mM. Sodium and potassium reversal potentials were fixed at E_Na = 60 mV and E_K = −88 mV, respectively. Simulations were performed at 22 °C with a fixed time step of 0.025 ms, CVODE disabled and an initial membrane potential of −65 mV.

The ERG3 component was constrained using native mouse Purkinje-cell ERG-current measurements and rat ERG-family kinetic data.^4,16^ Four conditions were used throughout Fig. 3 and Supplementary Figs. S3-1 and S3-2. Control lacked the ERG3 component. WT-like ERG3 used a maximal conductance density of 0.0006 S cm⁻² and an activation midpoint of −45.1 mV with a 4.2-mV slope factor. MUT-like (V₁/₂ +10 mV) used an approximately +10-mV depolarizing shift in activation half-voltage. High-K increased the maximal conductance density of the native kpkj mechanism from 0.008 to 0.012 S cm⁻² and omitted ERG3. Supplementary Table 1 identifies the experimental sources and the values used during calibration. The implemented steady-state activation midpoints were −45.1 mV (WT) and −35.1 mV (MUT), with the same 4.2-mV slope factor.

The MUT-like shift was selected to approximate the approximately +12-mV positive displacement of the normalized conductance–voltage relation reported for the *KCNH7* p.Arg394His variant.^17^ The reported variant also exhibited slower activation kinetics. Only the activation-voltage shift was represented in this reduced condition; the complete molecular and kinetic phenotype of p.Arg394His was not reconstructed.

### Near-threshold waveform and passive-state measurements

For the near-threshold waveform analysis, a 500-ms square current step was applied after a 300-ms prestimulus period and followed by a 200-ms poststimulus period. The lowest tested current amplitude eliciting a spike under noise-free conditions was selected separately for each model condition. The voltage trace was aligned to its maximum during stimulation and displayed over the interval from 6 ms before to 6 ms after that peak (Fig. 3b).

Passive-state properties were measured under noise-free conditions using hyperpolarizing current steps of −0.002, −0.004, −0.006, −0.008 and −0.010 nA. Each simulation included a 300-ms prestimulus period, a 400-ms current step and a 200-ms poststimulus period. For each step, baseline voltage and steady-state voltage were calculated over the final 50 ms before step onset and the final 50 ms of stimulation, respectively. Input resistance was estimated by linear regression of steady-state voltage deflection against injected current, using non-spiking responses.

For estimation of the membrane time constant, the non-spiking response with the largest absolute steady-state voltage deflection was selected and simulated again at full temporal resolution. Letting t denote time from step onset, the normalized response was defined as

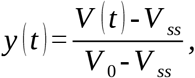

where V₀ and V_ss are the corresponding 50-ms baseline and steady-state means. A linear fit of log y(t) against t was performed within the first 100 ms of stimulation, retaining finite samples with 0.02 < y(t) < 0.98. The time constant was calculated as τ = −1/b, where b is the fitted slope. Fits required at least eight retained samples and a negative slope; otherwise, τ was recorded as missing. Resting membrane potential in Supplementary Fig. S3-1a was the stored 50-ms baseline mean V₀ from this selected response. Input resistance and membrane time constant are shown in Supplementary Fig. S3-1b,c.

### f–I analysis

For the f–I analysis, 23 square-current amplitudes were tested: 0.006–0.020 nA in 0.002-nA increments and 0.024–0.080 nA in 0.004-nA increments. Each simulation comprised a 400-ms prestimulus period, a 1,600-ms current step and a 200-ms poststimulus period. Spikes were detected at upward crossings of −20 mV, and firing rate was calculated over the final 500 ms of stimulation. Ten seeds were used for each condition and current amplitude. Seed identifiers were reused across model conditions and current amplitudes. Fig. 3c shows the across-seed mean ± sample s.d., calculated using Bessel’s correction.

A zero-mean Ornstein–Uhlenbeck current was added throughout each simulation. The process was initialized at zero and updated at 0.025-ms intervals using an exponential decay factor exp(−Δt/τ_n) and independent Gaussian innovations with s.d. σ√[1 − exp(−2Δt/τ_n)]. The stationary s.d. was σ = 0.001 nA and the correlation time was τ_n = 5 ms. These were fixed phenomenological noise settings rather than parameters fitted to measured Purkinje-cell current fluctuations.

### Pre-tonic spike-count repertoire

For the spike-count repertoire analysis, current amplitudes were sampled from 0.006 to 0.020 nA in 0.0005-nA increments and from 0.0205 to 0.0795 nA in 0.001-nA increments. Each simulation included a 200-ms prestimulus period, a 500-ms current step and a 100-ms poststimulus period. The Ornstein–Uhlenbeck noise settings were identical to those used in the f–I analysis. Thirty seeds were used for each condition and current amplitude, with seed identifiers reused across conditions and amplitudes.

Spikes were counted over the complete 500-ms stimulation interval. For each condition and current, the empirical probability of each original integer spike count was calculated across seeds. Counts from 0 to 9 were shown separately in the stacked display, and counts of 10 or greater were pooled into one display category. Modal-class probability, shown by the black line, was calculated before this pooling as the maximum probability among the original integer spike-count classes. Fig. 3d displays 0.006–0.030 nA. Source Data for Fig. 3d (Source_Data_Fig3d.csv) gives the unpooled results for all currents.

### Count- and timing-based square-step analyses

Condition-specific operating currents I₀ were obtained by linear interpolation of the mean final-500-ms f–I curve at an 80-Hz reference rate. Square-step responses were generated at I₀ and I₀ + ΔI, with ΔI ranging from 0.00025 to 0.00225 nA in 0.00025-nA increments. The stimulation and noise protocols were the same as those used for the f–I analysis. Cumulative responses were calculated from stimulus onset over windows of 200, 400, 500, 600, 800, 1,000, 1,200, 1,400 and 1,600 ms.

Ten seed-indexed replicates were generated for each condition and current offset. Baseline and shifted inputs used different random-number streams, with seeds s and s + 100,000, respectively. The same seed scheme was reused across model conditions and current offsets. Different decision windows were evaluated from the same simulated spike trains rather than generated as independent trials.

For count-based analyses, response entropy was calculated in bits from the unpooled spike-count distribution at I₀ + ΔI. Jensen–Shannon divergence was calculated between shifted-input count distributions at adjacent ΔI values. At an interior ΔI value, local separability was defined as the larger of the divergences to its two adjacent values; at an endpoint, the single adjacent divergence was used. Count-based d′ was the shifted-minus-baseline mean count difference divided by the square root of the mean of the two variances. Each variance was calculated with divisor n and floored at 0.25 spikes², corresponding to an s.d. floor of 0.5 spikes.

Timing scores were computed from the same spike trains using an 80-Hz reference frequency. The score grid included vector strength, the Rayleigh-related score N × VS², four linear interspike-interval (ISI) scores and five log-ISI scores. Linear ISI scores used Gaussian widths of 0.15, 0.20, 0.30 and 0.50 times the 12.5-ms reference period; log-ISI scores used widths of 0.35, 0.50, 0.65, 0.85 and 1.00 for log(ISI/12.5 ms). Vector-strength and Rayleigh-related scores were treated as missing for windows containing fewer than three spikes. ISI-based scores were assigned zero when no valid positive ISI was available.

Each of these 11 timing-score definitions was also evaluated after count residualization. Scores from baseline and shifted trials were pooled within each condition, offset and window, regressed on spike count with an intercept, and replaced by the residuals. Where count did not vary or fewer than three valid observations were available, the original scores were retained. For each configuration, Cliff’s delta was calculated as

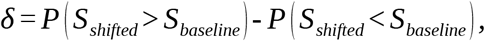

where S denotes the timing score. Timing-based d′ was calculated using the pooled sample variance, with the resulting denominator bounded below by 10⁻⁹.

Pairwise differences were calculated as WT-like ERG3 minus Control, WT-like ERG3 minus High-K, WT-like ERG3 minus MUT-like (V₁/₂ +10 mV), and High-K minus Control. The timing consensus used the 22 timing configurations; a count-only configuration was retained in the full source-data table but excluded from this consensus. For each contrast, current offset and decision window, the median signed difference and fraction of strictly positive differences were calculated separately for Cliff’s delta and timing-based d′ using finite estimates only. The number of contributing configurations was recorded, and summaries with no finite estimates were left missing. Count-based and timing-consensus maps are shown in Supplementary Fig. S3-2a,b.

### Periodic forcing and locking analysis

Condition-specific baseline currents were obtained by interpolation of noise-free current–rate scans to approximate unmodulated firing rates of 80 or 60 Hz. The baseline current was applied from simulation onset until the end of periodic stimulation. Sinusoidal modulation began after 400 ms, lasted 2,500 ms and was followed by a 200-ms poststimulus period. The first 400 ms of modulation was excluded, leaving a 2,100-ms analysis interval. Modulation amplitudes were 0.0005, 0.0010, 0.0015, 0.0020, 0.0030, 0.0040, 0.0060 and 0.0080 nA. Frequencies were sampled in 2-Hz increments over ±20 Hz around each target. The same Ornstein–Uhlenbeck noise settings as in the f–I analysis were used throughout each simulation. Ten seeds were used, with seed identifiers reused across model conditions, amplitudes and frequencies.

For each trial, vector strength was calculated from spike phases at the imposed modulation frequency during the analysis interval. P11 denoted a 1:1 modulation-to-spike relationship, with the target modulation frequency equal to the calibrated base frequency. P21 denoted a 2:1 modulation-to-spike relationship, with the target modulation frequency equal to twice the base frequency. A trial was classified as locked when vector strength was at least 0.5, at least 30 spikes were detected, and the firing-rate-to-modulation-frequency ratio differed from 1.0 for P11 or 0.5 for P21 by no more than 0.15. Locking probability was the fraction of seeds meeting all three criteria.

Target locking probability was evaluated at the exact predefined target-frequency bin. Off-target locking probability was averaged over bins within the same mode-specific ±20-Hz window that lay more than 4 Hz from the target. This definition yielded 16 off-target bins per mode at the 2-Hz spacing. Frequencies from the other locking mode were not included. Target selectivity was defined as target locking probability minus this mode-local off-target average. The target bin and off-target set were required to be present; missing target bins or empty off-target sets were not replaced by nearest-frequency estimates or zeros.

For condition contrasts, seed identifiers were required to be unique and identical between the two conditions and were aligned before analysis. The resampling unit was the seed, retaining its entire frequency profile. Ninety-five per cent confidence intervals for mean target-locking and selectivity differences were estimated from 5,000 seed-paired percentile-bootstrap resamples, using the same resampled seed indices for both conditions.

Two-sided P values were calculated separately from the confidence intervals using within-seed condition-label permutations. Under the null assumption of exchangeability of the two condition labels within each seed, all 2¹⁰ = 1,024 label assignments were enumerated. The test statistic was the mean within-seed condition difference; the P value was the proportion of assignments yielding an absolute statistic at least as large as the observed value. Bootstrap sign-tail fractions were not used for hypothesis testing.

P values were adjusted using the Benjamini–Hochberg procedure separately for each base frequency, locking mode and metric. Each family contained all eight amplitudes and four analysed contrasts: WT-like ERG3 minus Control, WT-like ERG3 minus High-K, WT-like ERG3 minus MUT-like (V₁/₂ +10 mV), and High-K minus Control, giving 32 comparisons per family. The first three contrasts are displayed in Fig. 3f. Adjusted values are reported as q_BH.

A target-centred, FWHM-like bandwidth was additionally calculated within the same mode-specific ±20-Hz window. When target locking probability was at least 0.2, the contiguous region containing the target at or above half the target probability was identified; its boundaries were linearly interpolated between adjacent sampled frequencies. Widths reaching either window boundary were flagged as edge-limited, and widths were recorded as missing when target probability was below 0.2. No interpolation was performed across the unsampled gap between P11 and P21 windows.

### Computational reproducibility

Public input files, simulation configurations, mechanism inventories, analysis scripts and output files were recorded with SHA-256 hashes. Completed simulations were checked for the required grids, seed identifiers, array shapes, finite values and protocol metadata before analysis. Missing or incomplete simulations were not assigned zero responses. Interrupted runs were resumed only from outputs or checkpoints compatible with the recorded configuration and available source fingerprints. Software versions and per-job logs were retained in the acquisition archive. The figure source data include the source-input hashes, mechanism inventory and analysis settings.

### Scope of the reduced adaptation models

Figure 4 used three distinct computational abstractions to examine history-dependent negative feedback. Q-learning used a decaying hidden-activity penalty, the spiking network used a spike-triggered potassium-like conductance, and the RBM used a hidden-state-dependent sampling penalty. Their strengths and time constants were model-specific; they were not fitted to a common ERG3 gating scheme. Measurements of native Purkinje-cell and recombinant rat ERG currents^4,16^ motivated the *KCNH7*-inspired feedback used here. MNIST-based experiments used the canonical 60,000-image training and 10,000-image test partitions.^31^

### GridWorld task and learning protocol

The reinforcement-learning task was a 3 × 4 grid with twelve one-hot-encoded states and four actions: up, down, left and right. Using row-major state indices 0–11, the start was state 8, the wall state 5, the goal state 3 and the penalty state 7. Moves outside the grid or into the wall left the agent in its current state. Entering the goal gave reward +1 and terminated the episode; entering the penalty state gave −1 without termination; other transitions gave zero reward. Episodes were capped at 1,000 actions. The plotted return was the undiscounted sum of episode rewards, not a per-step efficiency score.

The Q-network had architecture 12–100–4, He-scaled Gaussian weights and initially zero biases. Training used one-step semi-gradient updates with learning rate 0.03, discount factor γ = 0.9 and ε-greedy exploration with ε = 0.04. The loss for the chosen action was the squared temporal-difference error, with target r + γ max Q at non-terminal transitions and r at the true goal. Time-limit truncation did not remove the bootstrap term. Elementwise parameter gradients were clipped to [−5, 5]. ReLU hidden responses were capped at 500 and real-valued Q outputs at ±500. The step condition thresholded both hidden and output responses at 0.5 before output transmission; its updates used the explicitly implemented capped-ReLU hidden surrogate and linear-output surrogate, not derivatives of the binary response. The surrogate also did not differentiate through the response caps.

Independent transmission factors were drawn for hidden responses and the four outputs from U(ℓ, 1), with ℓ = 1, 0.66 or 0.60; ℓ = 1 gave unit transmission without random draws. Lower ℓ changed both the variance and the mean gain, and was not a mean-preserving noise manipulation. Matched updates reused the realized hidden and output masks from the current-state temporal-difference forward pass. Mismatched updates used the same forward perturbation mechanism but substituted unit masks in the backward computation. Policy exploration and transmission used separate random-number streams. Exploratory actions were uniformly random and used a noise-free behavioural response for updating adaptation; the target and current-state learning passes retained the specified transmission setting. The current-state temporal-difference pass was a fresh forward computation evaluated at the cached pre-behaviour adaptive state; it was not necessarily the same stochastic realization used to choose the action.

For slow-K conditions, the hidden preactivation was *z_t_* = *x_t_ W* _1_ + *b*_1_ - *a_t_*, and the adaptive state was updated once per behavioural response according to

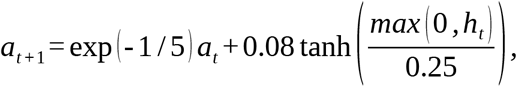

with elementwise operations. The drive was the unmasked hidden response. Adaptation was treated as a stop-gradient state, advanced neither during target computation nor during the fresh current-state update pass, and reset at episode boundaries. Target values used the post-behaviour history state. Slow-K-off conditions did not apply the penalty. Greedy ties were resolved by the first maximal action in the fixed action order.

The full factorial comprised two response types, two slow-K settings and two update-mask settings at each of the three transmission settings, with ten seeds per combination. Each run lasted 50,000 episodes; curves summarize consecutive 500-episode blocks. Corresponding seed numbers reused the initialization procedure, but action and state trajectories could diverge after the manipulations changed behaviour. Frozen evaluations used 200 greedy episodes per trained model with no parameter updating, fresh zero adaptation at each episode and native transmission. Evaluation random-number streams were isolated from training. Mean and SEM were calculated across the ten trained seeds, rather than treating evaluation episodes as independent training replicates.

### Spiking-network construction and training

I used the excitatory–inhibitory network architecture of Diehl and Cook^19^ and implemented the threshold and slow-K comparisons in Brian 2, with Brian2CUDA for training.^40,41^ It contained 784 input units, 400 excitatory conductance-based leaky integrate-and-fire units and 400 inhibitory units. Inputs connected to every excitatory unit. Each excitatory unit drove its corresponding inhibitory unit, which inhibited all excitatory units except its paired source. Static excitatory-to-inhibitory and inhibitory-to-excitatory conductance increments were 10.4 and 17.0 nS, respectively. Excitatory input delays were sampled uniformly from 0 to 10 ms during model construction and retained with their connection indices for evaluation.

Simulations used exponential Euler integration with a 0.5-ms timestep. Excitatory and inhibitory membrane time constants were 100 and 10 ms; excitatory and inhibitory synaptic time constants were 1 and 2 ms. Excitatory cells rested and reset at −65 mV; inhibitory cells rested at −60 mV and reset at −45 mV. Excitatory synaptic reversal was 0 mV, and inhibitory reversal was −100 mV in excitatory cells and −85 mV in inhibitory cells. Conductance variables in the code were dimensionless multipliers of 1 nS. All conductances and the spike timer began at zero, while the initial membrane potential was 40 mV below each population’s resting parameter.

Images were presented in training-set order, wrapping through the 60,000 images for a total of 180,000 presentations. Each presentation comprised 350 ms of input and 150 ms of zero-input rest. Input firing rates were *p_i_ ×* 2 / 8 Hz for original pixel intensities *p_i_* in [0, 255]. Incoming weights were initialized as 0.3 (*U* + 0.01), *U* ∼ *U* (0, 1), and each incoming excitatory weight column was normalized to a sum of 78.0. Column normalization was applied every image cycle, including the initial boundary.

I used a nearest-spike triplet-style rule with pre-, fast post- and slow post-synaptic traces.^42^ Their decay constants in this implementation were 20, 20 and 40 ms. The presynaptic event first transmitted the current weight, set the pre-trace to one and applied depression proportional to the fast post-trace. The postsynaptic event applied potentiation proportional to the pre-trace and the pre-event slow post-trace, then set both post-traces to one. Event-driven updates were clipped to [0, 1]; the subsequent column normalization was a separate operation. The normal depression/potentiation coefficients were (0.0001, 0.01), and the high-rate coefficients were (0.007, 0.7). The high-rate condition therefore scaled both coefficients 70-fold. No class labels were used in these synaptic updates.

### Nominal threshold and slow-K manipulations

The low, normal and high nominal threshold conditions used excitatory/inhibitory threshold parameter pairs of (−62, −44), (−52, −40) and (−42, −35) mV, respectively. These comparisons changed both populations’ nominal thresholds, not the excitatory threshold alone. Excitatory spiking additionally required

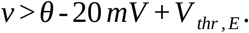

The adaptive threshold θ started at 20 mV, increased by 0.05 mV per spike and decayed with a time constant of 10^7^ ms during training. Thus, “nominal threshold” did not mean that the complete firing threshold remained constant during learning.

Slow-K was varied at the normal nominal excitatory/inhibitory threshold pair. The dimensionless conductance state satisfied *d g_K_* / *dt* =-*g_K_* / ( 200 *ms* ) and increased by 0, 0.02 or 0.05 per excitatory spike for off, weak or strong conditions. The additional current was

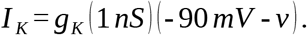

The nominal membrane refractory periods were 5 ms for excitatory and 2 ms for inhibitory cells. A separate excitatory timer, reset to zero at each spike, evolved at 0.1 s/s, and spiking also required timer > 5 ms. Consequently, the timer imposed an approximately 50-ms minimum interval, subject to timestep discretization, in addition to the nominal membrane refractory period. This constraint was shared by all conditions and was not the slow-K mechanism. Neither *g_K_* nor the other dynamical states were reset between images.

Six labelled conditions, two learning rates and three seeds produced 36 separately acquired models. Threshold-normal and slow-K-off had the same nominal model parameters but retained separate acquisition records and were not pooled. Weights and θ were saved every 10,000 presentations. Intermediate checkpoints retained the acquisition’s end-of-boundary-step phase, after the boundary update and normalization; the final checkpoint was the state after completion of the requested run. The recorded phase, end_after_boundary_update, was not relabelled as a pre-update checkpoint.

### Frozen-circuit STDP evaluation and drift measurements

For each of the eighteen checkpoints per model, evaluation reloaded the saved weights, θ, synaptic connection order and realized delays. Synaptic learning and θ adaptation were disabled. A common saved input-event tape contained 100 randomly selected examples per digit from the MNIST training partition for neuronal label assignment, followed by 100 per digit from the test partition for scoring. Events were generated using independent Bernoulli draws at each input unit and timestep with probability equal to its prescribed rate times the timestep; no input events occurred during rest. Image selection and event generation used separate streams derived from evaluation seed 123. The same image order and event tape were used across native and acute slow-K-OFF evaluation, avoiding changes in the stimulus caused by regenerating synaptic delays or input noise.

Each excitatory neuron was assigned the digit eliciting the largest summed spike count in the labelling subset. Classification summed spikes across neurons assigned to each digit; all-silent trials were assigned no class and counted as incorrect. Counts included the full 350-ms stimulus and 150-ms rest cycle. Accuracy and population spike output were computed from the held-out test subset. No membrane-state reset separated the labelling and test portions. Native evaluation retained the model’s trained slow-K setting. Acute OFF evaluation set the slow-K jump to zero while retaining the same trained circuit, weights, θ, input tape and neuron labels fitted in native evaluation. The main figure shows native evaluation; the 648 native/OFF checkpoint pairs were a separate evaluation contrast, not additional independent trained models.

For consecutive checkpoints, weight change was the Euclidean norm of the difference between the full input-weight vectors. Receptive-field drift was computed for corresponding excitatory neurons as

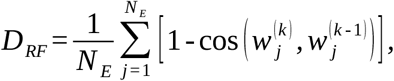

where *w _j_* is the 784-element incoming weight vector. The implementation stabilized squared norms and their product with 10^-12^ and clipped cosine values to [−1, 1]. Neurons were not reassigned or permuted to minimize drift. The first recorded checkpoint had no preceding recorded checkpoint for this comparison and was left undefined, not replaced with zero. These are net differences between saved checkpoints, not cumulative update magnitudes or direct measurements of class-information loss. Curves show means and sample standard deviations across three training seeds.

### Bernoulli RBM and matched adaptive learning

The image generator was a Bernoulli restricted Boltzmann machine^20,21^ with 784 visible and 48 hidden units. Five task sets contained digits {5}, {5,1}, {5,1,3}, {5,1,3,7,9}, or all ten digits, giving D = 1, 2, 3, 5 or 10. D therefore varied both load and the fixed class composition; it was not a manipulation or measurement of representation dimensionality. At most 6,000 training images and exactly 300 test images per included digit were selected. Pixel intensities were scaled to [0, 1] for data observations; unconstrained generated visible states and sampled hidden states were binary.

For each task, 150 independently initialized models were trained at each temperature T = 0.25, 1 or 4, with and without adaptation, giving 4,500 models across 30 conditions. Corresponding replicate identifiers used the same initial weight tensor across temperatures and adaptation conditions. Weights were normally distributed with standard deviation 0.01; both biases and the adaptive state began at zero. Task subsets, cue arrays and phase-specific random-number streams were paired across conditions. Pairing the streams did not imply identical sampled states once model responses diverged.

The hidden conditional probability was

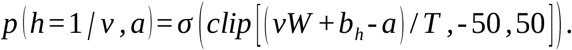

Each transition sampled the hidden state, updated adaptation as *a ’* =exp ( - 1 / 20) *a* + *κh*, and then sampled the visible state from the temperature-scaled visible conditional. The adaptation increment κ was 0 for baseline and 0.05 for adaptation. It depended on sampled binary hidden activity, not explicitly on intermediate probability or semantic ambiguity.

Both arms used persistent contrastive divergence^21^ with one negative-chain transition per update, minibatch size 16, learning rate 0.02 and 400,000 updates. Positive-phase hidden probabilities were computed from data with zero adaptation. Negative visible states and adaptive states persisted between updates, and negative hidden probabilities were recomputed from the terminal visible and adaptive states after the transition. Weights and visible/hidden biases were updated from positive-minus-negative statistics. Temperature scaled both hidden and visible conditional logits; the effective learning rate was held fixed without an additional 1/T multiplier. The adaptive sampler was history dependent, and its updates were not claimed to be an exact equilibrium likelihood gradient. No classifier loss was backpropagated into the generator.

### RBM completion, release and free-run evaluation

A separate scoring classifier was trained once on the complete MNIST training set and then frozen for all generator conditions. It was a 784–256–10 multilayer perceptron with a ReLU hidden layer, trained for three epochs with Adam (learning rate 0.001; batch size 256; seed 1234). The generator itself did not contain ReLU hidden units. The same classifier weights were used across all arms, so reported accuracy describes the classifier’s labels for generated images rather than a supervised generator-training objective.

For D > 1, each test image retained each pixel independently with probability 0.65; other pixels were replaced with uniform [0, 1] noise. Gaussian noise of standard deviation 0.05 was then added and values clipped to [0, 1]. Retained values of this noisy cue were clamped after every visible update during 25 recall transitions; adaptation began at zero. D = 1 instead retained every pixel without corruption, so its completion result is an intact-cue reference rather than a matched difficulty condition. Evaluation used the model’s training temperature. Minimum-class completion accuracy was the smallest per-digit accuracy within each model, measured at the final training checkpoint.

For post-cue evaluation, a separately sampled 25-step recall was followed by 5,000 fully unclamped transitions, continuing both visible and adaptive states. Panel f shows overall terminal classification accuracy relative to the original cue labels; it is neither a lower quartile nor a survival probability. For free-run evaluation, each trained model generated 128 particles from independent Bernoulli(0.5) visible initialization and zero adaptation for 5,000 transitions. The effective in-set class number was

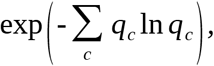

where *q_c_* was the terminal class frequency normalized over the trained digit set. It was set to zero when no generated samples belonged to that set. It summarizes class occupancy among terminal samples, not trajectory transition rate, stored memory capacity or image fidelity. The hidden mid-fraction was the fraction of terminal hidden conditional probabilities in the inclusive interval [0.2, 0.8], averaged over particles and hidden units. Panels e–h show medians and 25th–75th percentile ranges across the 150 trained models.

### Statistical scope and reproducibility

The independent computational replicates were the ten trained seeds for Q-learning, three training seeds for STDP and 150 trained model replicas per RBM condition. Checkpoints, test images, generated particles and native/OFF evaluations were repeated or nested measurements, not additional independent training replicates. The main figure is descriptive; no p-values, equivalence claims or causal-mediation claims were inferred from overlap or separation of its plotted bands. In particular, similar T = 4 medians do not establish equivalence, and similar evaluation spike counts do not establish activity-matched learning.

Final figures were regenerated from saved model-level measurements, with input identities, analysis definitions and panel-to-source mappings retained. STDP reanalysis used saved spike events and weight arrays; native/OFF comparison retained the trained circuit and common input tape. RBM reaggregation retained generated labels and hidden probabilities and checked occupancy counts, distinguishing float32 ratio-rounding differences from changes in underlying counts. These checks establish numerical provenance and reproducible aggregation, not independent biological validation. The figure-source inventory distinguishes redrawing from saved values, reanalysing archived data and training new models.

### Conductance and memory assays for Figure 5

#### Conductance-based reservoir and channel perturbations

The conductance networks in Figs. 5–7 used the same adapted Purkinje-cell membrane mechanisms described for Fig. 3, simulated in NEURON^43^. Each network contained 100 excitatory and 100 inhibitory cells. Integration used a 0.1-ms step, an initial membrane potential of −65 mV and a temperature of 22 °C. Ordered recurrent connections were sampled with probability 0.5. Weights were sampled from a uniform distribution between zero and the assigned cap (0 or 0.001 nS), with 1–10 ms delays and excitatory and inhibitory synaptic time constants of 5 and 10 ms. The tonic current was 0.015 nA per cell. Excitatory cells received an additional 0.03 nA times the positive part of the input, and inhibitory cells received 0.03 nA times the positive part of its negation. The forecast and history protocols used stored Poisson background events with a 10-ms mean interval and weights of 1.5 × 10⁻⁶ nS. Network IDs 1001–1030 paired the circuit and the background across matched conditions.

The reference WT ERG density was 0.0006 S cm⁻². MUT shifted the activation midpoint from −45.1 to −35.1 mV with the other gate equations held fixed. In the accelerated variant, multiplying both activation rate coefficients by ten shortened the activation time constant while preserving steady-state activation and the inactivation gate. The Fast K⁺ series scaled the native kpkj conductance (reference density 0.008 S cm⁻²) with ERG disabled. WT ×0.5 and ×0.577038 were fixed at pilot values selected using A1 training firing rate or current. Residual rates and mean E0 ERG current densities were evaluated during 5–40 s. Native event counts used (start, end] intervals.

#### Input construction and forecasting

For A1 and A2, I discretized the Mackey–Glass delay equation^44^ with a unit time step, giving x[t+1] = x[t] + βx[t−τ]/(1 + x[t−τ]¹⁰) − γx[t], with γ = 0.1, x[0] = 1.2 and zero delayed input before the first available delayed sample. For A1, β = 0.2 and τ = 17 steps. For A2, β = 0.3 and τ = 34 steps. The generator returned the series from x[1] onward, so the first raw sample for both waveforms was 1.08. A3 was a fixed iterative amplitude-adjusted Fourier-transform surrogate of A1, which preserves the sampled amplitude distribution and approximately preserves the spectrum. Each input was normalized using the mean and population standard deviation of its full 1,201-sample waveform and presented in 50-ms steps. The same waveform was used across seeds within an input family.

The float32 readout included a separate bias and 201 features. Two hundred features summarized recent spikes using 50-ms bins and a 50-ms exponential decay, and one feature gave the difference between the mean excitatory and inhibitory features. Initial weights were Gaussian draws scaled by 0.01 and the initial bias was zero. The one-step prediction error updated weights and bias in chronological order at a delta-rule learning rate of 10⁻⁴ during steps 0–799. Steps 800–899 were washout. During test steps 900–1199 (45–60 s), the final weights and bias remained fixed. Test MSE averaged squared error with respect to input samples 901–1200. For the persistence benchmark, the prediction for the next 50-ms sample was the current input value. Supplementary Fig. S5-2 reports model MSE divided by persistence MSE on exactly the same 300 targets. A ratio above one means the model had the larger error. Supplementary Fig. S5-1 shows the original held-out targets and predictions for seed 1001.

#### Reconstruction of the preceding input

Following the delayed-input reconstruction framework^45^, independent uniform inputs on [−1, 1] were sampled in 50-ms steps and shared across conditions within seed. In the main assay, background events were disabled. After a 200-bin wash-in, 201 weighted spike features (200 cell features and their excitatory-minus-inhibitory population mean contrast) or 200 endpoint membrane voltages were used. For a state measured after input bin t, the target at delay d was u[t−d]. The float64 ridge models used an intercept and training-only standardization, with feature standard deviations below 10⁻⁸ replaced by one. One penalty per representation and job was selected from 0.01, 0.1, 1, 10, 100, 1,000 and 10,000 by mean validation MSE across delays 1–60. The chronological inner split used 600 training states and 200 validation states, with scaling derived from the first 600 states. Ties selected the smaller penalty. The model was refitted on all 800 training states and evaluated on the final 360 held-out states.

Predictive R² was 1 − SSE/SST, with SST centred on the held-out target mean. At d = 1 the reported score was the positive part of the observed predictive R² minus the mean positive part across 200 target-permutation refits. In the archived analysis of delays 2–20, the positive parts were first summed across lags separately in each observed and permuted fit, and the mean permuted sum was then subtracted. Training and test target rows were permuted independently, with all lag columns in each row kept together. The training-feature scaling and design decompositions were reused across permutations. Penalty selection and refitting were repeated for each permuted target matrix. Signed test scores and negative adjusted scores were retained in the source data.

Pointwise 95% intervals for the main reconstruction estimates used Student’s t distribution across the 30 network seeds. For paired contrasts, the intervals were computed from within-seed differences in score.

#### Excitatory firing after different input histories

Each history simulation included a 3-s baseline at u = 0, a 1-s prefix and a 2-s suffix, with continuous network state. The displayed prefix levels were 0 and +0.5. These delivered 15 and 30 pA total injected current per excitatory cell, respectively. The externally injected current in each inhibitory cell remained at 15 pA. The shared suffix was 0.25 (22.5 pA total per excitatory cell) for the WT ×1.5 activation comparison, and zero (15 pA) for the comparison of WT ×0.5 with MUT ×1. In a seed-1 A1 pilot without recurrence, WT ×0.5 met the criterion for approximate matching of excitatory and inhibitory firing rates to MUT ×1 (relative root-mean-square discrepancy, 1.18%). Both scales were then held fixed, allowing firing and ERG current to differ under the history protocol. Within a seed, the circuit and the absolute-time background event sequence were shared across prefixes. Native excitatory spikes were counted in 50-ms intervals (start, end] and divided by 100 cells and 0.05 s. The integrated history contrast is the paired difference in spikes per excitatory cell for the +0.5 minus zero prefix. Rate ribbons and reported whole-window intervals are pointwise seed-based t intervals. Network seed is the replicate throughout. Descriptive Fast K⁺ extension contrasts used 5,000 paired seed bootstrap resamples. Original sign-flip tests in the source archive used 100,000 draws and the plus-one correction, with separate Holm families for MSE, memory and other state metrics. The main displayed MSE and reconstruction comparisons and their pointwise intervals are descriptive.

### Receptor expression and timed ERG modulation for Figure 6

#### Patch-seq cohorts and receptor scores

I used the publicly released mouse motor-cortex Patch-seq dataset recorded at room temperature^22^. The source was berenslab/mini-atlas, commit 3bf98a524b8e48aa8647e1b9ae8f2351d9c712c9 (GEO GSE163764). Seven cells with an author-provided exclusion reason were removed from 1,329 records. The remaining 1,322 cells formed the common expression-preprocessing cohort. Analyses with complete fine RNA-type and covariate information comprised 410 excitatory cells from 150 mice and 817 inhibitory cells from 220 mice. Ninety-five cells lacking the required classification were not included as association-analysis cases. The E and I cohorts partly shared mice and were analysed separately. All analysed cells came from the motor-cortex dataset.

Exon and intron counts were added and converted to counts per million using the total over all 42,466 measured genes in each cell. Expression was transformed as log2(1 + CPM). Each gene was standardized across the common 1,322-cell preprocessing cohort using the population standard deviation (ddof = 0). The receptor-family score was the equal-weight mean of standardized expression for the fixed 18 genes *Grm1*, *Grm5*, *Chrm1*, *Chrm3*, *Adra1a*, *Adra1b*, *Adra1d*, *Htr2a*, *Htr2c*, *Hrh1*, *Bdkrb2*, *Tacr1*, *Tacr3*, *Oxtr*, *Avpr1a*, *Cckbr*, *Ednra* and *Ednrb*. The predefined hypothesis set indexes receptor transcript expression. The nine-gene main-panel subset was chosen by the four receptor groups and display scope, not by the strongest or most significant coefficients.

Within each cell class, expression or score values were converted to average tied ranks. The *Kcnh7* and receptor ranks were adjusted for mouse effects, implemented by within-mouse centring, and the saved nuisance design comprising sequencing batch, fine RNA type, mouse age and the log-expression contrast *Calb1* minus *Calb2*. Pearson correlation of the adjusted rank residuals defined the reported partial rank coefficient. Age was constant within mouse and therefore did not provide a separately identifiable within-mouse effect. Additional adjustment for library size and detected-gene count was retained as a sensitivity analysis and was not substituted for the displayed base model.

Uncertainty was estimated with 2,000 prespecified bootstrap draws of mouse IDs from the common 265-mouse preprocessing pool. Both cell-class analyses used the same draws. Gene standardization, family scores, ranks and nuisance projections were recomputed in each draw. Pointwise 95% intervals used the 2.5th and 97.5th percentiles of valid estimates. An interval was withheld if more than 1% of the planned draws were invalid. Valid draws formed the interval calculation. The intervals are pointwise and assume independent, exchangeable sampled mice. The mouse was the biological replicate. E and I associations were estimated separately.

#### Housekeeping-derived gene-set reference

I used the mouse housekeeping-transcript list from HRT Atlas^46^ and counted 3,984 data rows and 3,277 distinct candidate gene names in the downloaded file. After excluding *Kcnh7* and the Gq-panel genes, candidates measured in the M1 dataset were required to have nonzero expression in at least 80% of cells in both classes and nonzero variance. This yielded 568 candidates. Within each class, candidates were assigned to quintiles of mean logCPM and ranked by variance within those bins. The 100 genes with the lowest average E/I within-bin variance rank were retained, with gene name resolving ties. *Kcnh7* correlations did not enter selection. Using PCG64 seed 20261022, 1,000 sets of 18 distinct genes were sampled from the fixed 100-gene pool for each class. Each set was scored and correlated with *Kcnh7* using the same common gene-Z reference and base partial-rank estimator as the Gq score. Exact memberships are retained in the source data.

The median and central 95% range across gene-set coefficients are descriptive. The HRT-derived candidates were selected for high detection and stable expression in this dataset. Set size was matched by design. Mean expression, detection, variance and internal covariance can differ between sets. The overlapping sets yield a descriptive gene-set reference for the observed expression matrix.

#### Network inputs and learned readout

The simulations reused the 200-cell conductance-based model described for Fig. 5, with 100 excitatory and 100 inhibitory cells, a 0.1-ms integration step and a 50-ms input and feature-sampling interval. Recurrence OFF set both recurrent conductance caps to zero. ON set both caps to 0.001 nS. The original membrane mechanisms, connection construction and paired saved background-event protocol were retained. ERG maximal conductance was present from initialization at 0.0006 S cm⁻² multiplied by 0.5, 0.577038, 1, 1.5, 2.5 or 3. The ×0.577038 WT scale was selected using A1, recurrence OFF and network seed 1. Two successive bracketed interpolations gave a WT scale at which the mean outward ERG current density reconstructed for excitatory cell 0 from 5 to 40 s exceeded the MUT ×1 value by 1.0% (tolerance, ±2%). Forecast MSE was excluded from the calibration. The resulting scale was retained across the grid, where the currents could differ. The activation midpoint was −45.1 mV at baseline. During an intervention it was set to −35.1 mV in all 200 cells, while maximal conductance and other gate equations were unchanged. Midpoint onset and offset did not directly reset voltage, gates or synaptic state.

A1 was generated with Mackey–Glass parameters β = 0.2, γ = 0.1, exponent 10 and delay 17. A2 used β = 0.3 and delay 34, with the other parameters retained. A3 was an iterative amplitude-adjusted Fourier-transform surrogate of A1, preserving the sampled amplitude distribution and approximately its Fourier magnitudes while changing temporal organization. Stage 1 used the three saved raw input arrays, each containing 1,201 values. Input normalization used the mean and population standard deviation of raw samples [0, 801), including the last supervised training target, with no clipping or fitting to later samples. Normalized values were stored as float32. These parameters were fixed for washout and held-out testing. Figure 5 used full-array normalization, including for the raw A2 waveform shared with Stage 1 of Fig. 6. I therefore interpret the Fig. 6 intervention effects relative to their own unshifted baselines.

Training, washout and testing comprised 800, 100 and 300 input steps, respectively. The feature vector contained one exponentially weighted spike feature per cell over the preceding 50-ms bin, with a 50-ms decay constant, plus the difference between the excitatory and inhibitory feature means. A linear readout with a separate bias predicted the next normalized input. Initial weights were Gaussian draws scaled by 0.01 and the initial bias was zero. The chronological delta update was based on the next-target residual. The weight increment was the residual times the feature vector and a learning rate of 10⁻⁴. The bias increment was the residual times the same learning rate. Weights and features followed the saved float32 implementation. Updates occurred only during training. Final weights and bias were held fixed during testing. The target for test feature indices [900, 1200) was input indices [901, 1201), corresponding to one-step prediction over 45–60 s. Persistence predicted the next sample from the current sample.

#### Activation-shift schedules and endpoints

All schedule intervals were left-closed and right-open. The eight finite pulses used two durations (1.5 and 4.5 s) at each of four onsets. Onsets at 5 and 30 s marked early and late training, and onsets at 45 and 52 s marked early and late testing. Sustained interventions occupied training (0–40 s), testing (45–60 s), or both intervals. The training-plus-test condition retained an unmodulated 40–45-s washout. Together with the no-shift baseline, these gave 12 schedules. Three input families, two recurrence settings, six conductance factors, 12 schedules and ten network seeds produced 4,320 conditions per stage.

For each seed, an intervention effect was MSE minus the unshifted baseline MSE matched for stage, input, conductance and recurrence. Test-pulse windows comprised the pulse itself, an equally long post-offset interval clipped to the test endpoint, and the common final interval 57.5–60 s. For the late 4.5-s pulse, the post-offset and common-final windows overlap. Each window was summarized separately within seed. The training-pulse analysis used final readout displacement relative to the matched unperturbed final weights. The plotted horizontal coordinate is the mean of seed-specific normalized L2 distances. The relative MSE coordinate uses a ratio of condition means. The bias is excluded from the distance and retained separately. All 1,440 seed-level training-pulse pairs contributed to 144 complete ten-seed means. Three unavailable weight pairs were recomputed under the original conditions and verified against saved controls.

#### Independent-input extension and statistical summaries

Stage 2 repeated the complete grid with ten independently constructed realizations per input family paired one-to-one with network seeds 1–10. Within a pair, the same input array was reused across doses, schedules and recurrence settings. Mackey–Glass initial values were sampled uniformly from 1.08 to 1.32. After 10,000 generator updates, a 1,201-point window was taken at an additional uniformly sampled integer offset in [0, 20000). A1 and A2 shared the initial-value and offset draws within each pair. The delayed generator prehistory was zero. Generator burn-in preceded neural simulation. A3 was generated from the paired new A1 series using the iterative amplitude-adjusted Fourier transform algorithm^47^, stopping at an exact repeat or 1,000 iterations. PCG64/SeedSequence namespaces separated initial-value, time-origin and surrogate draws. Their entropy vectors are retained in the frozen manifest. No input was selected or resampled on the basis of neural outcomes or favourable input diagnostics. Normalization and readout training were repeated separately for each new realization. Stage 2 tested the full training protocol on a new input for each circuit. Stage 1 used the original generator output from its initial condition, whereas Stage 2 used windows after generator burn-in. The cross-stage comparison therefore also includes a change in temporal origin for the Mackey–Glass series. Input and network variability are coupled by the paired design. Independent random origins do not guarantee distinct realized shapes, especially for the periodic A2 family.

Means and all ten paired values were retained. Pointwise 95% intervals for computational effects used Student’s t distribution with nine degrees of freedom. The source tables also contain Bonferroni intervals for their named families. Two-sided exact sign-flip tests enumerated all 1,024 sign assignments of ten paired effects, with Holm adjustment separately within each prespecified family. Within a stage, the families comprise 396 intervention-versus-own-baseline effects, 216 absolute ON-minus-OFF comparisons and 198 recurrence-by-intervention contrasts. The cross-stage families comprise 432 absolute MSE differences, 396 intervention-effect differences and 198 changes in recurrence-by-intervention contrasts. These tests require independent, sign-symmetric network-level differences under the null. The two-sided minimum attainable p value is 2/1,024, above the first Holm threshold at α = 0.05 in every listed family. The tests cannot establish equivalence. Panel g displays all three sustained schedules at ERG ×3 as a post-analysis descriptive selection. Full-grid sign concordance summarizes correlated condition means. The cross-stage primary endpoint is whole-test MSE. Panels e and f describe Stage 1 endpoints.

### Fixed readout perturbation assays for Figure 7

#### Reservoir input and frozen readouts

The Fig. 7 reservoir retained 100 excitatory and 100 inhibitory cells, 0.1-ms integration and 50-ms input and feature bins. Its fixed parameter set used connection probability 0.45, excitatory and inhibitory synaptic decay constants of 5 and 8 ms, tonic current 0.014 nA and input-current scale 0.026 nA. A common Poisson event train had a 12-ms mean interval and a background synaptic weight of 1.125 × 10⁻⁶ nS per cell. Recurrence was ON with a 0.001-nS cap. A1 and A2 were fixed deterministic Mackey–Glass series with (β, γ, exponent, delay, initial value) equal to (0.2, 0.1, 10, 17, 1.2) and (0.3, 0.1, 10, 34, 1.2), respectively. The input path and paired background-noise realization were retained for each matched comparison. Low and high contexts used stationary zero-mean OU current with standard deviations of 0.002 and 0.006 nA, a 10-ms time constant and 1-ms sampling. The low and high labels refer to current-noise amplitude.

The protocol comprised 80 pre-training steps, 300 training steps, 50 washout steps and 150 test steps. With 50-ms bins, training occupied 4–19 s and testing began at 21.5 s. Raw input samples [80, 381), including the last training target, supplied the mean and population standard deviation for normalization. The saved normalized vector was float32. Readout features were the 200 per-cell exponentially weighted spike features and their E-minus-I mean contrast. The 200 spike features summed exponentially weighted spikes within each 50-ms bin (50-ms decay). The readout used 300 chronological delta updates at learning rate 10⁻⁴, a separate bias and no weight decay. Once learned on Normal transmission, the final weights and bias were kept fixed when evaluating the corresponding Normal and perturbed states. Different channel settings in a and different ERG settings in b retained separately learned readouts trained under their own Normal conditions. In c–f, all arms within an input and seed used the same readout trained with native ERG ×1.5.

#### Whole-trajectory recurrent-transmission perturbation

Panel a used ERG off and native-kinetic ERG factors 0.5, 1.5 and 3. The Fast K⁺ factors were 1.125, 1.5, 2 and 3, and the Slow K⁺ factors were 2, 4, 8 and 16. Fast/Slow dose series in this panel were in the ERG-off background. Factors multiply each channel’s own native conductance. Each condition was tested with A1 and A2 at low and high OU noise across ten paired network seeds.

Normal recurrent arrivals retained their nominal synaptic increments. The Uniform arm multiplied each increment by an independent nonnegative uniform random value with mean 1 and standard deviation 0.55, generated per connection and arrival. The multiplier varied event amplitude throughout the perturbed circuit trajectory, including the training interval. The evaluated readout came from the matched Normal trajectory. For each network, the waveform endpoint was the square root of the mean squared difference between the Uniform and Normal prediction traces over the test samples. The plotted mean averages network-level RMS values. Normal MSE is the mean squared difference between the Normal prediction and the target. The main-panel RMS is expressed in prediction units. Silence denotes no test spikes in either trajectory and is reported independently of prediction error.

#### Acute transmission perturbations

For panel b, both ERG off and native ERG ×1.5 followed the matched Normal history until test onset at 21,500 ms. The intervention changed recurrent arrivals at or after that time without resetting existing synaptic conductance or cellular state. Private-bin fluctuations used independent edge-specific draws in each 1-ms arrival bin. Repeated arrivals on the same edge within a bin shared its draw. Shared-bin fluctuations used a common multiplier across edges within that bin. Both distributions had mean 1 and standard deviation 0.55. For the E- or I-presynaptic bias condition, targeted edges used mean 1.4 and standard deviation the square root of (0.55² − 0.4²), while nontargeted edges retained mean 1 and standard deviation 0.55. Thus, the nominal marginal RMS deviation from multiplier 1 was 0.55 in each population. Realized arrival-weighted moments can differ because endogenous arrivals sample bins unevenly. The bias condition changes both mean and variance. Block set the post-onset recurrent-arrival multiplier to zero. Previously delivered conductance continued to decay and background drive was retained.

Identical input, graph, noise and saved pre-intervention feature histories were checked against the corresponding clean reference. The primary acute comparison used test bins [20, 40), 1–2 s after onset. MSE differences were calculated against each ERG setting’s own Normal error over those same targets. Waveform RMS used the difference from its own Normal prediction over the same bins. Comparisons between ERG settings therefore concern trained-system sensitivity. The whole-trajectory Uniform and acute perturbation assays were analysed separately. Let y be the target, p the unperturbed prediction and δ the perturbed prediction minus p. Over the same samples, waveform RMS = √mean(δ²) and ΔMSE = mean(δ²) + 2 mean[(p – y)δ]. The cross-term gives the alignment of the prediction change with the original error.

#### Population drive and conductance interventions

Population-drive experiments began from the native-ERG ×1.5 reference, with recurrent weights unchanged. A current of −20 or +10 pA per cell was applied to all 100 cells in either the E or I population from test-relative time 0 to 2 s. Within each input, seed and challenge, arms shared the first second of the challenged trajectory and then branched into continued DC alone or a conductance change in the same targeted population. For negative drive, the intervention set ERG, Fast K⁺ or Slow K⁺ to ×0.75, ×0.5 or ×0.5. For positive drive the settings were ×2.25, ×1.25 or ×8. The two DC polarities had unequal magnitudes, and these conductance choices were not matched for actual current or firing rate. Fast and Slow K⁺ arms retained ERG ×1.5. Both DC and altered conductance ended at 2 s, followed by observation to 4 s. Restoring conductance parameters did not directly reset the network state.

The early, active and post-release windows were [0, 1), [1, 2) and [2, 4) s after test onset, corresponding to bins [0, 20), [20, 40) and [40, 80). Main panels c–e use active-window absolute MSE, and the comparison to continued DC alone is paired within seed. A negative intervention-minus-DC MSE difference denotes lower error during continued drive. The intact reference was the prediction of the paired, undriven circuit for the same input, seed, target samples and time window. Panel c compares direct current applied to the E or I population. Panels d and e show conductance changes in I cells, and f shows the corresponding +10-pA A2 example for seed 1, selected by index.

#### Replication uncertainty and retained outcomes

All ten network seeds were retained, including adverse and silent outcomes. Existing pilot seeds 1–3 were preserved, and seeds 4–10 supplied the additional matched observations. Inputs were fixed within each Fig. 7 experiment. The network seed was the unit of replication. The source tables report the pilot, the seven additional seeds and the pooled sample separately. The combined sample includes the hypothesis-forming pilot.

For acute comparisons, source tables report all paired values, raw positive/negative/zero sign counts, mean paired differences and pointwise unadjusted 95% Student-t intervals. The ten-seed intervals use nine degrees of freedom and assume approximately normal paired differences across independent network seeds. These are pointwise descriptive intervals for panels b–e. The original whole-trajectory analysis used one exploratory exact sign-flip comparison. Each seed contributes the equally weighted mean of its four input-by-noise ERG ×1.5-minus-off RMS contrasts, and all 1,024 sign assignments are enumerated. The asterisks in panel a mark silence. Undefined normalized endpoints remain missing when their denominators are zero. The main RMS endpoints are unnormalized.

### Receptor signal history and value updating for Figure 8

#### Receptor-linked gating and state retention

Two continuous rate-model architectures were used: competing populations for state maintenance in Fig. 8a,b, and an E/I circuit with a cue-dependent readout in Fig. 8f,g. Their activity variables are dimensionless and are not calibrated firing rates in hertz. The receptor-to-ERG3 link is an explicit model assumption. Transcript associations and the published orexin experiments provide separate biological context; neither establishes OX2R- or mGluR5-mediated ERG3 current in the modeled populations.

The effective receptor signal q shifted the activation midpoint according to V₁/₂(q) = −45.1 + 10q mV. Activation approached m∞ = σ[(V − V₁/₂)/4.2], where σ denotes the logistic function. Current was proportional to ḡ3mh(V − E_K), with E_K = −88 mV and ḡ3 = 0.0009 S cm⁻². The voltage proxy was V = −58 + 24r mV, using P1 activity in the competing-population model or the targeted E or I activity in the task model. The native voltage-dependent activation and inactivation time constants and h∞ were retained from the archived ERG implementation. q shifted m∞ only: it did not shift the voltage arguments of the time constants or h∞, and did not change ḡ3. Signal-filter time constants were varied separately from channel-gate kinetics.

For reproducibility, the native gate functions used A = σ[(V − 114.224)/30.612], B = σ[−(V + 194.405)/31.002], A_h = σ[(V − 123.983)/35.132] and B_h = σ[−(V + 88.914)/25.716]. In seconds, τ_m = 0.001/{[0.4 + 1.2σ((−43.168 − V)/19.186)](A + B)}, τ_h = 0.001/{[0.55 + 0.15σ((V + 62.5805)/38.8568)](A_h + B_h)}, and h∞ = 1 − A_h/(A_h + B_h). Gate updates used the exact first-order exponential relaxation over each 5-ms step. A reference activation gate followed the same voltage trajectory with its activation midpoint fixed at −35.1 mV (q = 1) for the state-maintenance assays and −40.1 mV (q = 0.5) for the task assays. The difference between the evolving active and reference currents was converted into a change in rate-model drive. Even an intact, time-varying cue command could differ from the tonic reference in the task assay. Current-to-drive conversion was a reference calibration, not a cell-type physiological fit.

The two rates obeyed τ_r dr₁/dt = σ(5u₁) − r₁ and τ_r dr₂/dt = σ(5u₂) − r₂, with τ_r = 0.1 s. Their drives were u₁ = a₁r₁ − b₁₂r₂ + d₁ + g + Δu_K + ξ₁ and u₂ = a₂r₂ − b₂₁r₁ + d₂ + ξ₂. Nominal self-excitation was 1.2, cross-inhibition 1.6 and tonic support g = 0.5. Each circuit received independent fixed ±5% uniform variation in these coefficients, a P1 drive offset within ±0.01, and d₂ = −0.25 with an offset within ±0.03. Twelve circuit seeds (101–112) were paired across conditions. Four independent stationary OU noise sequences per circuit were shared across the compared arms; OU correlation time was 0.2 s. Rates were integrated by Euler updates with a 5-ms time step.

For sustained-signal conditions, the current-to-drive conversion equated the full q = 1 to q = 0 steady current difference at the intact P1 reference voltage to that circuit’s g. The compensation arm added the corresponding constant drive. Each condition underwent 30 s of deterministic settling from the calibrated intact P1 starting point. A valid initial state required r₁ − r₂ ≥ 0.5 and a maximum final-step increment below 10⁻⁹. Failure to establish this state was retained in the planned denominator and was not scored as a subsequent exit. Initially valid conditions were followed for 120 s. The full saved drive grid ranged from −0.55 to −0.85 and included noise s.d. 0, 0.05 and 0.075; panel a uses 0.075. P2 entry required r₁ − r₂ ≤ −0.5 for 0.5 s continuously. First-entry time was the confirmation time, rather than the beginning of that run. Supplementary Table 2 reports establishment, failed starts and subsequent exits for the three ERG control conditions at drive −0.65. This first-entry endpoint was evaluated once per trial.

For panel b, the retained comparison used drive −0.85, receptor efficacy 0.75, stationary OU noise s.d. 0.05 and a 120-s horizon. Six 3-s interruptions formed two triplets. Within-triplet recovery intervals were 1 or 7 s, with paired jitter of ±0.35 s. First onset was sampled between 32 and 35 s, and the last onset occurred 49–51 s after the first. First and last onsets, circuit parameters and noise were held across the paired schedules. q relaxed toward the commanded efficacy with a 10-s time constant. For each circuit–noise pair, the flat control extended from the first clustered gap onset to the final clustered gap end. Its depth matched the clustered arm’s integrated filtered q deficit through 120 s by trapezoidal integration. The faster-recovery control retained the clustered commands and 10-s loss kinetics but used 2-s recovery when input returned. It was not area-matched. Baseline eligibility required P1 occupancy of at least 95% during 20–30 s and no confirmed P2 entry before 30 s; planned and common-eligible denominators were retained. Broader drive, efficacy, interval and time-constant conditions remain in the accompanying source tables.

#### Transcript detection and sleep data

I analysed complete-case Patch-seq data from primary motor cortex (410 excitatory and 817 inhibitory cells, Smart-seq2)^22^ and primary visual cortex (4,270 GABAergic interneurons, SMART-Seq v4)^14^. The ventral posterior hypothalamus dataset used 10x Chromium single-cell RNA sequencing. I analysed 185 cells from histaminergic cluster 17 across four pooled libraries^23^. I used the saved cortical CPM-based partial-rank estimates with the original cohort-specific adjustment design described for the cortical expression analysis. For the TMN, I assigned average ranks to the archived normalized expression of each receptor and *Kcnh7*, regressed each rank vector on library indicators and log(1 + total UMI count), and correlated the residuals. An additional common-design sensitivity residualized each gene’s ranks against technical unit and measured sequencing depth within each cohort. Detection was a positive expression value. Conditional co-detection used the number of cells with detectable *Kcnh7* as its denominator. The TMN joint-detection counts also use all sampled histaminergic cells as a separate denominator. Comparisons between cohorts are descriptive.

Mouse-level values were extracted from the public source workbook^8^, without reprocessing raw photometry or sleep recordings. Panel d preserves the authors’ normalized calcium z scores for six mice: sheet Fig.1F–2, S.Fig1F-2, cells B4:E9. The transition-NREM measure is the final 30 s before NREM-to-REM transition. Panel e uses the recorded percentage of NREM bouts followed by REM and subtracts each mouse’s baseline from its intervention value. Inhibition in peptide-expressing orexin-tTA mice and peptide-null orexin-FlpKI/KI mice used ZT7–8; the respective source sheets are Fig. 6E, F, G,H and Fig. 6M,N,O,P, rows 4–9, columns O/P. Intermittent activation used a separate four-mouse cohort at ZT6–7, with four 8-s, 20-Hz trains per minute for 1 h; source values are S.Fig 10F-H, rows 4–7, columns V/W. The displayed summaries are arithmetic means of within-mouse changes, with all mice shown. They are not a formal genotype-by-intervention interaction test. These endpoints concern NREM-to-REM transitions and sending-neuron activity, rather than direct postsynaptic ERG3 measurements. BioStudies S-EPMC10576136.

#### Cue-timed loss and value updating

The task circuit contained two cue-selective excitatory populations E₁ and E₂ and one inhibitory population I, with common excitatory activity E = (E₁ + E₂)/2. Its excitatory transfer function was F_E(u) = {σ[5(u − 0.9)] + σ[5(u + 0.9)]}/2 and its inhibitory transfer function was σ(u). Common drives were u_E = w_EE(E − 0.5) − 2.5(I − 0.5) and u_I = 3(E − 0.5) − (I − 0.5), before cue, noise, challenge and channel terms. Rate time constants were 0.05 s for E and 0.03 s for I, and integration used exponential relaxation over 5-ms steps. Recurrent gains 4.25, 4.5, 4.75 and 5 were each evaluated in eight circuits (seeds 2101–2108), with the same circuit-specific uniform jitter in [−0.05, 0.05] added across the gain grid. All initial rates were 0.5.

Opposite cues drove excitatory subpopulations with opposite signs. Each signed cue command (amplitude 0.32, duration 160 ms) was filtered with a 240-ms time constant. Each population also received an independent Ornstein–Uhlenbeck input initialised at zero (time constant 80 ms, stationary s.d. 0.10). Gaussian measurement noise (s.d. 0.008) was added to the cue features. I used the voltage-dependent relative ERG3 current in the excitatory input equation. The voltage proxy used the mean of the two excitatory populations. Paired conditions shared circuit weights, cue labels, dynamic and measurement noise, and initial values. Panel f used one archived linear decoder per circuit. Each was fitted at gain 4.25 and held fixed across all four nominal gains and compared signal-loss conditions. Decoder inputs were the mean rates of the three populations 200–550 ms after the cue. Cue MSE was scored against cue labels of ±1 and sign accuracy was scored separately, using five late cues per trajectory.

The intact cue-modulated command redistributed gating toward cue onset at a mean q of 0.50 during the intervention epoch. In panel f, the specified q waveform set the instantaneous target activation midpoint directly; the channel gates then relaxed with their voltage-dependent time constants. The shallow condition removed 0.10 q for 300 ms, beginning either at cue onset or 450 ms beforehand, at each of seven cues. The matched continuous condition removed 0.21/13 q from 3 to 16 s. The clustered condition set q to zero from one of five prespecified onsets (3, 5.5, 8, 10.5 or 13 s), with the final numerical step scaled to attain the desired deficit integral (0.21 or 1.30 q·s). I evaluated 24 paired cue/noise trajectories per circuit for each schedule and onset. Persistent low activity required mean excitatory activity below 0.2 for at least 80% of the late 11–16 s window, a value below 0.2 at 16 s and a value below 0.2 at a second endpoint 4 s after task input, noise and differential ERG3 current were removed.

For the value-updating assay in Fig. 8f, each circuit contributed 512 resampled sessions. Starting values came from intact acquisition. The same cue labels, score-sample indices, imposed actions and binary rewards were used across conditions. Rewards followed the reversed rule with probabilities 0.95 for the new-rule action and 0.05 for the alternative. A positive score calibration transformed decoder output into a soft estimate of cue identity. The cue-conditioned action values were updated after each imposed trial with the chosen-action reward prediction error weighted by cue probabilities and learning rate 0.24, clipping stored values to [0,1]. Value-reversal time was the fifth trial of the first five consecutive post-update trials on which the new rule had higher latent value. I assigned 100 trials if the criterion was not reached. The primary curves and comparisons used pointwise paired t intervals across eight circuit means. For f, the displayed mean timing contrasts first average the four gain results within each circuit.

#### Comparison of true reversal and stable adverse feedback

For g, I used archived intact and selective-loss cue scores from the task circuit at recurrent gain 4.25. Each component received five events at 50-ms intervals every 0.5 s. The fast component had no imposed release delay or clearance filter. The slow component had a 1-s delay followed by first-order filtering with a 1-s time constant. Both passed through a 0.25-s downstream filter. The 1-s delay is exactly two cycles of the periodic stimulus. Thus, for the intact periodic signal, the slow component differs in waveform through filtering, while this delay alone leaves its phase unchanged. The intact command was their equally weighted sum. Removing the fast component from the outset reduced the mean command from 0.50 to 0.25 during the 2–3 s reference interval. A fixed decoder generated the scores for each circuit. Cue samples from two independently generated cohorts were pooled for calibration, and two further cohorts were pooled for evaluation. For each circuit and model, a positive score scale was calibrated in independent acquisition sessions against a target of 72% correct over the last 40 of 120 trials. The main replay uses the same starting action values for both conditions, obtained from intact acquisition, and retains each model’s calibrated score scale. There was no new circuit simulation or decoder fitting for the replay.

The learner has one action-value vector for each of the two cues. The score-derived cue probabilities give expected action values as a weighted average of the two vectors. After the forced first old-rule choice, actions are sampled with probability of action 1 equal to the logistic of seven times the difference of its expected value from that of action 0. The selected action values within each cue are updated by 0.24 × cue probability × (reward − expected value of the chosen action), with projection to [0,1]. Resampled sessions share cue labels, sample indices and choice and reward random numbers across matched arms. Both true-reversal and stable-omission arms impose the old-rule choice and no reward on trial 1, with identical updated values within model across those environments. A stable sham earns the expected reward on trial 1. Afterward, the reward rule either reverses or remains stable. The principal figure uses deterministic reward. The sensitivity grid also uses correct-action reward probabilities of 0.95 and 0.75 (incorrect-action probabilities 0.05 and 0.25), two starting-value conventions and clear versus mixed cue scores (25% opposite-cue contribution in the mixed setting).

The old-rule value margin averages across cues the value of the old-rule action minus that of the new-rule action. The time to value reversal is the fifth trial of the first run of five consecutive post-update trials with a negative margin. Sessions that failed to reach the criterion were assigned 100 trials. In panel g, trial 0 denotes the initial margin and trial 1 follows the first forced omission. The action-based measure uses at least eight new-rule choices within ten successive free choices. The model × environment interaction used the circuit-level mean margin across trials 1–20, comparing loss minus intact in true reversal against the same difference under stable omission. The unit for intervals was the paired circuit (eight circuits, 512 simulated sessions each). Confidence intervals in the replays are nominal and pointwise.

## Data availability

This study used public datasets and computational simulations. Developmental expression records are available from the Allen Developing Mouse Brain Atlas and, for P56, also from the Allen Mouse Brain Atlas. Source Data Fig. 2 provides selected Allen measurements, image identifiers, source links, API queries and the author’s summaries. It identifies the inputs from the Allen mouse isocortex and hippocampal formation taxonomy (mouse_ctx-hpf_10x). Cortical Patch-seq data are available from the Allen Institute and NeMO Archive and, for motor cortex, from GEO (GSE163764). Ventral posterior hypothalamus expression data are available from GEO (GSE146692). The Ito et al. workbook is archived at BioStudies (S-EPMC10576136). The digit-recognition models used MNIST. The mouse housekeeping-transcript list is from HRT Atlas v1.0 (Housekeeping_GenesMouse.csv, HRT_Atlas repository, revision 3690f04). Source Data for Fig. 3d (Source_Data_Fig3d.csv) provides the spike-count distributions. The review repository lists source identifiers, figure-level measurements and access instructions for the underlying data. No new animal or human participant data were collected.

## Code availability

The corresponding author can provide author-controlled code and source data to editors and reviewers for confidential evaluation through the editorial office. A versioned public release of these materials is planned at journal publication through the project repository (https://github.com/BreamResearch/Neural-Maturation) and a persistent archive. The original Purkinje-cell implementation is available from ModelDB (https://modeldb.science/48332). Peter U. Diehl’s original STDP implementation, on which earlier project notebooks were based, is available at https://github.com/peter-u-diehl/stdp-mnist. Third-party materials retain their original terms.

## Acknowledgements

I thank Junichiro Yoshimoto for helpful discussions and comments on this work. I also thank Hiroshi Watanabe, Yukihiro Nakamura, and Atsuhito Toyomaki for comments and support. I used ChatGPT Pro (OpenAI) to develop the Discussion, edit the English text and assist with coding. I used the latest version available to me at each stage, including GPT-6.2-Astra at manuscript completion. The author reviewed, edited, and takes full responsibility for all analyses, interpretations, and final manuscript text.

## Author contributions

H.W. designed the study, performed the analyses and simulations, interpreted the results and wrote the manuscript.

## References

1. Rumelhart, D. E., Hinton, G. E. & Williams, R. J. Learning representations by back-propagating errors. Nature 323, 533–536 (1986). 10.1038/323533a0

2. Maass, W., Natschläger, T. & Markram, H. Real-time computing without stable states: a new framework for neural computation based on perturbations. Neural Comput 14, 2531–2560 (2002). 10.1162/089976602760407955

3. Taschenberger, H. & von Gersdorff, H. Fine-tuning an auditory synapse for speed and fidelity: developmental changes in presynaptic waveform, EPSC kinetics and synaptic plasticity. J. Neurosci. 20, 9162–9173 (2000).

4. Niculescu, D., Hirdes, W., Hornig, S., Pongs, O. & Schwarz, J. R. Erg potassium currents of neonatal mouse Purkinje cells exhibit fast gating kinetics and are inhibited by mGluR1 activation. J. Neurosci. 33, 16729–16740 (2013).

5. Schwarz, J. R. et al. Purkinje cell hyperexcitability and depressive-like behavior in mice lacking erg3 (ether-à-go-go–related gene) K⁺ channel subunits. Sci. Adv. 10, eadn6836 (2024).

6. Xiao, K. et al. ERG3 potassium channel-mediated suppression of neuronal intrinsic excitability and prevention of seizure generation in mice. J. Physiol. 596, 4729–4752 (2018).

7. Mochizuki, T., et al. Orexin receptor 2 expression in the posterior hypothalamus rescues sleepiness in narcoleptic mice. Proc. Natl Acad. Sci. USA 108, 4471–4476 (2011). 10.1073/pnas.1012456108

8. Ito, H., et al. Deficiency of orexin signaling during sleep is involved in abnormal REM sleep architecture in narcolepsy. Proc. Natl Acad. Sci. USA 120, e2301951120 (2023). 10.1073/pnas.2301951120

9. Sidiropoulou, K. et al. Dopamine modulates an mGluR5-mediated depolarization underlying prefrontal persistent activity. Nat Neurosci 12, 190–199 (2009). 10.1038/nn.2245

10. Thompson, C. L. et al. A high-resolution spatiotemporal atlas of gene expression of the developing mouse brain. Neuron 83, 309–323 (2014). 10.1016/j.neuron.2014.05.033

11. Allen Institute for Brain Science. Developing Mouse Brain Atlas [dataset]. Allen Brain Map. https://developingmouse.brain-map.org/

12. Yao, Z. et al. A taxonomy of transcriptomic cell types across the isocortex and hippocampal formation. Cell 184, 3222–3241.e26 (2021). 10.1016/j.cell.2021.04.021

13. Allen Institute for Brain Science. Mouse Isocortex & Hippocampal Formation Taxonomy, mouse_ctx-hpf_10x [dataset]. Allen Brain Map. https://brain-map.org/our-research/cell-types-taxonomies/cell-types-database-rna-seq-data/mouse-whole-cortex-and-hippocampus-10x

14. Gouwens, N. W. et al. Integrated morphoelectric and transcriptomic classification of cortical GABAergic cells. Cell 183, 935 –953.e19 (2020). 10.1016/j.cell.2020.09.057

15. Khaliq, Z. M., Gouwens, N. W. & Raman, I. M. The contribution of resurgent sodium current to high-frequency firing in Purkinje neurons: an experimental and modeling study. J. Neurosci. 23, 4899–4912 (2003). 10.1523/JNEUROSCI.23-12-04899.2003

16. Sturm, P., Wimmers, S., Schwarz, J. R. & Bauer, C. K. Extracellular potassium effects are conserved within the rat erg K⁺ channel family. J. Physiol. 564, 329–345 (2005).

17. Strauss, K. A. et al. A population-based study of KCNH7 p.Arg394His and bipolar spectrum disorder. Hum. Mol. Genet. 23, 6395–6406 (2014).

18. Watkins, C. J. C. H. & Dayan, P. Q-learning. Mach. Learn. 8, 279–292 (1992). doi:10.1007/BF00992698.

19. Diehl, P. U. & Cook, M. Unsupervised learning of digit recognition using spike-timing-dependent plasticity. Front. Comput. Neurosci. 9, 99 (2015). doi:10.3389/fncom.2015.00099.

20. Hinton, G. E. Training products of experts by minimizing contrastive divergence. Neural Comput. 14, 1771–1800 (2002). doi:10.1162/089976602760128018.

21. Tieleman, T. Training restricted Boltzmann machines using approximations to the likelihood gradient. In Proceedings of the 25th International Conference on Machine Learning, 1064–1071 (ACM, 2008). doi:10.1145/1390156.1390290.

22. Scala, F. et al. Phenotypic variation of transcriptomic cell types in mouse motor cortex. Nature 598, 144–150 (2021). 10.1038/s41586-020-2907-3

23. Mickelsen, L. E. et al. Cellular taxonomy and spatial organization of the murine ventral posterior hypothalamus. eLife 9, e58901 (2020). 10.7554/eLife.58901

24. Codazzi, F. et al. Synergistic control of protein kinase Cgamma activity by ionotropic and metabotropic glutamate receptor inputs in hippocampal neurons. J Neurosci 26, 3404–3411 (2006). 10.1523/jneurosci.0478-06.2006

25. Waltz, J. A. & Gold, J. M. Probabilistic reversal learning impairments in schizophrenia: further evidence of orbitofrontal dysfunction. Schizophr Res 93, 296–303 (2007). 10.1016/j.schres.2007.03.010

26. Griffin, J. D. et al. Distinct alterations in probabilistic reversal learning across at-risk mental state, first episode psychosis and persistent schizophrenia. Sci Rep 14, 17614 (2024). 10.1038/s41598-024-68004-7

27. Gastambide, F. et al. Selective remediation of reversal learning deficits in the neurodevelopmental MAM model of schizophrenia by a novel mGlu5 positive allosteric modulator. Neuropsychopharmacology 37, 1057–1066 (2012). 10.1038/npp.2011.298

28. Parmentier-Batteur, S. et al. Mechanism based neurotoxicity of mGlu5 positive allosteric modulators — development challenges for a promising novel antipsychotic target. Neuropharmacology 82, 161–173 (2014). 10.1016/j.neuropharm.2012.12.003

29. Sturm, S. et al. Results and evaluation of a first-in-human study of RG7342, an mGlu5 positive allosteric modulator, utilizing Bayesian adaptive methods. Br J Clin Pharmacol 84, 445–455 (2018). 10.1111/bcp.13466

30. Rook, J. M. et al. Biased mGlu5-Positive Allosteric Modulators Provide In Vivo Efficacy without Potentiating mGlu5 Modulation of NMDAR Currents. Neuron 86, 1029–1040 (2015). 10.1016/j.neuron.2015.03.063

31. LeCun, Y., Bottou, L., Bengio, Y. & Haffner, P. Gradient-based learning applied to document recognition. Proc. IEEE 86, 2278–2324 (1998). doi:10.1109/5.726791.

32. Lein, E. S. et al. Genome-wide atlas of gene expression in the adult mouse brain. Nature 445, 168–176 (2007). 10.1038/nature05453

33. Allen Institute for Brain Science. Allen Mouse Brain Atlas [dataset]. https://mouse.brain-map.org/

34. Brandt, M. D. et al. Transient calretinin expression defines early postmitotic step of neuronal differentiation in adult hippocampal neurogenesis of mice. Mol. Cell. Neurosci. 24, 603–613 (2003).

35. Kobayashi, K., et al. Reversal of hippocampal neuronal maturation by serotonergic antidepressants. Proc. Natl Acad. Sci. USA 107, 8434–8439 (2010).

36. Lee, B. R. et al. Scaled, high fidelity electrophysiological, morphological, and transcriptomic cell characterization. eLife 10, e65482 (2021).

37. Martini, L. et al. Neuronal Spike Shapes (NSS): A straightforward approach to investigate heterogeneity in neuronal excitability states. Comput. Biol. Med. 168, 107783 (2024). 10.1016/j.compbiomed.2023.107783

38. McDougal, R. A. et al. Twenty years of ModelDB and beyond: building essential modeling tools for the future of neuroscience. J. Comput. Neurosci. 42, 1–10 (2017). 10.1007/s10827-016-0623-7

39. Raman, I. M. & Bean, B. P. Inactivation and recovery of sodium currents in cerebellar Purkinje neurons: evidence for two mechanisms. Biophys. J. 80, 729–737 (2001). 10.1016/S0006-3495(01)76052-3

40. Stimberg, M., Brette, R. & Goodman, D. F. M. Brian 2, an intuitive and efficient neural simulator. eLife 8, e47314 (2019). doi:10.7554/eLife.47314.

41. Alevi, D., Stimberg, M., Sprekeler, H., Obermayer, K. & Augustin, M. Brian2CUDA: Flexible and efficient simulation of spiking neural network models on GPUs. Front. Neuroinform. 16, 883700 (2022). doi:10.3389/fninf.2022.883700.

42. Pfister, J.-P. & Gerstner, W. Triplets of spikes in a model of spike timing-dependent plasticity. J. Neurosci. 26, 9673–9682 (2006). doi:10.1523/JNEUROSCI.1425-06.2006.

43. Hines, M. L. & Carnevale, N. T. The NEURON simulation environment. Neural Comput. 9, 1179–1209 (1997). 10.1162/neco.1997.9.6.1179

44. Mackey, M. C. & Glass, L. Oscillation and chaos in physiological control systems. Science 197, 287–289 (1977). 10.1126/science.267326

45. Dambre, J., Verstraeten, D., Schrauwen, B. & Massar, S. Information processing capacity of dynamical systems. Sci. Rep. 2, 514 (2012). 10.1038/srep00514

46. Hounkpe, B. W., Chenou, F., de Lima, F. & De Paula, E. V. HRT Atlas v1.0 database: redefining human and mouse housekeeping genes and candidate reference transcripts by mining massive RNA-seq datasets. Nucleic Acids Res. 49, D947–D955 (2021). 10.1093/nar/gkaa609

47. Schreiber, T. & Schmitz, A. Improved surrogate data for nonlinearity tests. Phys. Rev. Lett. 77, 635–638 (1996). 10.1103/PhysRevLett.77.635

