## Supplementary Fig for "Maturation-associated ERG3 and timed regulation of circuit stability"

**Supplementary Fig. S3-1**

### Passive-state measurements in the ERG3 Purkinje-cell model

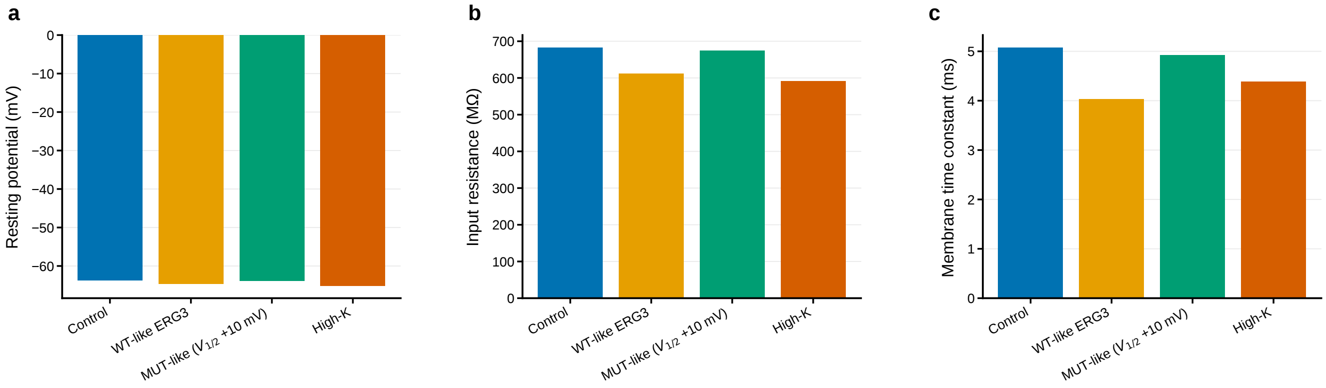


a, Resting membrane potential estimated from the final 50 ms before the selected passive current step. b, Input resistance estimated from the slope of steady-state voltage deflection against injected current across non-spiking hyperpolarizing-step responses. c, Membrane time constant estimated from a log-linear fit of the normalized passive voltage response. Conditions were Control, WT-like ERG3, MUT-like (V1/2 +10 mV) and High-K. Measurements were obtained from noise-free simulations; the step-selection and fitting procedures are described in Methods.

**Supplementary Fig. S3-2 · a**

### Count- and timing-based response contrasts across input offsets and decision windows

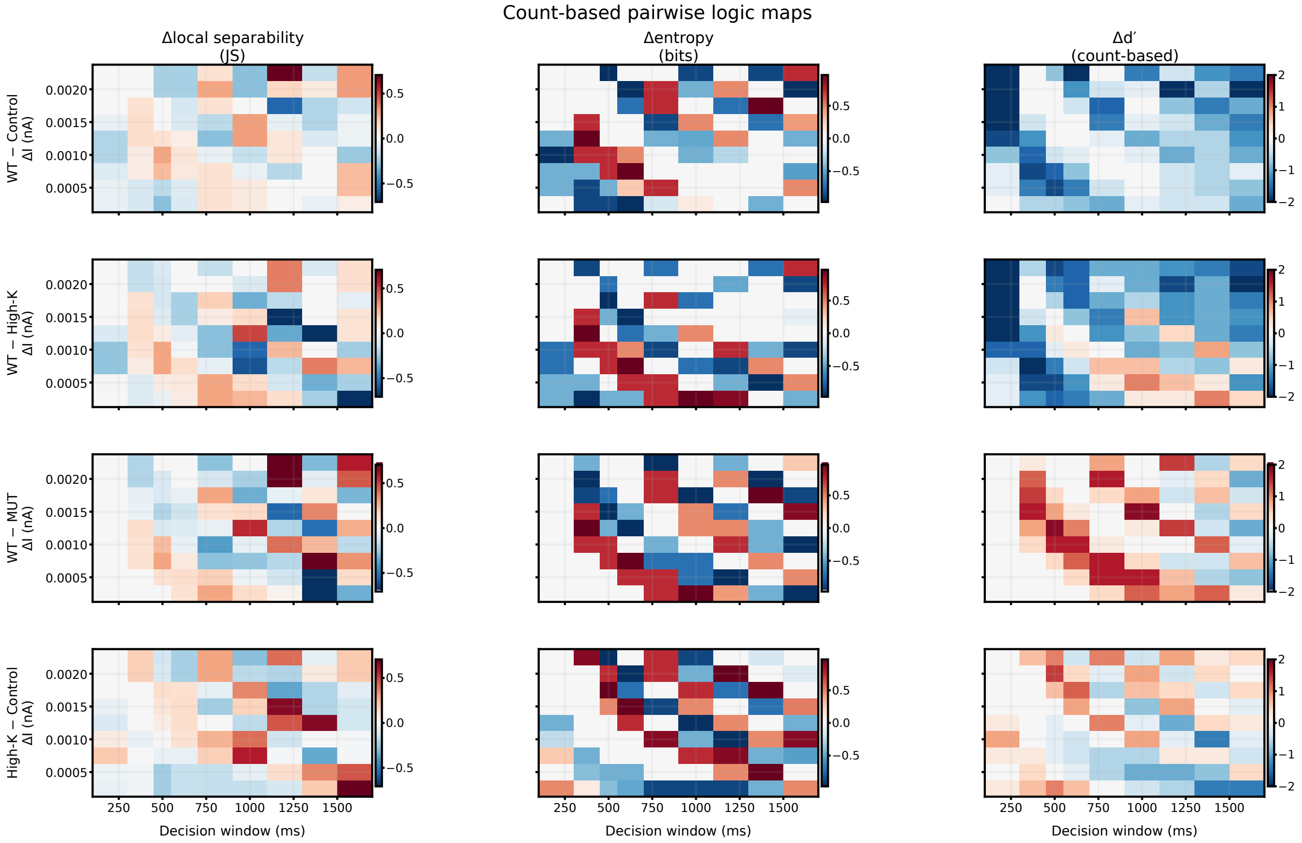


a, Count-based pairwise maps around condition-specific operating currents referenced to an 80-Hz firing-rate target. Columns show differences in local separability, response entropy and count-based d′. Rows show WT-like ERG3 minus Control, WT-like ERG3 minus High-K, WT-like ERG3 minus MUT-like (V1/2 +10 mV), and High-K minus Control. Local separability was based on Jensen–Shannon divergence between count distributions at adjacent current offsets; entropy was calculated within each shifted-input distribution; d′ compared shifted and baseline inputs.

**Supplementary Fig. S3-2 · b**

### Count- and timing-based response contrasts across input offsets and decision windows

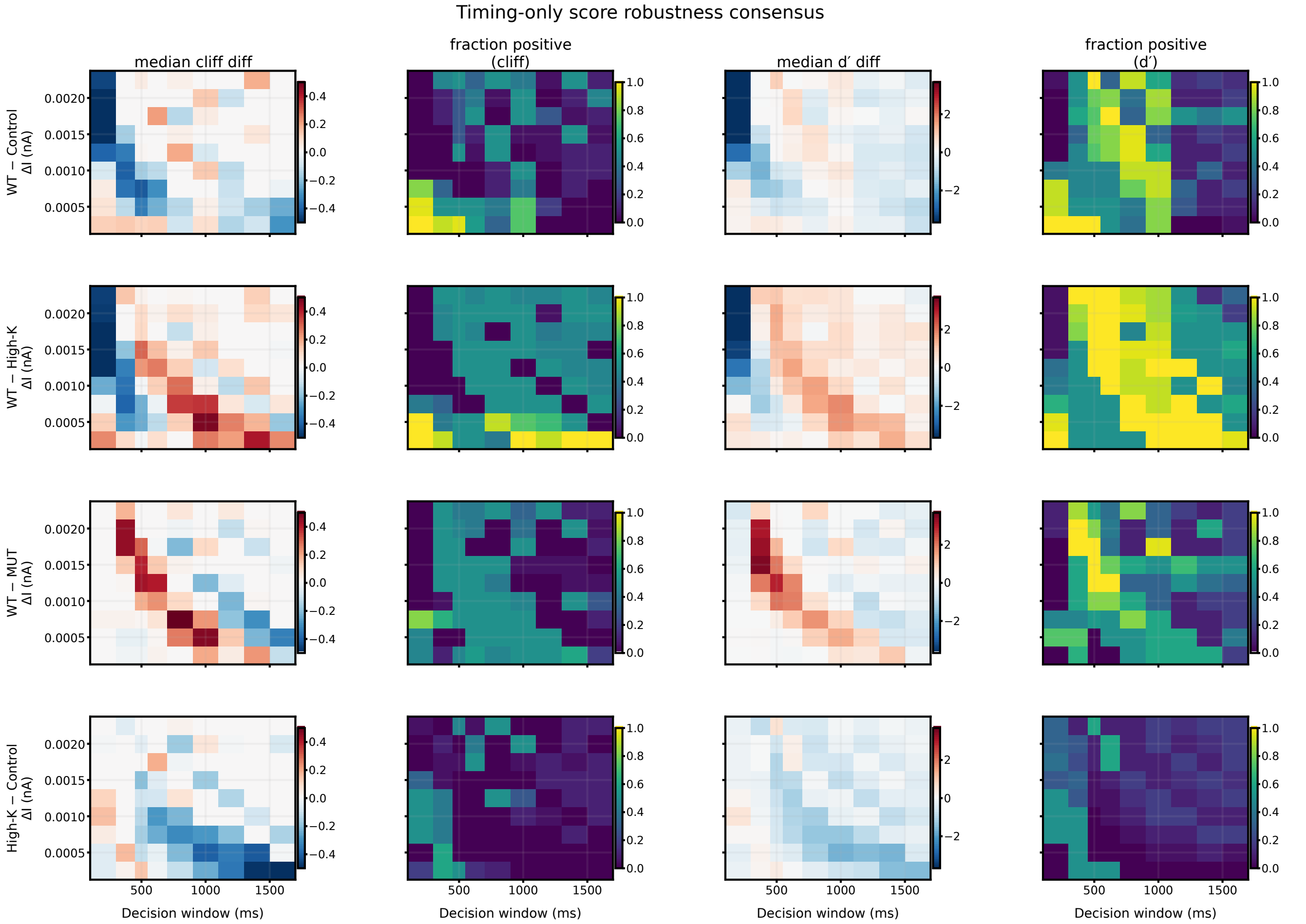


b, Timing-score consensus for the same contrasts. Maps show median signed differences and fractions of positive differences for Cliff’s delta and timing-based d′ across 22 timing-score configurations, excluding the count-only control. Configurations included vector-strength, Rayleigh-related, linear ISI and log-ISI scores, with and without count residualization. Summaries used finite estimates only; the number of contributing configurations is provided in Source Data.

Supplementary Fig. S5-1 · a

### Held-out forecast trajectories

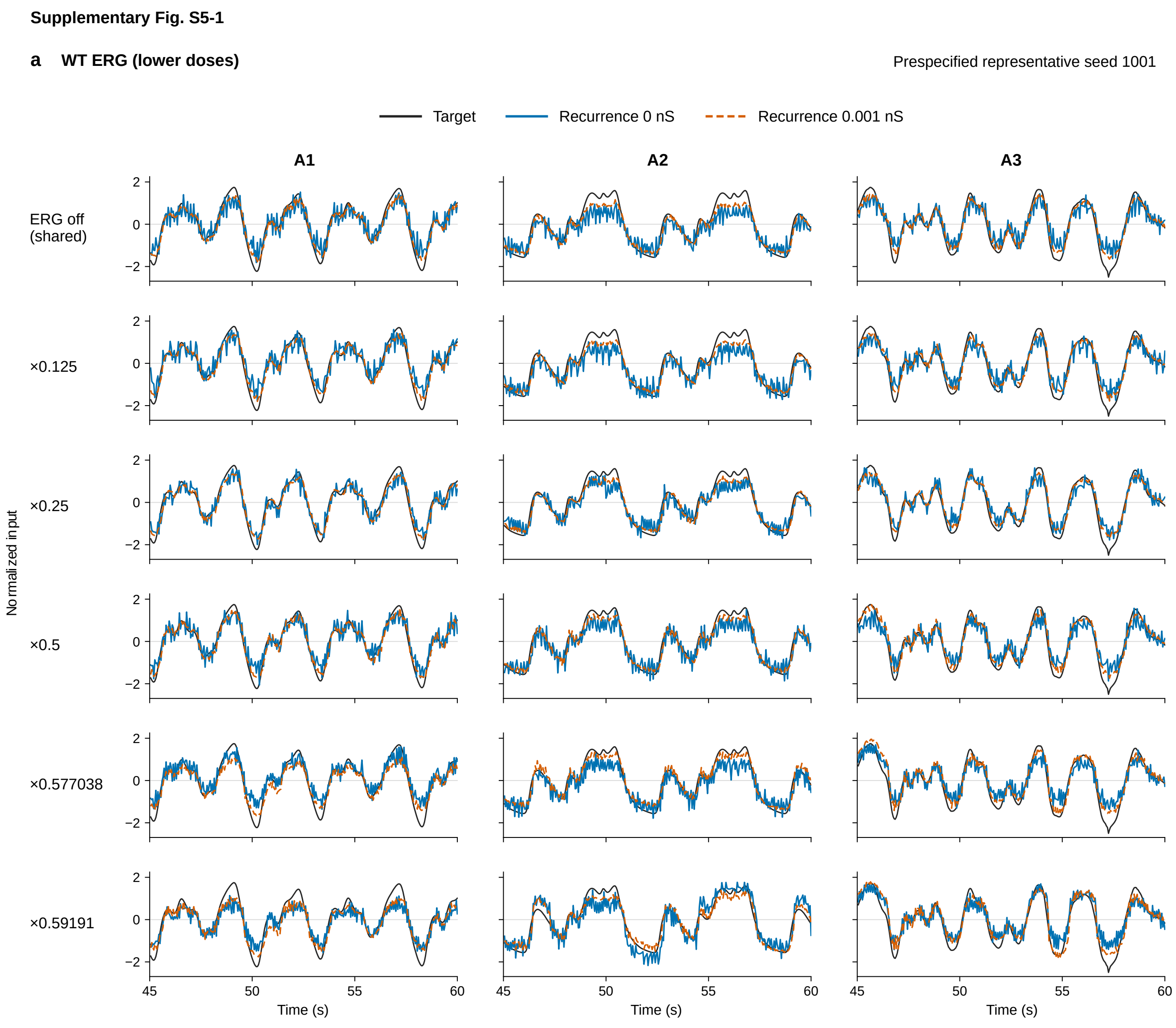


Original held-out samples at 50-ms intervals. Target and prediction are in normalized input units.
Identical or near-identical traces may overlap. The shared ERG-off / native Fast K control is in a.

Native WT ERG, lower conductances. Columns show A1–A3 and rows show conductance factors. All plots show the 300 original held-out samples from representative network seed 1001, including the target and the predictions at recurrence caps of 0 and 0.001 nS. The target is the next 50-ms input sample. The shared ERG-off condition appears in a. Overlapping traces can obscure a line. All panels share the same ordinate.

Supplementary Fig. S5-1 · b

### Held-out forecast trajectories

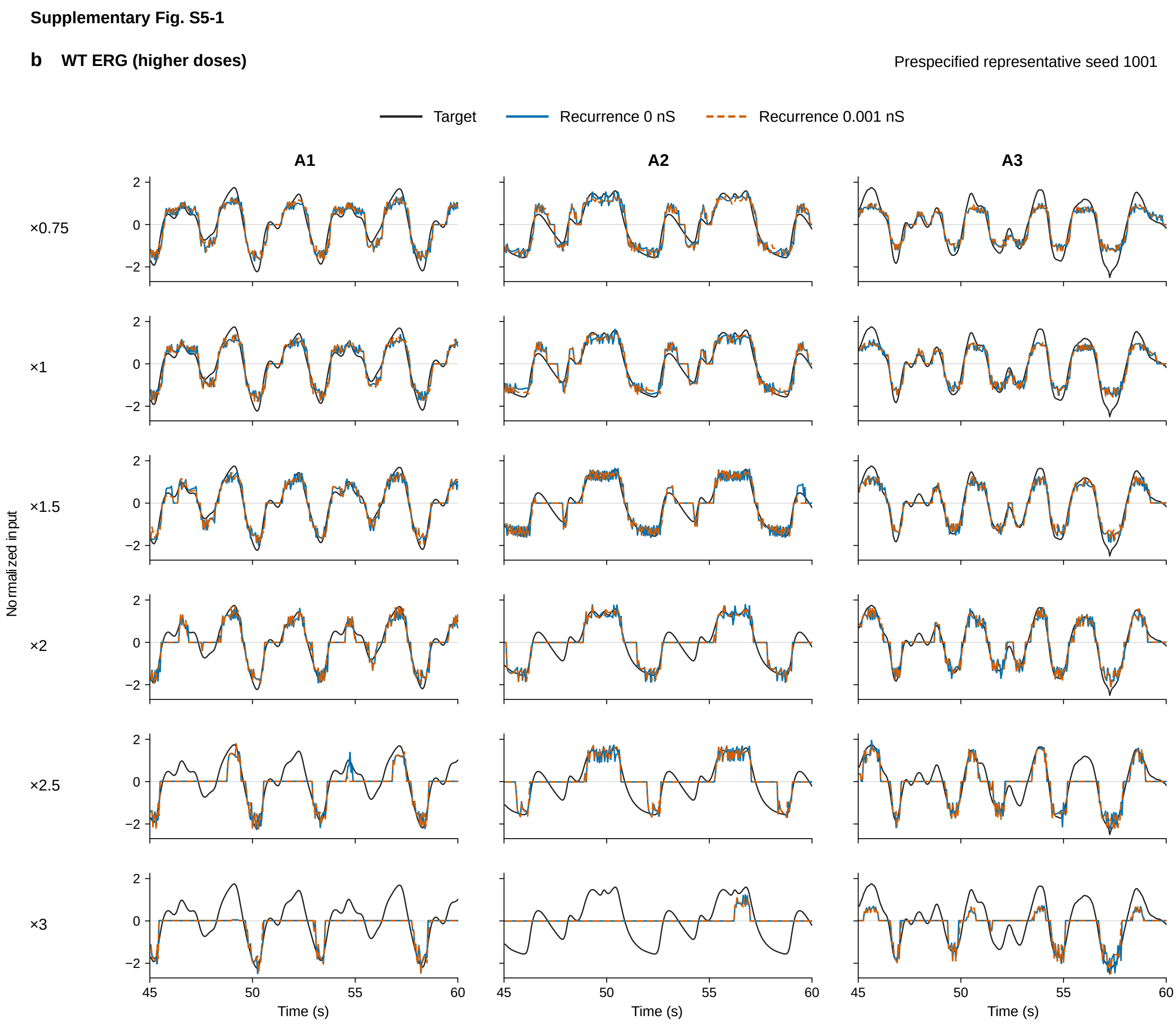


Original held-out samples at 50-ms intervals. Target and prediction are in normalized input units.
Identical or near-identical traces may overlap. The shared ERG-off / native Fast K control is in a.

Native WT ERG, higher conductances. Columns show A1–A3 and rows show conductance factors. All plots show the 300 original held-out samples from representative network seed 1001, including the target and the predictions at recurrence caps of 0 and 0.001 nS. The target is the next 50-ms input sample. The shared ERG-off condition appears in a. Overlapping traces can obscure a line. All panels share the same ordinate.

#### Supplementary Fig. S5-1 · c

### Held-out forecast trajectories

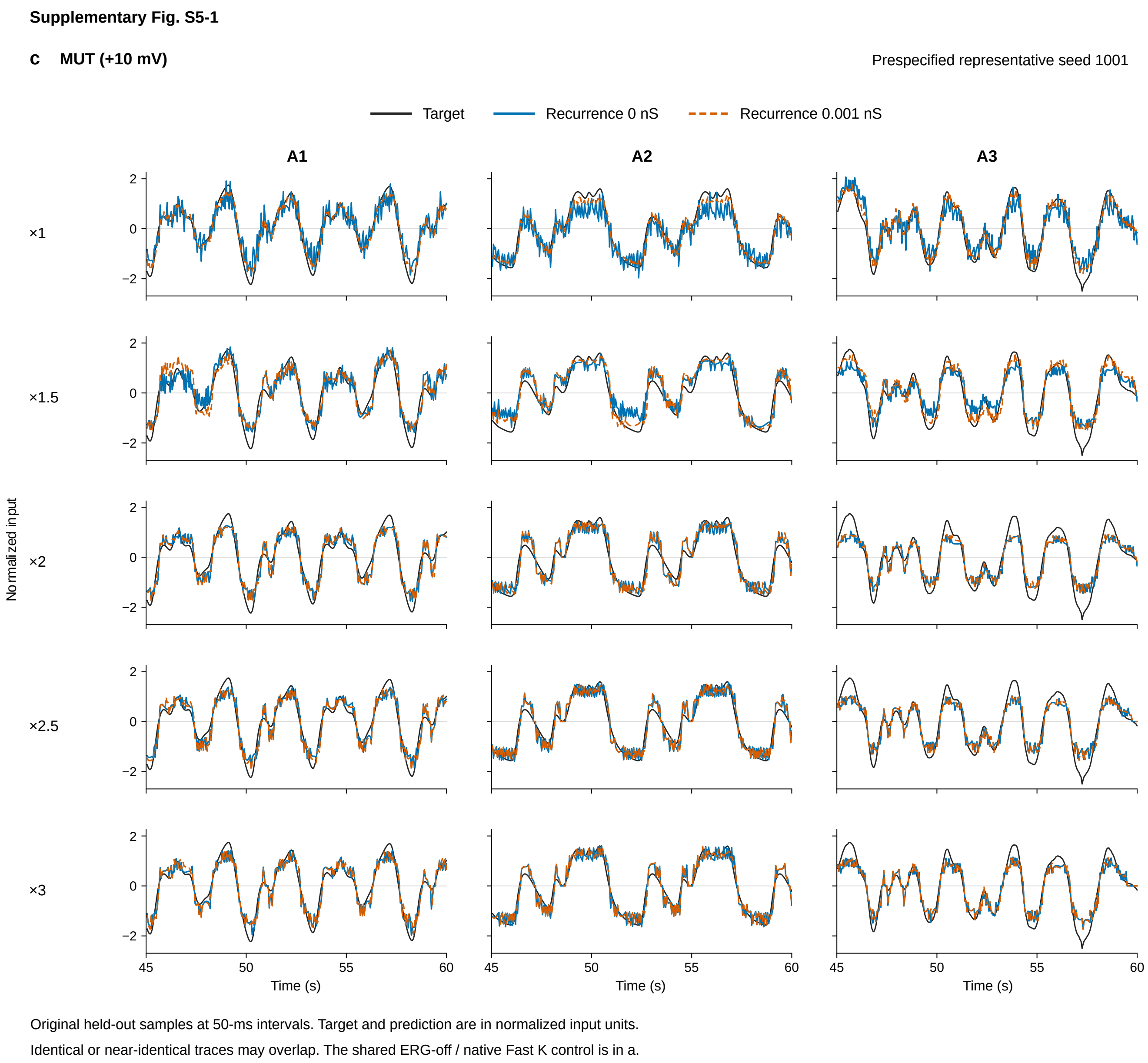


Original held-out samples at 50-ms intervals. Target and prediction are in normalized input units.
Identical or near-identical traces may overlap. The shared ERG-off / native Fast K control is in a.

MUT. Columns show A1–A3 and rows show conductance factors. All plots show the 300 original held-out samples from representative network seed 1001, including the target and the predictions at recurrence caps of 0 and 0.001 nS. The target is the next 50-ms input sample. The shared ERG-off condition appears in a. Overlapping traces can obscure a line. All panels share the same ordinate.

Supplementary Fig. S5-1 · d

### Held-out forecast trajectories

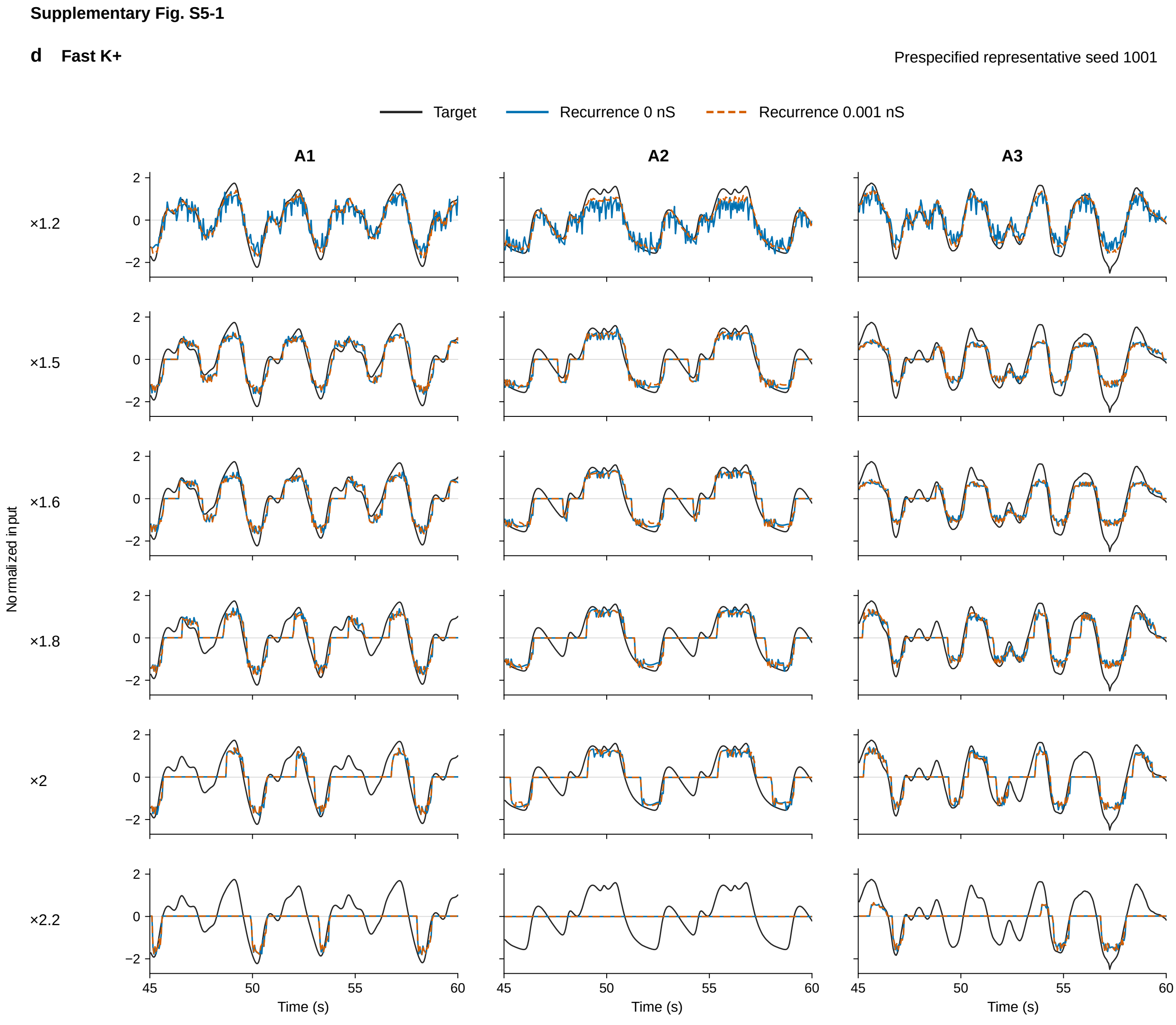


Original held-out samples at 50-ms intervals. Target and prediction are in normalized input units.
Identical or near-identical traces may overlap. The shared ERG-off / native Fast K control is in a.

Fast K+ through ×2.2. Columns show A1–A3 and rows show conductance factors. All plots show the 300 original held-out samples from representative network seed 1001, including the target and the predictions at recurrence caps of 0 and 0.001 nS. The target is the next 50-ms input sample. The shared ERG-off condition appears in a. Overlapping traces can obscure a line. All panels share the same ordinate.

##### Supplementary Fig. S5-1 · e

### Held-out forecast trajectories

**Supplementary Fig. S5-1**

**e WT ERG, activation 10×**

Prespecified representative seed 1001

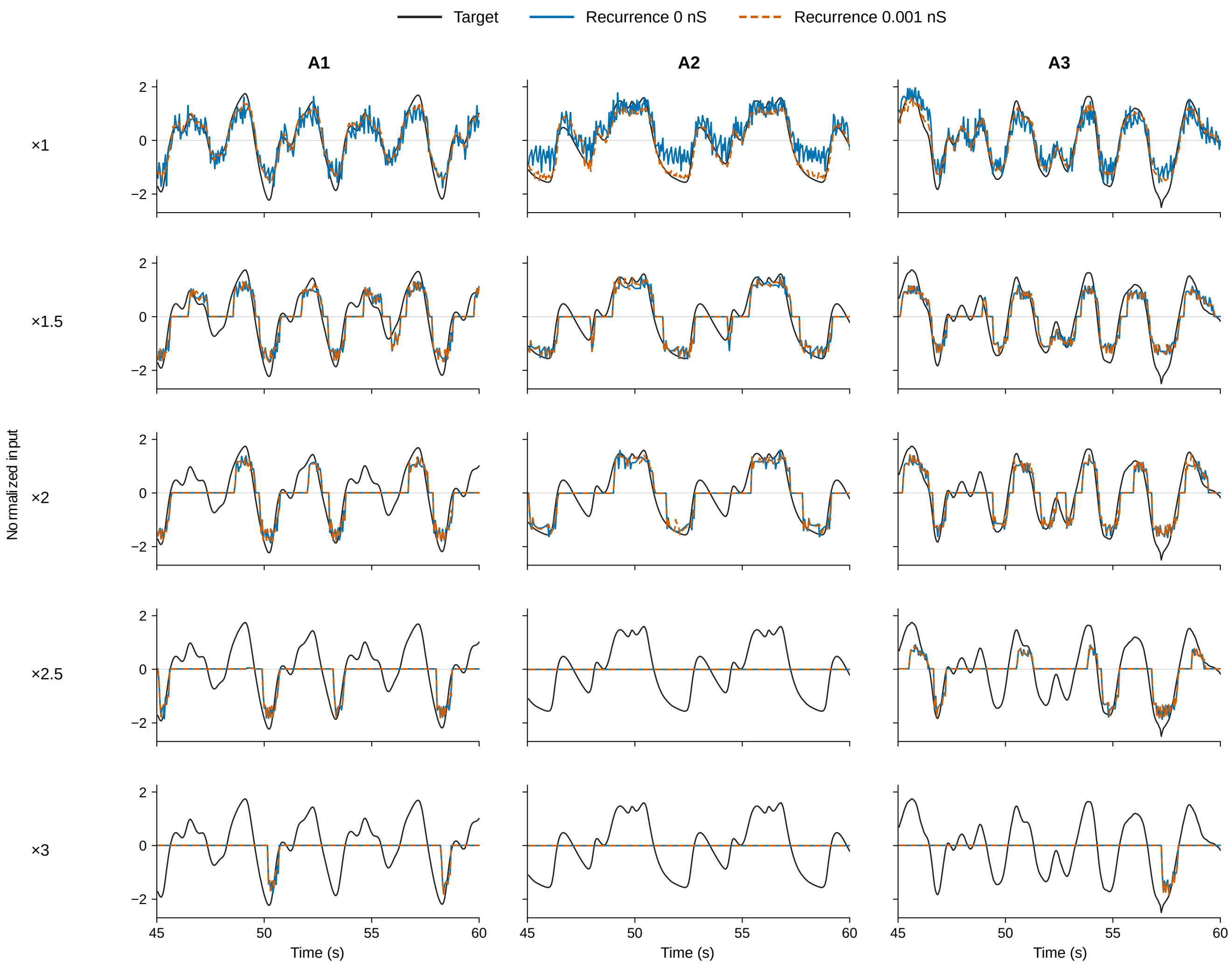


Original held-out samples at 50-ms intervals. Target and prediction are in normalized input units.
Identical or near-identical traces may overlap. The shared ERG-off / native Fast K control is in a.

WT ERG with tenfold accelerated activation. Columns show A1–A3 and rows show conductance factors. All plots show the 300 original held-out samples from representative network seed 1001, including the target and the predictions at recurrence caps of 0 and 0.001 nS. The target is the next 50-ms input sample. The shared ERG-off condition appears in a. Overlapping traces can obscure a line. All panels share the same ordinate.

**Supplementary Fig. S5-1 · f**

### Held-out forecast trajectories

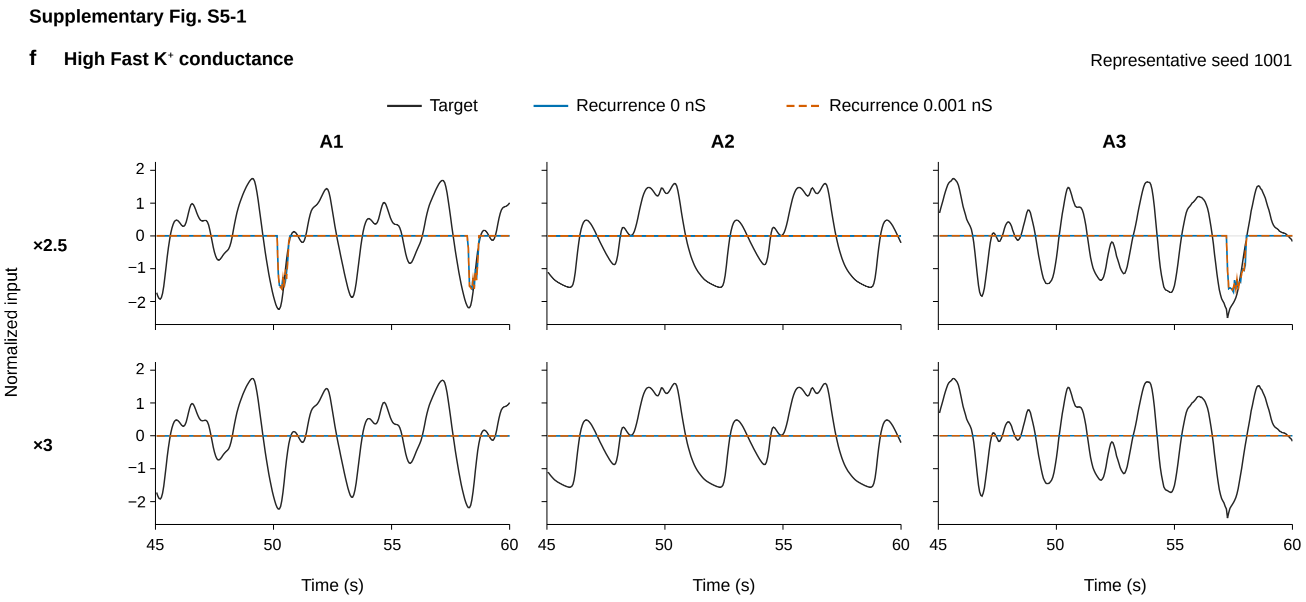


Fast K+ ×2.5 and ×3. Columns show A1–A3 and rows show conductance factors. All plots show the 300 original held-out samples from representative network seed 1001, including the target and the predictions at recurrence caps of 0 and 0.001 nS. The target is the next 50-ms input sample. The shared ERG-off condition appears in a. Overlapping traces can obscure a line. All panels share the same ordinate.

##### Supplementary Fig. S5-2

### Forecast error relative to persistence

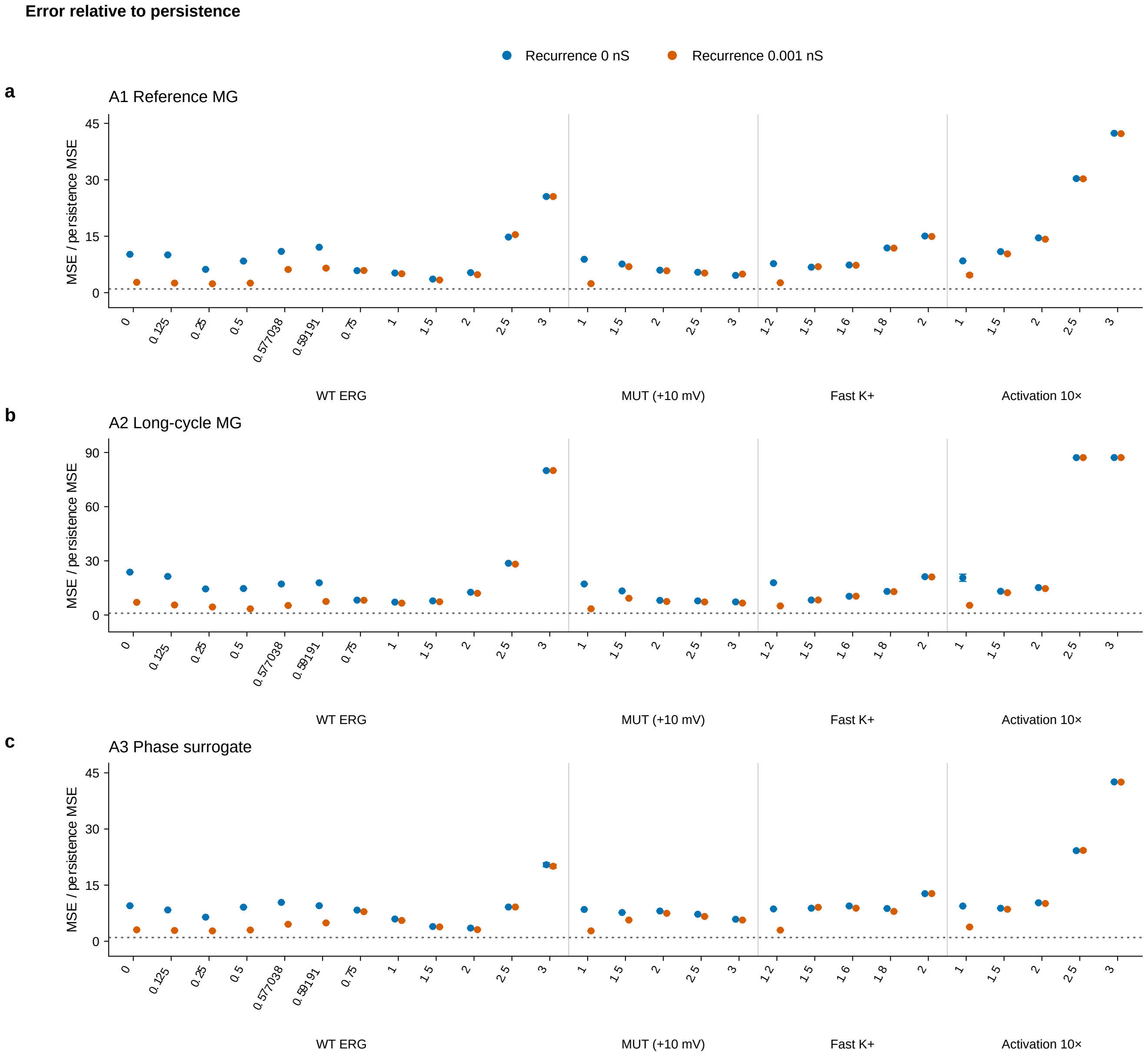


Test MSE is divided by that of predicting the next sample from the current sample using the same 300 targets. Panels a–c show A1–A3. Values above one indicate larger errors than persistence. Points and intervals summarize 30 seeds per condition in the original dose grid.

Supplementary Fig. S5-3 · a

### Forecast errors and spiking at high Fast K+ conductance

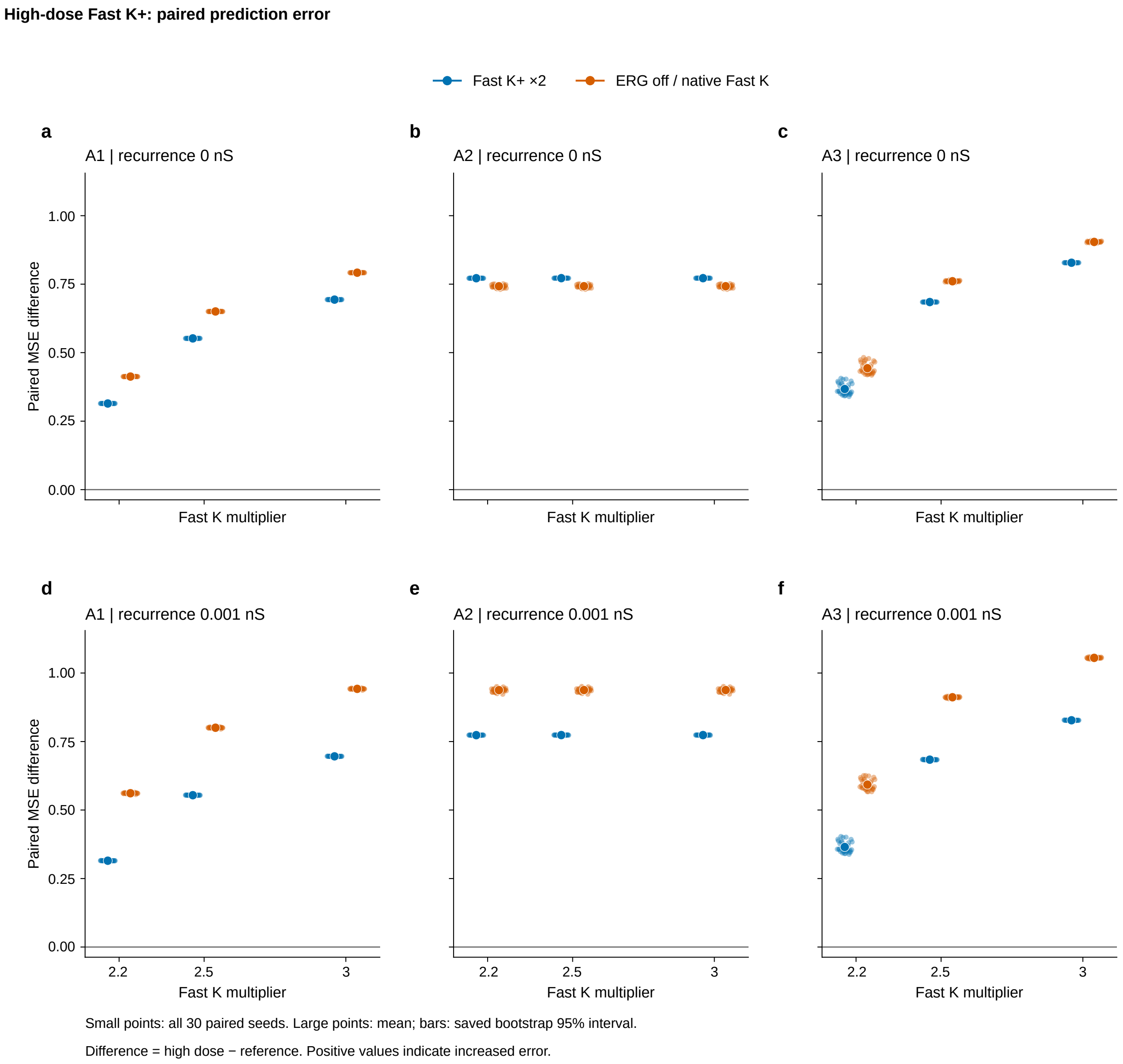


a, Paired test-MSE changes at Fast K+ ×2.2, ×2.5 and ×3 relative to Fast K+ ×2 and ERG off. Positive differences indicate larger error at the higher conductance. Points show 30 matched seed differences and summaries show means with pointwise 95% seed-bootstrap intervals.

Supplementary Fig. S5-3 · b

### Forecast errors and spiking at high Fast K+ conductance

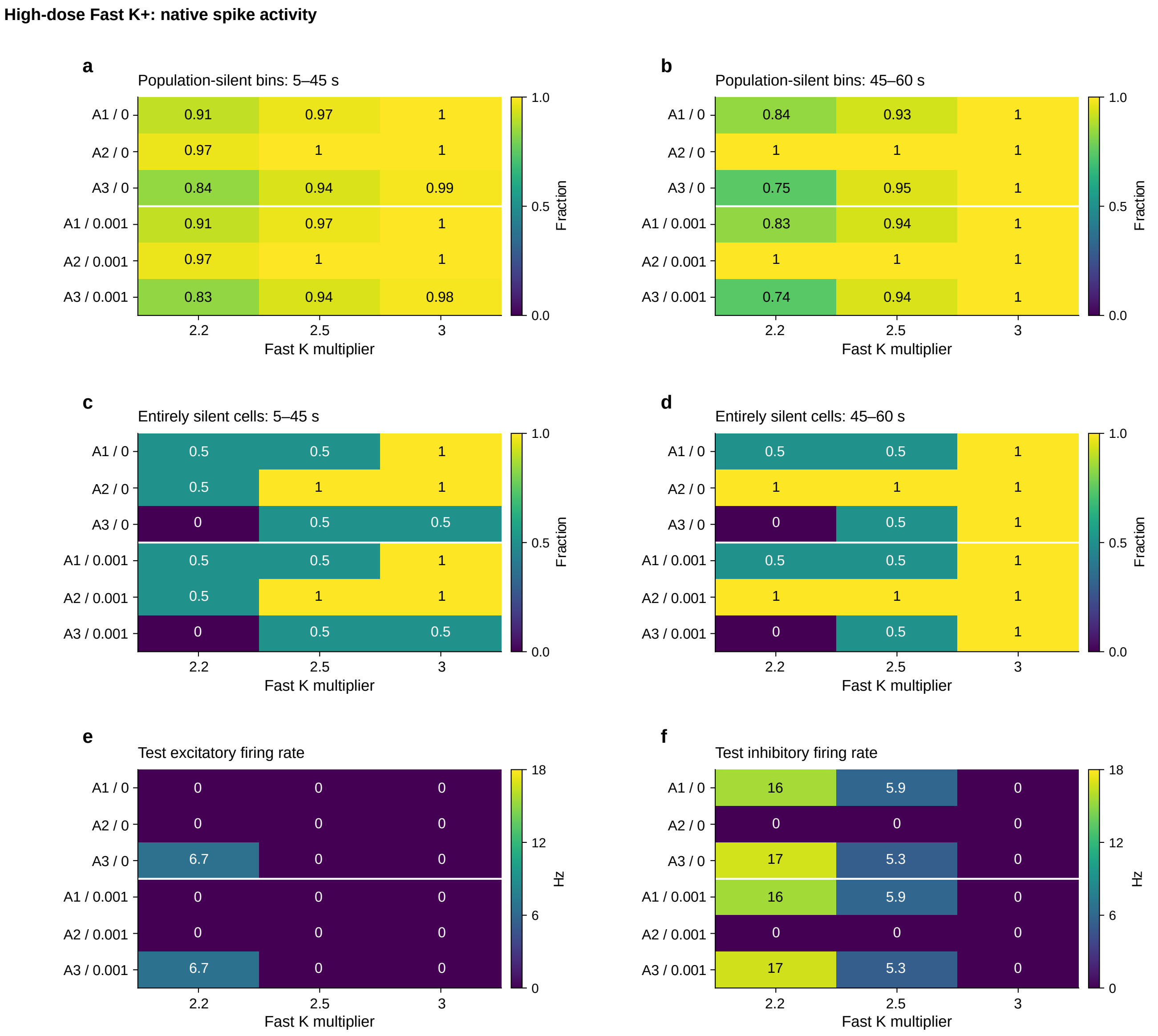


Rows: input / recurrence cap (nS). Cell values: means across 30 network seeds.

Fraction scales: 0–1. E and I firing-rate scales: 0–18 Hz. NA denotes unavailable data.

b, Native excitatory and inhibitory activity, zero-spike bins and silent cells in the diagnostic (5–45 s) and test (45–60 s) windows. Displayed rates refer to the test window. Heatmap cells are means over 30 seeds. NA denotes undefined or unavailable measurements.

##### Supplementary Fig. S5-4

### Input structure and forecast contrasts

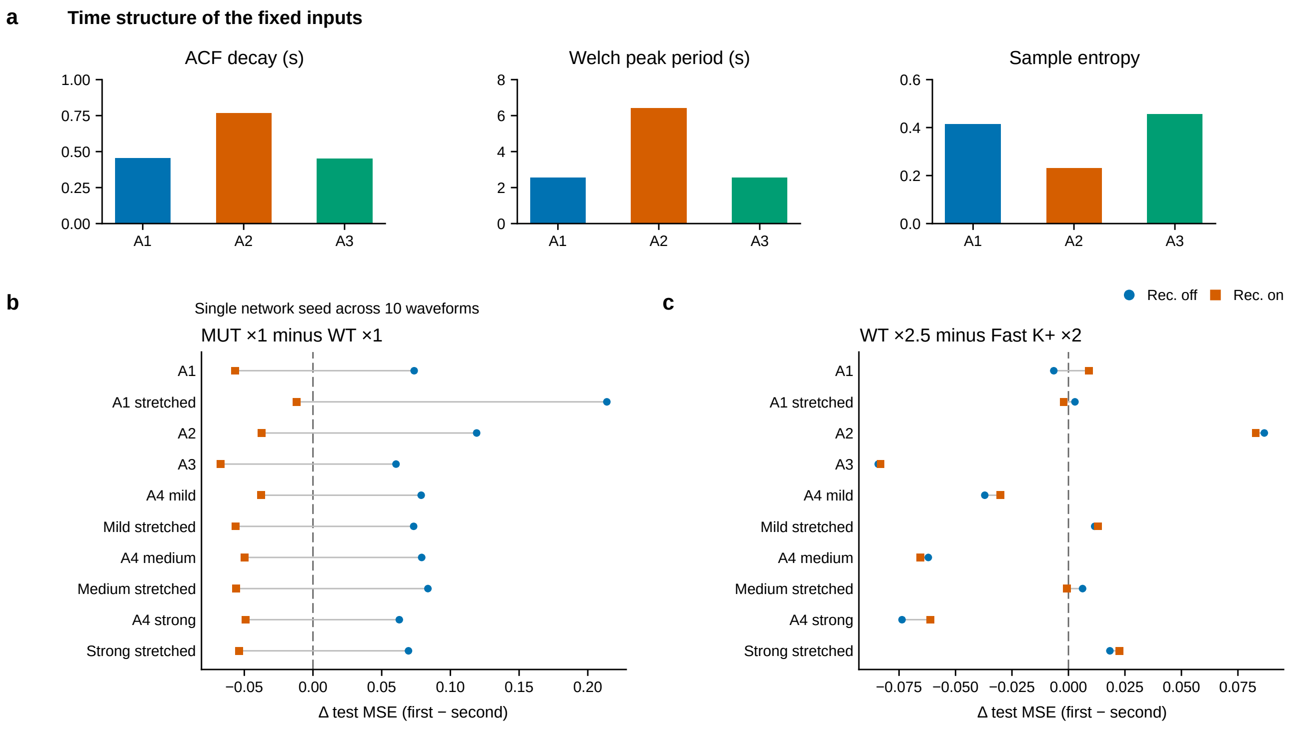


a, Autocorrelation decay time (first interpolated 1/e crossing), dominant Welch spectral period and sample entropy (embedding dimension 2 and tolerance 0.2 standard deviations) for A1–A3. b,c, Forecast MSE contrasts for ten fixed inputs in one exploratory network seed. Positive values indicate higher MSE for the first named channel condition. Connected symbols compare recurrence caps. A4 is a regularized surrogate and temporal stretching used interpolation.

##### Supplementary Fig. S6-1 · a

### Schedule effects under independent input realizations

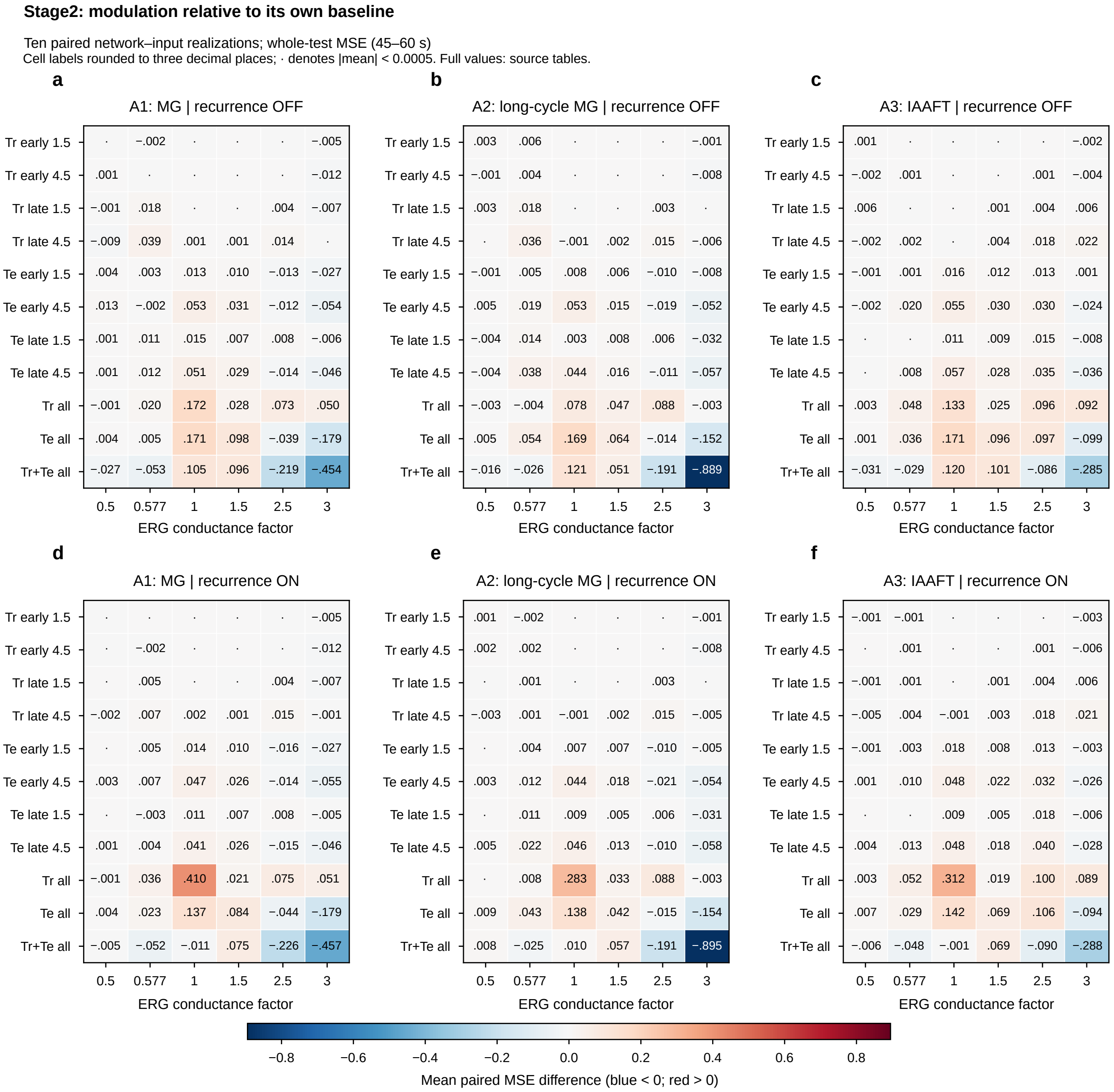


The first sheet shows Stage 2 means across the full schedule and conductance grid for each input and recurrence cap. Each intervention effect subtracts the matched unshifted baseline MSE. Stage 1 uses a fixed waveform per input family. Stage 2 pairs each network seed with one independently constructed waveform per family and reuses that waveform across conditions. Each cell summarizes ten pairs. The test window is 45–60 s.

#### Supplementary Fig. S6-1 · b

### Schedule effects under independent input realizations

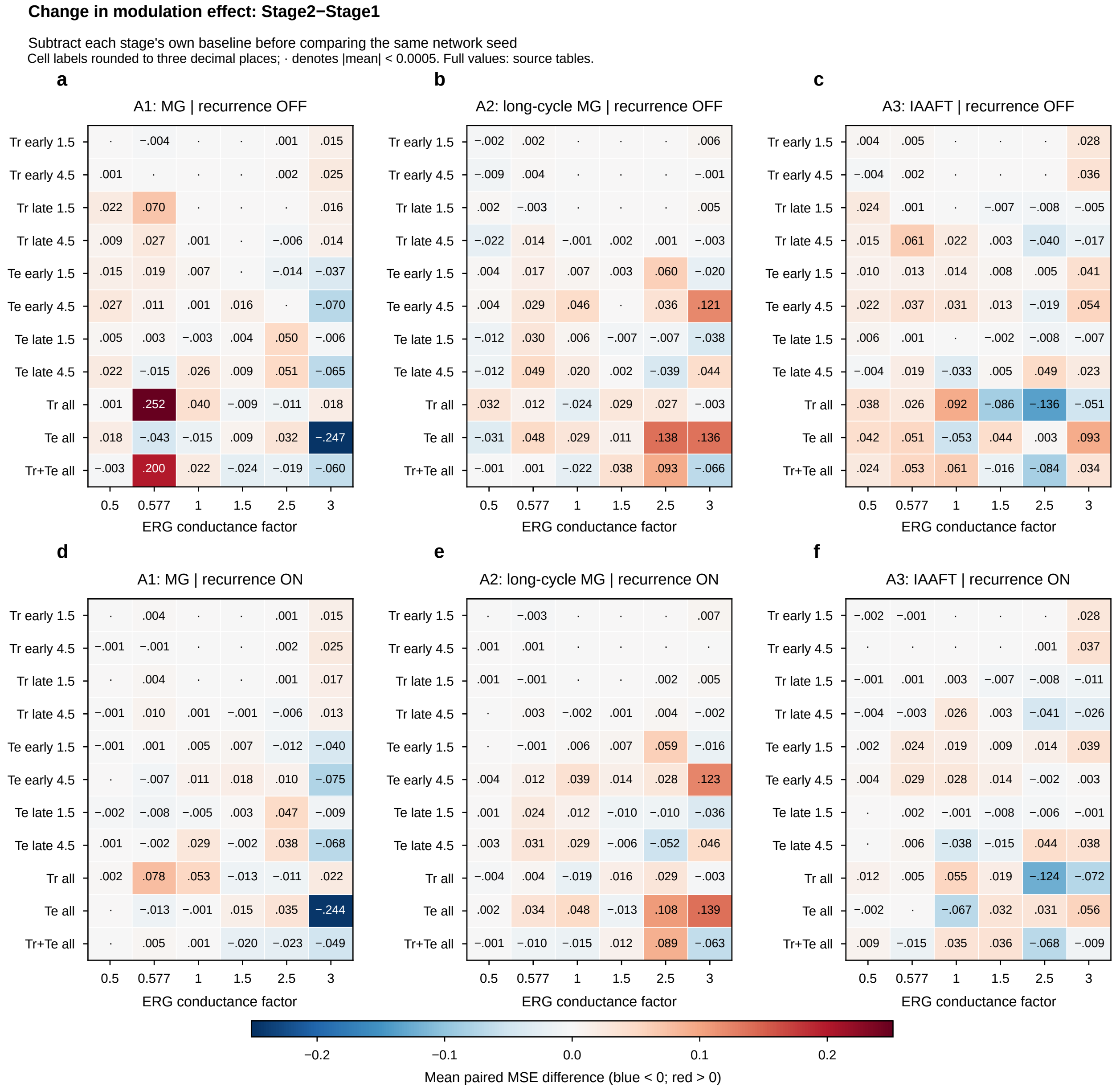


Stage 2 minus Stage 1 differences in matched intervention effects. Stage 1 uses a fixed waveform per input family. Stage 2 pairs each network seed with one independently constructed waveform per family and reuses that waveform across conditions. Each cell summarizes ten pairs. The test window is 45–60 s.

##### Supplementary Fig. S6-2

### Associations for all 18 Gq-linked receptors

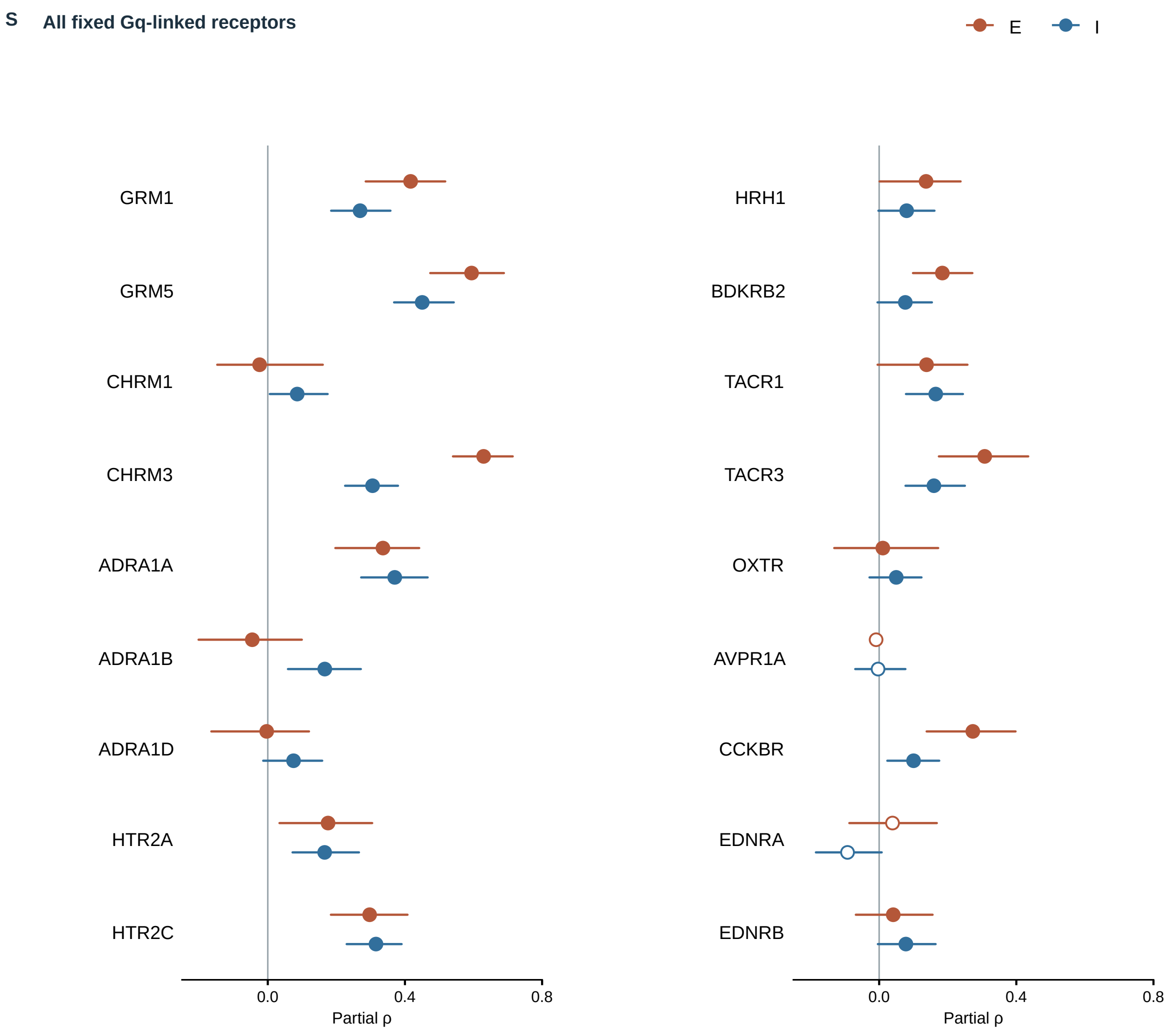


Adjusted rank correlations with *Kcnh7* are shown for the full predefined receptor set, grouped by ligand family. Estimates are shown separately for excitatory and inhibitory cells of mouse M1. Bars are saved pointwise 95% mouse-bootstrap intervals. Hollow points indicate fewer than ten expression-positive cells. The excitatory AVPR1A interval is unavailable under the bootstrap validity criterion.

##### Supplementary Fig. S8-1

### Timing sensitivity of a matched shallow deficit

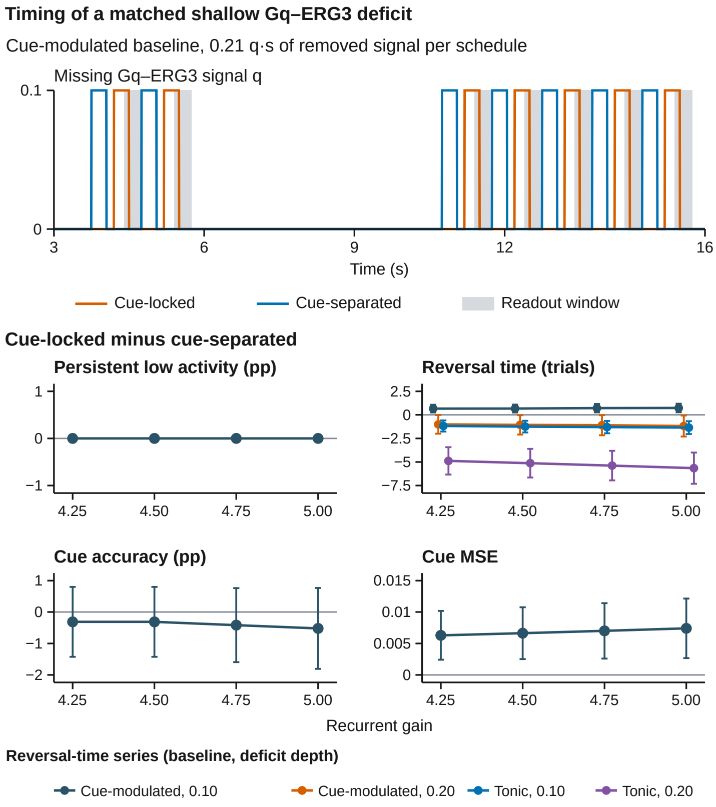


The waveform panel shows the cue-locked and cue-separated 300-ms deficits over the task, with the scoring intervals shaded. Lower panels show paired differences (cue-locked minus cue-separated) in persistent low activity, value-reversal time, cue-sign accuracy and cue MSE at each recurrent gain. The low-depth condition removes 0.10 q at each of seven cues (0.21 q·s total). Additional series in the latency panel compare peak deficits of 0.10 q and 0.20 q on cue-modulated and tonic baselines of equal mean q. Each condition uses 24 matched cue/noise trajectories per circuit, and the value analysis uses 512 matched sessions per circuit. Error bars, pointwise 95% paired t intervals across eight circuits.

**Supplementary Table 1**

### ERG3 gating measurements and conductance-model calibration

| Parameter | Niculescu et al. (2013) | Sturm et al. (2005) | Simulation | Modified | KCNH7 MUT |
|---|---|---|---|---|---|
| V½ (mV) | −44.2 | −45.1 | −45.9 | −46.19 | −36.74 |
| Slope, k (mV) | 4.2 | 8.5 / 6.2 | 13.7 | 3.76 | 3.56 |
| Activation (ms) | — | 50.1 | 19.5 | 46.99 | 47 |
| Inactivation (ms) | — | Fig. 4Bc | 17.2 | 22.82 | 22.81 |
| Recovery (ms) | 2.1 | Fig. 6Ac | 1.1 | 1.29 | 1.29 |
| Deactivation (ms) | 16.9 | Fig. 6Bc, Cc | 22 | 14.99 | 14.99 |

**Niculescu et al.** V½ denotes the activation midpoint. Table 3 reports recombinant rat ERG3 expressed in HEK293 cells with 40 mM extracellular $K^+$. Recovery from inactivation and deactivation were measured at −100 mV. Dashes indicate parameters omitted from this calibration column.

**Sturm et al.** In CHO cells expressing rat ERG3, V½ was fitted to the mean activation curve at 10 mM extracellular $K^+$ (Fig. 2Bc). The two slope factors are means of individual fits at 2 and 10 mM $K^+$, respectively. The activation time constant was measured at +20 mV in 2 mM $K^+$ (Fig. 3Bc). The figure references identify voltage-dependent time courses, including fast and slow deactivation components.

**Model values.** The three right-hand columns give the historical calibration values. The Fig. 3 simulations used steady-state activation midpoints of −45.1 mV (WT) and −35.1 mV (MUT), both with a slope factor of 4.2 mV.

Niculescu D, Hirdes W, Hornig S, Pongs O, Schwarz JR. Erg potassium currents of neonatal mouse Purkinje cells exhibit fast gating kinetics and are inhibited by mGluR1 activation. J Neurosci. 33, 16729–16740 (2013). doi:10.1523/JNEUROSCI.5523-12.2013.

Sturm P, Wimmers S, Schwarz JR, Bauer CK. Extracellular potassium effects are conserved within the rat erg K+ channel family. J Physiol. 564, 329–345 (2005). doi:10.1113/jphysiol.2004.078840.

**Supplementary Table 2**

### State retention after sustained signal loss

At background drive −0.65, the same 12 circuits and four noise realisations per circuit were followed under intact modulation, sustained loss, and sustained loss with compensatory drive (Fig. 8a). Counts refer to the 48 planned trials in each condition.

| Condition | Planned | Established | Failed to establish | First exit | Retained at 120 s |
|---|---|---|---|---|---|
| Intact modulation | 48 | 48 | 0 | 0 | 48 |
| Sustained signal loss | 48 | 44 | 4 | 14 | 30 |
| Signal loss with tonic compensation | 48 | 48 | 0 | 0 | 48 |

Establishment was assessed after 30 s of settling. The activity of population 1 had to exceed that of population 2 by at least 0.5, with the largest absolute step change below one billionth at the end of settling. Among trials that established this initial state, the first exit required population 2 to exceed population 1 by at least 0.5 for 0.5 s continuously during the subsequent 120 s. The standard deviation of the Ornstein–Uhlenbeck noise was 0.075.

Tonic compensation supplied the constant drive corresponding to the lost steady ERG3 current at the intact reference voltage. The receptor signal remained absent in this condition.
